# Spatio-molecular gene expression reflects dorsal anterior cingulate cortex structure and function in the human brain

**DOI:** 10.1101/2025.07.14.664821

**Authors:** Kinnary Shah, Michael S. Totty, Svitlana V. Bach, Madeline R. Valentine, Atharv Chandra, Heena R. Divecha, Ryan A. Miller, Sang Ho Kwon, Anthony D. Ramnauth, Madhavi Tippani, Sanjana Tyagi, Joel E. Kleinman, Leonardo Collado-Torres, Shizhong Han, Thomas M. Hyde, Stephanie C. Page, Kristen R. Maynard, Stephanie C. Hicks, Keri Martinowich

**Affiliations:** Department of Biostatistics, Johns Hopkins Bloomberg School of Public Health, Baltimore, MD, USA; Lieber Institute for Brain Development, Johns Hopkins Medical Campus, Baltimore, MD, USA; Department of Psychiatry and Behavioral Sciences, Johns Hopkins School of Medicine, Baltimore, MD, USA; Solomon H. Snyder Department of Neuroscience, Johns Hopkins School of Medicine, Baltimore, MD, USA; Biochemistry, Cellular, and Molecular Biology Graduate Program, Johns Hopkins School of Medicine, Baltimore, MD, USA; Center for Computational Biology, Johns Hopkins University, Baltimore, MD, USA; Department of Genetic Medicine, Johns Hopkins School of Medicine, MD, USA; Department of Neurology, Johns Hopkins School of Medicine, Baltimore, MD, USA; Department of Biomedical Engineering, Johns Hopkins University, Baltimore, MD, USA; Malone Center for Engineering in Healthcare, Johns Hopkins University, Baltimore, MD, USA; Johns Hopkins Kavli Neuroscience Discovery Institute, Baltimore, MD, USA

**Keywords:** dorsal anterior cingulate cortex, postmortem human brain, single-nucleus RNA-sequencing, spatially-resolved transcriptomics

## Abstract

In the human brain, the dorsal anterior cingulate cortex (dACC) plays key roles in various components of cognitive control, and is particularly relevant for reward processing and conflict monitoring. The dACC regulates expression of fear and pain, and its dysfunction is implicated in a number of neuropsychiatric disorders. Compared to more recently specialized neocortical areas, such as the dorsolateral prefrontal cortex (dlPFC), the dACC is evolutionarily older. The region’s agranular structure, and other evolutionary specializations, such as the presence of von Economo neurons (VENs), contribute to its specialized roles in cognitive and emotional processing. Here, we generated paired spatially-resolved transcriptomics (SRT) and single-nucleus RNA-sequencing (snRNA-seq) data from adjacent tissue sections of the dACC in ten adult neurotypical donors to define molecular profiles for dACC cell types and spatial domains. Using non-negative matrix factorization (NMF), we integrated these data by identifying gene expression patterns within the snRNA-seq data, which were projected onto the SRT data to infer the spatial localization. Combining these data with publicly available resources, we revealed insights about molecular profiles, spatial topography, enrichment of disease risk, and putative connectivity of spatially-localized dACC cell types, including VENs. Utilizing published dlPFC snRNA-seq and SRT data collected in the same neurotypical brain donors used here, we deployed cross-region comparison analyses between dACC and dlPFC to understand spatio-molecular specializations and laminar organization across human brain evolution. To make this comprehensive molecular resource accessible to the scientific community, we made both raw and processed data freely available, including through interactive web applications.

## 1 Introduction

The dorsal anterior cingulate cortex (dACC) plays key roles in various components of cognitive control, and is particularly relevant for processes associated with response selection, conflict monitoring, and error detection (Devinsky et al. 1995; Carter et al. 1998, 2009; Heilbronner and Hayden 2016). The dACC’s role in these cognitive processes is especially pronounced in the context of reward processing. Neural firing patterns in the dACC reflect reward prediction error, which is related to its proposed function as a reward monitor (Bush et al. 2002; Hayden et al. 2011), and reward seeking is associated with changes in neural activity of dACC neurons (Seo and Lee 2007). The dACC has a notable role in sustained attention and is linked to disorders that feature attentional deficits (Carter et al. 1998; Weissman et al. 2005; Seidman et al. 2006; Bryden et al. 2011). dACC neural activity is associated with expression of pain, anxiety, and fear, and is implicated in the etiology of neuropsychiatric disorders related to these symptoms (Rainville et al. 1997; Bush et al. 2000; Milad et al. 2007; Etkin et al. 2011; Shackman et al. 2011; Lieberman and Eisenberger 2015).

The dACC is a subdivision of the larger anterior cingulate cortex (ACC), which lies in the medial wall on each cerebral hemisphere, located above and adjacent to the corpus callosum. The ACC is highly connected to both the prefrontal cortex and to subcortical regions within the limbic system, and the topography of connections to these areas differs across subdivisions of the ACC (Stevens et al. 2011; van Heukelum et al. 2020). The ACC is typically subdivided into the more ventral aspect that contains the subgenual ACC (sgACC) and pregenual ACC (pgACC) versus the more dorsal aspect, the dACC, which is also referred to as midcingulate cortex (MCC) in several nomenclature systems (2009; Stevens et al. 2011; van Heukelum et al. 2020). Anatomically, the dACC is located dorsal to the genu of the corpus callosum and encompasses parts of Brodmann areas (BA) 33, 24, and 32 (2009; Stevens et al. 2011; van Heukelum et al. 2020) (**Supplementary Fig. 1**).

Compared to more recently specialized neocortical areas, including the dorsolateral prefrontal cortex (dlPFC), the ACC is an evolutionarily older structure whose most prominent cytoarchitectural feature is the lack of a definable Layer 4 (L4) (Vogt et al. 1995; Petrides and Pandya 2002). The region’s agranular structure and other notable specializations, such as the unique presence of von Economo neurons (VENs), are hypothesized to contribute to its specialized roles in cognitive and emotional processing. VENs are a type of spindle neuron present in great apes and humans, but their presence is more controversial in other primates or mammals (Nimchinsky et al. 1999; Watson et al. 2006; Allman et al. 2011). VENs differ from canonical cortical pyramidal neurons in their size and shape, and are thought to play an adaptive function in mediating fast communication across long distances (Allman et al. 2001). Postmortem human brain studies in other cortical regions have suggested that genetic risk for various functions and neuropsychiatric disorders may manifest with cell type and laminar specificity (Skene et al. 2018; Maynard et al. 2021; Batiuk et al. 2022; Huuki-Myers et al. 2024; Wamsley et al. 2024), but whether and how these findings are similar or differ in the dACC is not known.

Here, we generated spatially-resolved transcriptomic (SRT) maps with paired single-nucleus RNA-sequencing (snRNA-seq) data to provide a comprehensive data resource of the human dACC. We highlight how this data resource can be used to better understand biological features unique to this region, and to address questions about anatomical conservation with more recently evolved cortical regions that can provide insight into the biology underlying differences in function and connectivity. dACC snRNA-seq and SRT data were generated in ten adult neurotypical donors to define molecular profiles for cell types and spatial domains. Using non-negative matrix factorization (NMF), we integrated these data by defining gene expression patterns within the snRNA-seq data and inferred their expression in the SRT data. Combining these data with publicly available resources, we reveal insights about molecular profiles, spatial topography, enrichment of disease risk, and putative connectivity of spatially-localized dACC cell types, including VENs. Utilizing dlPFC snRNA-seq and SRT data previously collected in the same ten neurotypical brain donors used to generate the dACC data (Huuki-Myers et al. 2024), we deployed cross-region comparative analyses between dACC and dlPFC to understand spatio-molecular specializations and laminar organization across human brain evolution. To make this comprehensive molecular resource accessible to the scientific community, both raw and processed data are freely available, including through interactive web applications.

## 2 Results

### 2.1 Spatio-molecular profiling of the human dorsal anterior cingulate cortex (dACC)

While the molecular organization of the human dlPFC has been well-characterized using single-cell and spatial transcriptomics (Zhu et al. 2018; Maynard et al. 2021; Emani et al. 2024; Huuki-Myers et al. 2024), other cortical regions, including the evolutionarily older ACC, are less defined. The dlPFC is classified as a neocortex, containing six prominent cortical layers with a well-defined granular L4. In contrast, the ACC is an evolutionarily older type of allocortex, classified as dysgranular or agranular cortex, which is characterized histologically by a poorly defined or non-existent granular L4 (**Fig. 1A**) (Vogt et al. 1995; Petrides and Pandya 2002). The ACC is divided into three major components - the subgenual (sgACC), pregenual (pgACC), and dorsal (dACC). The dACC (also referred to as the caudal ACC [cACC] or midcingulate [MCC] in other nomenclature systems) can be further divided into an anterior aspect (e.g. adACC, acACC, aMCC) and a posterior aspect (e.g. pdACC, pcACC, pMCC) (2009; Stevens et al. 2011; van Heukelum et al. 2020) (**Supplementary Fig. 1**). We dissected tissue blocks from fresh-frozen coronal hemislabs of postmortem human brain from ten neurotypical control donors (6 male/4 female) targeted at the anterior aspect of the dACC (**Supplementary Fig. 1**). Inclusion of dACC on the tissue blocks was confirmed by histological staining (H&E) and RNAScope using established laminar marker genes (**Fig. 1B**, **Supplementary Fig. 2**). To generate SRT data, tissue blocks were vertically scored in ∼6.5 mm strips to match the width of capture areas on the Visium platform (10x Genomics). Four independent donors were processed per Visium slide, with a total of 17 total capture areas. For a subset of donors, we generated technical replicates from directly adjacent sections (2-3 additional capture areas per donor). After standard preprocessing and quality control workflows to remove low-quality Visium spots (**Methods**, **Supplementary Fig. 3**, **Supplementary Fig. 4**, **Supplementary Fig. 5**), 73,367 spots were retained. Following tissue collection for SRT assays, 2-4 100μm cryosections from each tissue block were collected (**Fig. 1B**) for snRNA-seq on the Chromium platform (10x Genomics). Nuclei were isolated from these cryosections for each donor, and sorted on propidium iodide (PI) to sample all cell types (**Methods**). After standard preprocessing and quality control to remove empty droplets, doublets, and poor-quality nuclei, 35,161 high-quality nuclei across all 10 donors were retained (**Methods**). To facilitate comparative molecular neuroanatomy studies between the evolutionarily specialized, granular dlPFC and the evolutionarily ancient, agranular dACC, we leveraged existing dlPFC data from the same donors previously used to characterize the spatial and cellular landscape of the dlPFC (**Fig. 1A**, **Supplementary Table 1**) (Huuki-Myers et al. 2024).

**Fig. 1.**
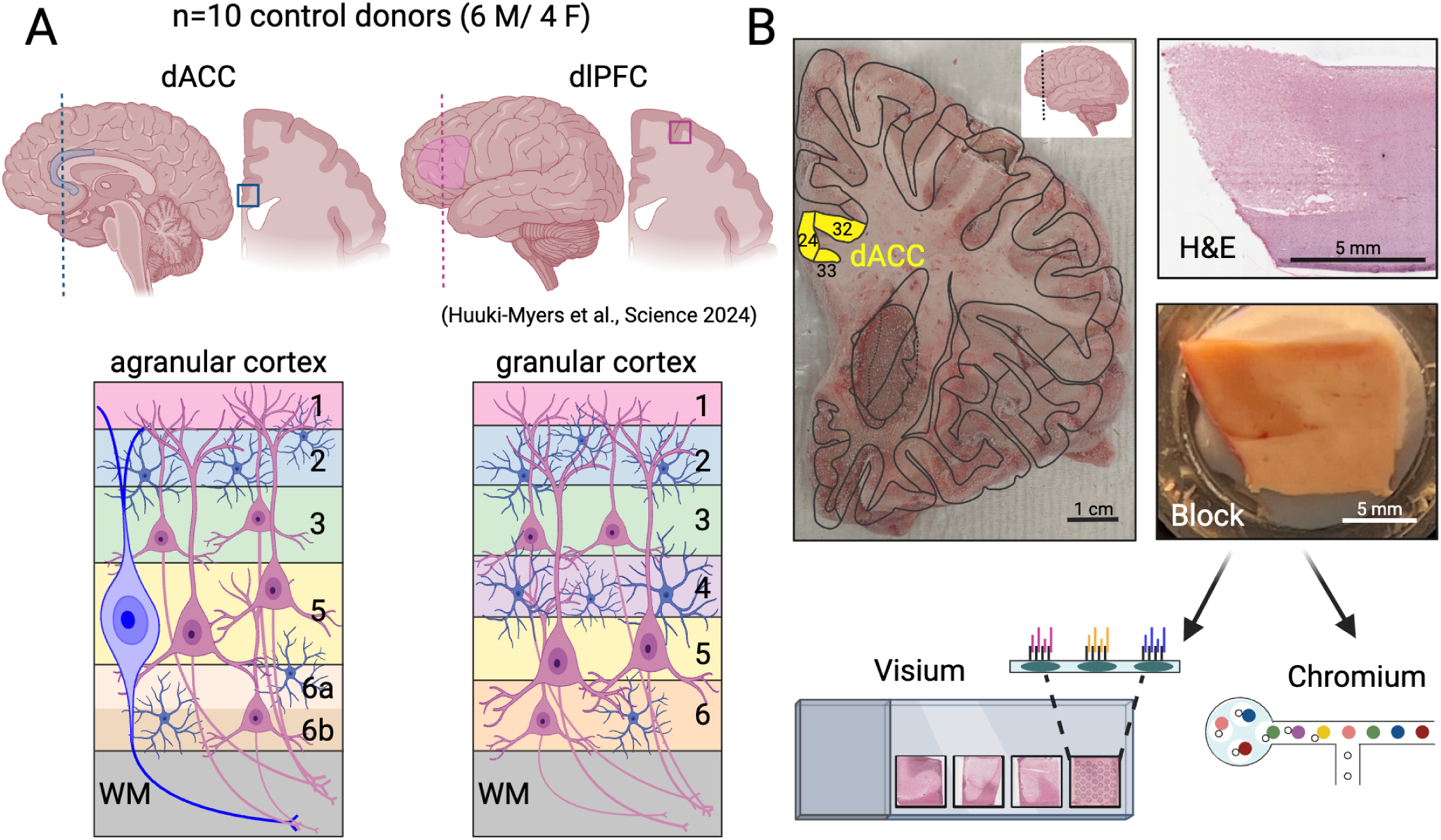
Experimental design to generate paired single-nucleus RNA-sequencing (snRNA-seq) and spatially-resolved transcriptomics (SRT) data in the human dorsal anterior cingulate cortex (dACC). (**A**) dACC (blue) and dlPFC (pink) regions outlined in midsagittal (dACC) and lateral (dlPFC) brain views, as well as in coronal hemislabs. dACC tissue (blue) used here was sourced from the same ten neurotypical control donors previously profiled for dlPFC (pink) (Huuki-Myers et al. 2024) to facilitate within-donor comparisons of dACC agranular cortex versus dlPFC granular cortex (bottom). (**B**) Fresh-frozen coronal brain slab from donor Br3942 taken at the level of the anterior striatum overlaid with the outline of major landmarks from the *Atlas of the Human Brain* (Mai et al. 2015). Three Brodmann areas (BA) 33, 24, and 32 corresponding to the dACC are highlighted in yellow (top left). H&E staining of tissue cryosections confirmed inclusion of dACC on the tissue block (top right). Following anatomical validation, cryosections from 10 neurotypical control donors were collected for Visium and Chromium assays (10x Genomics) from the same block for each donor (bottom).

### 2.2 Molecular signatures of spatial gene expression identify discrete spatial domains and reveal agranular laminar patterning in the human dACC

We used log_2_-transformed normalized gene counts as input to `PRECAST`, a spatial cluster detection algorithm, to find data-driven clusters in the dACC SRT data (Liu et al. 2023). As input to the `PRECAST` algorithm, we identified a set of spatially variable genes (SVGs) using `nnSVG` (Weber et al. 2023), as described in **Methods**. These SVGs are listed in **Supplementary Table 2**. We generated clusters at a range of *k*=5 to *k*=20 using `PRECAST` guided by the set of SVGs, and used both average cluster purity and H_+_ as statistical metrics coupled with predicted laminar organization to choose a final set of clusters for downstream analyses (**Methods**, **Supplementary Fig. 6**, **Supplementary Fig. 7**). Ultimately, we selected the clusters generated with *k*=9 (**Supplementary Fig. 8**). Independently of the `PRECAST`-derived data-driven clusters, we used visualization of histological features and expression of established layer marker genes to generate a set of histologically-driven domains, which we annotated as L1, L2/3, L5, L6, white matter (WM), and the corpus callosum (CC) (**Fig. 2A,B, Supplementary Fig. 9**). Based on visual comparison to the histology-driven domains and expression of canonical marker genes, we noted that data-driven clusters 7, 3, and 8 mapped to WM or CC in the histology-driven domains. Data-driven cluster 7 corresponded with very low library size (**Supplementary Fig. 8**), and was hence removed from downstream analysis. Data-driven cluster 8 contained a small number of spots that were speckled and contained within data-driven cluster 3; this larger cluster matched the known cytoarchitecture and localization of WM beneath L6. Furthermore, data-driven clusters 3 and 8 did not differentiate between WM and CC, as seen in the manual annotations of the dACC SRT data (**Supplementary Fig. 9**, **Supplementary Fig. 10**). Thus, we retained data-driven clusters 3 and 8, but collapsed them into a single WM spatial cluster, resulting in a final set of seven data-driven spatial domains (**Fig. 2C**). Each of these spatial domains contained a comparable number of spots from each of the ten donors, with no individual donor dominating the composition of any spatial domain (**Supplementary Fig. 11**).

**Fig. 2.**
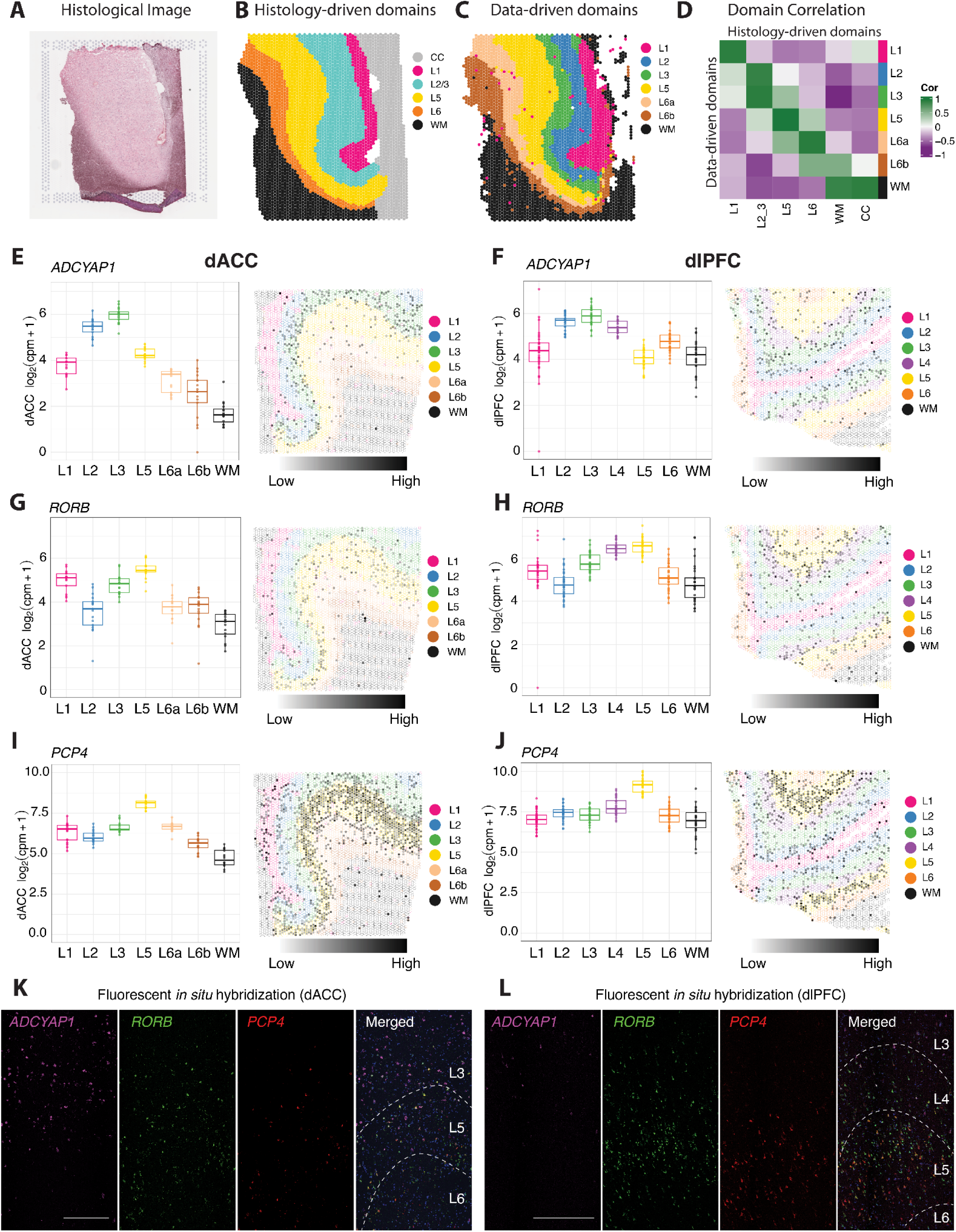
Signatures of gene expression in data-driven spatial domains provide molecular evidence for agranular organization of the dACC. (**A**) Histological image of Visium capture area from donor Br8667 (sample ID: V12Y31-080_B1). (**B**) Spot plot of Visium capture area from donor Br8667 (sample ID: V12Y31-080_B1) with spots colored by histology-driven domains, which were annotated as Layer 1 (L1), L2/3, L5, L6, white matter (WM), and corpus callosum (CC). (**C**) Spot plot of Visium capture area from donor Br8667 (sample ID: V12Y31-080_B1) with spots colored by the 7 spatial domains annotated from the *k*=9 `PRECAST` clusters, which were annotated as L1, L2, L3, L5, L6a, L6b, and WM. (**D**) `spatialLIBD` spatial registration heatmap displays Pearson’s correlation values between the top 100 marker genes in each data-driven spatial domain (*y*-axis) and in each histology-driven manually annotated spatial domain (*x*-axis). (**E**) Boxplots of *ADCYAP1* log_2_(counts per million + 1) expression (computed manually) (*y*-axis) for each spatial domain (*x*-axis) in the pseudobulked dACC SRT data. Color represents the spatial domain. `escheR` spot plot of dACC Visium capture area from donor Br6432 (sample ID: V12N28-331_B1) with spots colored by the dACC spatial domains. Fill represents *ADCYAP1* log_2_-normalized expression per spot. (**F**) Boxplots of *ADCYAP1* log_2_(counts per million + 1) expression (computed manually) (*y*-axis) for each spatial domain (*x*-axis) in the pseudobulked dlPFC SRT data. Color represents the spatial domain. Spot plot of dlPFC Visium capture area from donor Br6432 (sample ID: Br6432_ant) with spots colored by the dlPFC spatial domains. Fill represents *ADCYAP1* log_2_-normalized expression per spot. (**G**) Same as (**E**), but for *RORB* expression. (**H**) Same as (**F**), but for *RORB* expression. (**I**) Same as (**E**), but for *PCP4* expression. (**J**) Same as (**F**), but for *PCP4* expression. (**K**) Multiplex RNAScope single molecule fluorescence *in situ* hybridization (smFISH) in dACC of donor Br6432 for *ADCYAP1* (magenta)*, RORB* (green)*, PCP4* (red), and merged all channels with DAPI (blue). Approximate cortical layer boundaries are indicated by dashed lines. Scale bar 500 µm. (**L**) Same as (**K**), but for dlPFC.

To better understand the relationship between the seven data-driven spatial domains to canonical histological layers, we used a combination of approaches. We first pseudobulked spots within the data-driven and histology-driven domains, which collapses data from spot-level to spatial domain-level data within each sample. In contrast to sparse spot-level information, this information is similar to bulk RNA-sequencing for each domain, enabling the use of differential expression (DE) tools optimized for bulk data. Prior to proceeding with DE analysis, we examined the samples in principal component (PC) space to ensure that the data-driven domains were relatively well-separated, and demonstrated that any variation was not driven by total UMI counts per sample (**Supplementary Fig. 12**). For these analyses, we pseudobulked samples by aggregating total UMI counts across spatial domains and samples. Comparison of the pseudobulked data-driven domains to the histology-driven annotations revealed high correlations across layers and WM (**Fig. 2D**).

Next, we capitalized on the availability of an SRT dataset of the dlPFC (Huuki-Myers et al. 2024) generated from the same 10 brain donors used in this study to directly investigate the molecular organization of granular versus agranular cortex. Specifically, we visualized expression of established cortical layer markers derived in the dlPFC data, including *RELN* (L1)*, LAMP5* (L2)*, ADCYAP1* (L3), *RORB* (L4), *PCP4* (L5), *NR4A2* (L6), and *MOBP* (WM) across the data-driven domains in both the dACC and dlPFC (**Fig. 2E-J, Supplementary Fig. 13**). The data-driven domains in the dlPFC include domains for WM and the 6 layers typical of this granular structure (**Fig. 2F,H,J**, **Supplementary Fig. 13**). We also performed multiplex single molecule fluorescence in situ hybridization (smFISH) for three of these markers, *ADCYAP1*, *RORB*, and *PCP4*, in dACC and dlPFC tissue sections from the same donor (**Fig. 2K,L**). Based on the comparison to histology-driven domains (**Fig. 2D**) and enrichment of layer marker genes (**Fig. 2E-J, Supplementary Fig. 13**), we identified single dACC data-driven domains that distinctly map to L1, L2, L3, L5, and WM. We noted that two of the dACC data-driven domains showed high correlation to the histology-driven domain annotated as L6 (**Fig. 2D**). These two domains expressed canonical markers of L6, including *NR4A2* (**Supplementary Fig. 13I-L**). We therefore annotated these two dACC data-driven domains as L6a and L6b. In the dlPFC SRT data, expression of the canonical L4 marker *RORB* is high in both L4 and L5 (**Fig. 2H**), while *RORB* maps tightly to L5 in the dACC SRT data (**Fig. 2**). Confirming these results, smFISH reveals that *RORB* expression was co-localized with the L5 marker *PCP4* in the dlPFC, but also extended superficially into L4, which contains no *PCP4* expression (**Fig. 2L**). However, in the dACC, *RORB* expression is limited to the L5 area defined by *PCP4* expression (**Fig. 2K**). In summary, we generated molecular profiles for dACC data-driven domains using spatial gene expression data at transcriptome-scale, which we annotated as L1, L2, L3, L5, L6a, L6b, and WM. While the agranular structure of the ACC has been long appreciated from a cytoarchitecture perspective, these data provide complementary molecular evidence for the absence of L4 in dACC in the human brain.

### 2.3 Non-negative matrix factorization (NMF) reveals cell type-specific gene expression patterns shared between snRNA-seq data in the dACC

To better characterize the molecular identity of the data-driven spatial domains in the dACC, we integrated the SRT data with paired snRNA-seq data, which was generated from adjacent tissue sections in each donor. Following quality control of the snRNA-seq data (**Supplementary Fig. 14**, **Supplementary Fig. 15**, **Supplementary Fig. 16**), we used the `Azimuth` Human Motor Cortex reference to predict and transfer cell type annotations to our dataset (Bakken et al. 2021; Hao et al. 2024). Each cell type contained a comparable proportion of nuclei from each of the ten donors with no individual donor dominating the composition of any cell type (**Supplementary Fig. 17**). To integrate the snRNA-seq across donors, we selected the top 2,000 features based on Poisson deviance (Townes et al. 2019) (**Supplementary Fig. 18**) followed by `Harmony` batch correction (Korsunsky et al. 2019) on the GLM-PCA reduced dimensions. We found comparable numbers of nuclei in each cell type with good separation across cell types (**Fig. 3A**). Similar to the SRT data, we used pseudobulked DE analysis to find marker genes for each cell type compared to all other cell types. The cell types were well-separated in principal components (PC) space and this variation was not driven by total UMI counts per sample (**Supplementary Fig. 19**). We generated volcano plots for each cell type (**Supplementary Fig. 20**, **Supplementary Fig. 21**) and extracted the top 30 markers (**Supplementary Table 3**). To annotate cell types from the snRNA-seq data with the spatial domains identified using the SRT data, we used the `spatialLIBD` spatial registration framework of computing the correlation of *t*-statistics of the top 100 markers for each combination of cell type and spatial domain (Maynard et al. 2021; Pardo et al. 2022; Huuki-Myers et al. 2024). Layer-specific cell types, such as L2/3 intratelencephalon (IT)-projecting neurons, L5 IT, L6 corticothalamic (CT)-projecting neurons, and L6b, were enriched for their respective spatial domains, while astrocytes were enriched in L1 and oligodendrocytes in WM (**Fig. 3C**). Several layer-enriched marker genes in the SRT data were also cell type marker genes in the snRNA-seq data, including *ADCYAP1*, *RORB*, and *PCP4* (**Fig. 2E-J**). Strong expression of these genes within their respective layer-specific cell types further confirms laminar identity assignment for the dACC data-driven spatial domains (**Fig. 3B**). Comparing the dlPFC and dACC snRNA-seq data, for the dlPFC, expression of *RORB* is highest in the L4 cell type, but also expressed in the L5 cell type, while in the dACC, *RORB* expression is localized more specifically to the L5 cell type (**Supplementary Fig. 22**). These data are concordant with the SRT data (**Fig. 2G, H**), and provide additional molecular evidence at the cellular level of agranularity in the dACC.

**Fig. 3.**
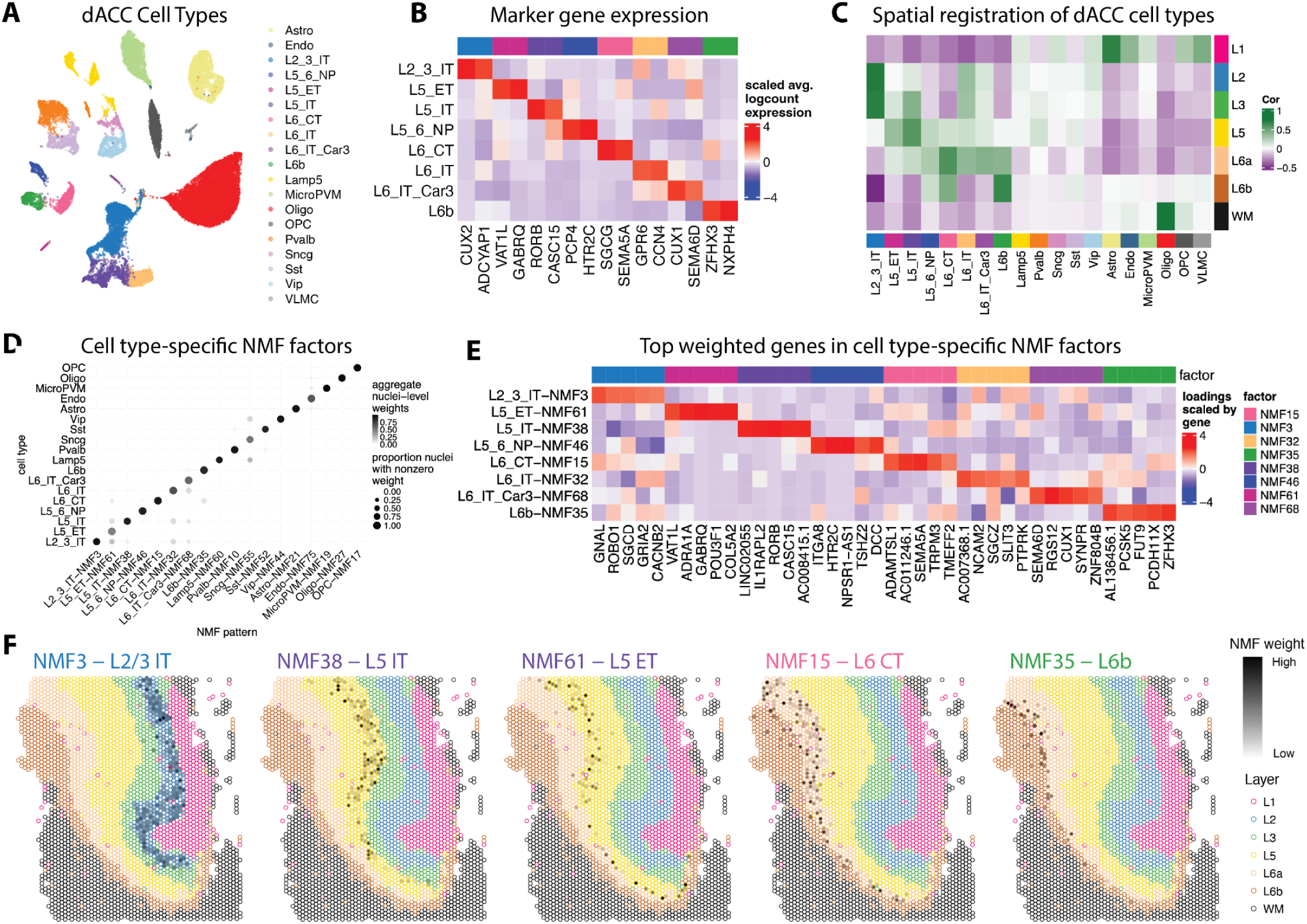
Single-nucleus RNA-sequencing (snRNA-seq) cell type annotation and non-negative matrix factorization (NMF) reveals distinct cell types in the human dACC. (**A**) Uniform manifold approximation and projection (UMAP) representation of the snRNA-seq dataset. Each point is an individual nucleus colored by its `Azimuth` reference cell type. Astro: Astrocyte; Endo: Endothelial cell; L2_3_IT: Layer 2-3 glutamatergic neuron, intratelencephalon-projecting; L5_ET: Layer 5 glutamatergic neuron, extratelencephalon-projecting; L5 IT: Layer 5 glutamatergic neuron, intratelencephalon-projecting; L5_6_NP: Layer 5-6 glutamatergic neuron, near-projecting; L6_CT: Layer 6 glutamatergic neuron, corticothalamic-projecting; L6_IT: Layer 6 glutamatergic neuron, intratelencephalon-projecting; L6_IT_Car3: Layer 6 Car3+ glutamatergic neuron, intratelencephalon-projecting; L6b: Layer 6b glutamatergic neuron; Lamp5: Lamp5+ GABAergic neuron; MicroPVM: microglia / perivascular macrophage; Oligo: oligodendrocyte; OPC: oligodendrocyte precursor cell; Pvalb: Pvalb+ GABAergic neuron; Sncg: Sncg+ GABAergic neuron; Sst: Sst+ GABAergic neuron; Vip: Vip+ GABAergic neuron; VLMC: Vip+ GABAergic neuron. (**B**) Heatmap showing two selected marker gene log_2_-normalized expression averaged by cell type and scaled per gene (columns) across the layer-specific cell types (rows): L2/3 IT, L5 ET, L5 IT, L5/6 NP, L6 CT, L6 IT, L6 IT Car3, and L6b. Grouping and color across the top indicates the cell types. (**C**) `spatialLIBD` spatial registration heatmap displays Pearson’s correlation values between the top 100 marker genes in each SRT data-driven spatial domain and in each snRNA-seq cell type. The *x*-axis displays the snRNA-seq cell types described in (**A**). The *y*-axis displays the data-driven domains: L1, L2, L3, L5, L6a, L6b, and white matter (WM). (**D**) Dot plot showing the top NMF pattern (*x*-axis) selected for each snRNA-seq cell type (*y*-axis). Dot size represents the proportion of nuclei with nonzero weight for each NMF pattern across each cell type. Dot color represents the aggregated nuclei-level weights for each NMF pattern across each cell type. (**E**) Heatmap showing 5 selected marker gene loadings scaled by total gene loadings (columns) across the layer-associated NMF patterns (rows): L2/3 IT, L5 ET, L5 IT, L5/6 NP, L6 CT, L6 IT, L6 IT Car3, and L6b. Grouping across the top indicates the NMF patterns and the color represents the cell type each pattern is associated with. (**F**) `escheR` spot plots of dACC Visium capture area from donor Br8667 (sample ID: V12Y31-080_B1) with spots colored by the dACC spatial domains. Grey scale fill represents weights of distinct cell type-specific NMF gene expression patterns learned from the snRNA-seq dataset and projected into the SRT data using transfer learning.

Next, we used non-negative matrix factorization (NMF) as a complementary approach to identify cell type-specific gene expression programs (Thompson et al. 2024). NMF reduces the dimensions of the gene expression data to *k* factors that represent cell types, cell patterns, and various biological processes (Stein-O’Brien et al. 2019; DeBruine et al. 2021). Using cross-validation, we determined that the optimal rank to factorize the snRNA-seq gene expression matrix was 75 factors (**Supplementary Fig. 23**). After factorizing the gene expression matrix, we first annotated these 75 patterns based on association with specific cell types (**Methods**, **Supplementary Fig. 24**, **Supplementary Table 4**). We removed NMF patterns associated with technical variables by computing the correlation with each pattern’s weight and variables related to quality control metrics, brain donor, and sex of brain donor and removed them from further downstream analyses (**Supplementary Fig. 25**). The dot plot identifies a top NMF pattern for each cell type (the vascular leptomeningeal cell (VLMC) cell type did not have a specific NMF pattern), and demonstrates that each pattern is highly specific to the cell type to which it was annotated (**Fig. 3D**). For the layer-specific cell types, we extracted the top marker genes for each pattern, highlighting genes whose function has been associated with function in these layers (**Fig. 3E**). Finally, we projected these NMF patterns into the SRT data using a transfer learning approach in order to predict the spatial location of distinct neuronal subtypes (**Fig. 3F**). We found that these cell type-specific NMF patterns correspond to the laminar organization of the dACC, with patterns enriched in specific neuronal cell types aligning to their expected cortical layer (e.g., L5 NMF patterns localizing to the L5). These findings provide a foundation for exploring biologically meaningful, layer-specific gene expression programs and mapping them into SRT data.

### 2.4 Localization of a signature for von Economo neurons (VENs) to a spatially-restricted area of deep L5 in dACC

A highly unique feature of the dACC is the presence of VENs. VENs are a specialized type of bipolar spindle neuron that is localized to L5, but only in a restricted set of human brain regions including the pgACC, dACC, and frontoinsula. VENs are large in size, have a unique morphology, and are more abundant in the human brain compared to nonhuman primates, possibly representing an evolutionary specialization (Nimchinsky et al. 1999; Watson et al. 2006; Allman et al. 2011). We utilized a published dlPFC snRNA-seq dataset generated in the same 10 donors as the dACC snRNA-seq dataset (Huuki-Myers et al. 2024), to explore molecular patterns that might map specifically to region-specific cell types, including VENs. Similar to previous analyses, we computed pseudobulked DE genes by comparing each cell type to all other cell types, first visualizing the pseudobulked data along the first two principal components (**Supplementary Fig. 26**), and then calculating the correlation between the top 100 markers for each dACC cell type and each dlPFC cell type (**Supplementary Fig. 27**). Every dACC cell type was strongly correlated with at least one dlPFC cell type, except for L5 extratelencephalon-projecting neurons (ET) and L6 IT Car3, a subpopulation of L6 neurons marked by expression of *CAR3 (Peng et al. 2021)* (**Supplementary Fig. 27**).

We further explored the L5 ET cell type, hypothesizing that it may contain VENs, which specifically reside in L5 of dACC, but not in dlPFC (**Fig. 4A**). We identified both NMF38 and NMF61 as patterns from the dACC snRNA-seq data specifically associated with L5; however, we found that NMF38 was highly specific to L5 IT, while NMF61 was specific to L5 ET (**Fig. 3D**). Further, we showed that several VEN markers: *VAT1L*, *ADRA1A*, *GABRQ*, and *POU3F1*, were specifically associated with L5 ET/NMF61 (**Fig. 3E**). We extended these findings to the SRT data by using transfer learning to project the NMF38 and NMF61 patterns into both the dACC SRT data, as well as the dlPFC SRT data, generated from the same paired ten donors (Huuki-Myers et al. 2024) (**Fig. 4B,C**). The output that we obtained is spot-level weights for these two patterns in each dataset. First, in the dACC SRT data, we determined that the spots associated with NMF38 and NMF61 were distinct for all samples (**Supplementary Fig. 28**). While both NMF patterns localized to L5, they had a distinct topography within the layer. Specifically, NMF61 is localized to deep L5 while NMF38 is localized to superficial L5 (**Fig. 4D**). Comparing directly across the dACC and dlPFC SRT datasets, we showed that NMF38 is expressed relatively similarly across L5 in both the dACC and the dlPFC, while NMF61 is highly expressed only in L5 of the dACC (**Fig. 4D**). Quantifying these observations, we demonstrated that the fraction of nonzero L5 spots across donors is similar between the dACC and dlPFC for NMF38, but higher in the dACC for NMF61. To make an analogous comparison with snRNA-seq data, we also mathematically projected these two NMF patterns into the dlPFC snRNA-seq data to predict nuclei-level weights for each pattern. In the snRNA-seq data, we computed sample averages of each NMF pattern’s weights from nuclei in the cell type of interest, using cell type L5 IT or L5 ET for dACC and cell type Excitatory L5 in dlPFC. We found that NMF38 was more highly expressed in the dACC L5 IT cell type compared to the dlPFC Excitatory L5 cell type, and that NMF61 was more highly expressed in the dACC L5 ET cell type compared to the dlPFC Excitatory L5 cell type. These data demonstrate that the average NMF pattern weight is higher in the dACC snRNA-seq data compared to the dlPFC snRNA-seq data for both NMF38 and NMF61 (**Fig. 4E**). We calculated differentially expressed genes between the dACC L5 IT and L5 ET cell types in the snRNA-seq data, which revealed that several established VEN markers (Yang et al. 2019; Hodge et al. 2020), specifically *HAPLN4, SULF2*, *FEZF2*, and *GABRQ*, are more highly expressed in L5 ET compared to L5 IT (**Fig. 4F**). We also calculated differentially expressed genes between the NMF61-positive and NMF38-positive spots in the dACC SRT data (**Methods**), which similarly showed that several established VEN markers (Yang et al. 2019; Hodge et al. 2020), specifically *VAT1L*, *SULF2*, *FEZF2*, and *GABRQ*, are more highly expressed in NMF61 compared to NMF38 (**Fig. 4F**).

**Fig. 4.**
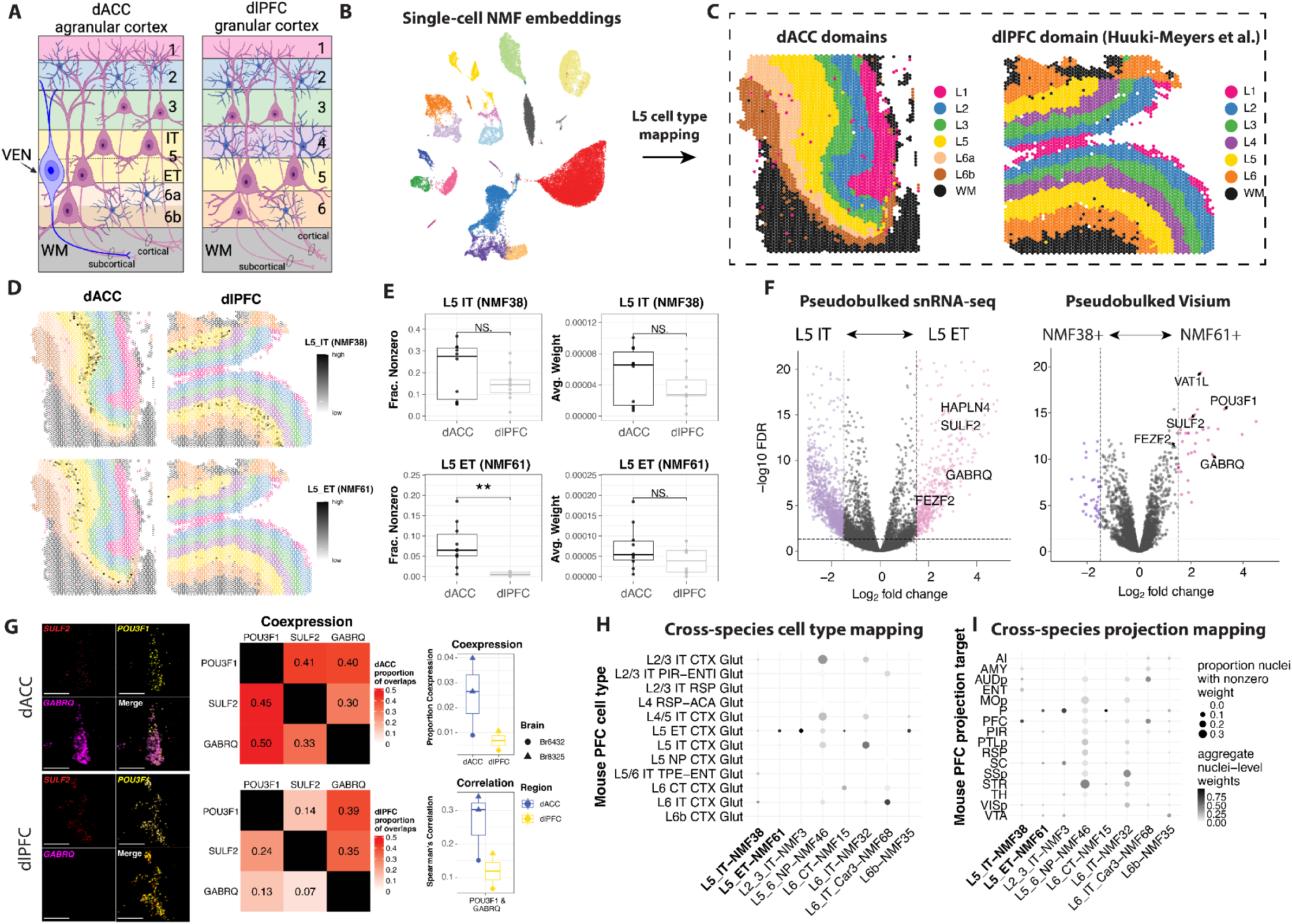
Identification of discrete molecular signatures for von Economo neurons (VENs) in Layer 5 of the dACC. (**A**) Cytoarchitecture of dACC agranular cortex and dlPFC granular cortex at laminar resolution highlighting the relative location of VENs in Layer 5 ET. This cell type was captured by NMF61. (**B**) UMAP plot from Fig. 3A shows cell types from the dACC snRNA-seq data. The L5 NMF patterns derived from these data are projected into the SRT data shown in Fig. 4C. (**C**) First spot plot is the same as Fig. 2C. The second spot plot shows Visium capture area from dlPFC donor Br8667 (sample ID: Br8667_mid) with spots colored by the 7 spatial domains annotated from the previous dlPFC study (Huuki-Myers et al. 2024). (**D**) TOP ROW: `escheR` spot plot of dACC Visium capture area from donor Br8667 (sample ID: V12Y31-080_B1) with spots colored by the dACC spatial domains. Spot plot of dlPFC Visium capture area from donor Br8667 (sample ID: Br8667_mid) with spots colored by the dlPFC spatial domains. Fill represents NMF38 projected into the dlPFC and dACC SRT datasets. BOTTOM ROW: Spot plot of dACC Visium capture area from donor Br8667 (sample ID: V12Y31-080_B1) with spots colored by the dACC spatial domains. Spot plot of dlPFC Visium capture area from donor Br8667 (sample ID: Br8667_mid) with spots colored by the dlPFC spatial domains. Fill represents NMF61 projected into the dlPFC and dACC SRT datasets. (**E**) FIRST COLUMN: Boxplots comparing NMF38 and NMF61 between dACC and dlPFC Layer 5 spatial domain. The top boxplots are made up of sample averages of the fraction of nonzero NMF38 spots in the dACC SRT Layer 5 and the dlPFC SRT Layer 5. The bottom boxplots are made up of sample averages of the fraction of nonzero NMF61 spots in the dACC SRT Layer 5 and the dlPFC SRT Layer 5. SECOND COLUMN: Boxplots comparing NMF38 and NMF61 between dACC and dlPFC Layer 5 cell types. The top boxplots show sample averages of NMF38 weights within the dACC snRNA-seq L5 IT cell type and the dlPFC snRNA-seq Excitatory L5 cell type. The bottom boxplots show sample averages of NMF61 weights within the dACC snRNA-seq L5 ET cell type and the dlPFC snRNA-seq Excitatory L5 cell type. (**F**) Left: Differential expression volcano plot compares pseudobulked snRNA-seq cell types L5 IT and L5 ET. Right: Differential expression volcano plot compares pseudobulked spots with nonzero NMF38 and NMF61 projections in the dACC SRT data. Known VEN markers (*VAT1L*, *SULF2*, *HAPLN4*, *FEZF2*, and *GABRQ*) are highlighted in both volcano plots as upregulated in L5 ET or NMF61. (**G**) Multiplex RNAScope single molecule fluorescence *in situ* hybridization (smFISH) in dACC (top box) and dlPFC (bottom box) for *SULF2, POU3F1, GABRQ,* and merged of all. Heatmaps: Top heatmap displays the average proportion of cells called as expressing one gene (row label) that are also called as expressing another gene (column label), averaged across the three dACC samples. Bottom heatmap is similar, except averaged across the two dlPFC samples. Boxplots: Top boxplots display the proportion of cells (*y*-axis) from Layer 5 that were called as expressing both *POU3F1* and *GABRQ* (*x*-axis). Bottom boxplots display Spearman’s correlation values (*y*-axis) for *POU3F1* and *GABRQ* expression (*x*-axis). Color indicates the region of the sample, either dACC or dlPFC. Shape indicates the brain donor of the sample, either Br6432 or Br8325. Scale bar 30 µm. (**H**) Dot plot of cortical layer-specific NMF patterns (columns) in the dACC human snRNA-seq dataset corresponding to the nuclei collection source (rows) for the mouse retroviral tracing dataset. Dot size represents the proportion of nuclei with nonzero weight for each NMF pattern across each cell type. Dot color represents the aggregated nuclei-level weights for each NMF pattern across each cell type. (**I**) Dot plot of mouse retroviral tracing using projection of cortical layer-specific NMF patterns (columns). Rows represent axonal projection target regions from the retroviral tracing experiments: AI: agranular insular cortex, AMY: amygdala, AUDp: primary auditory cortex, ENT: entorhinal cortex, MOp: primary motor cortex, P: pons, PFC: prefrontal cortex, PIR: piriform area, PTLp: posterior parietal cortex, RSP: retrosplenial cortex, SC: superior colliculus, SSp: primary somatosensory cortex, STR: striatum, TH: thalamus, VISp: primary visual cortex, and VTA: ventral tegmental area. Dot size represents the proportion of nuclei with nonzero weight for each NMF pattern across each cell type. Dot color represents the aggregated nuclei-level weights for each NMF pattern across each cell type.

As orthogonal validation, we used RNAScope smFISH to visualize 3 VEN marker genes, *POU3F1*, *SULF2*, and *GABRQ*, with the L5 marker gene *PCP4* from 5 tissue sections across 2 brain donors. Two sections were from dlPFC (Br6432 and Br8325) and 3 sections were from dACC (one from Br6432 and two from Br8325). We used *PCP4* copy count to subset to only L5 cells (**Methods**) and then scaled each sample’s copy counts by cell area. Due to fluorescence saturation, copy counts tend to be unreliable estimates, especially in the upper distribution of copy counts, so we used a Gaussian mixture model to call each cell as expressing or not expressing each gene within each sample. The proportion of cells for each sample classified as expressing *POU3F1*, *SULF2*, and *GABRQ* is presented in **Supplementary Fig. 29A**. The dACC samples had higher coexpression of 2-3 of the VEN markers compared to the dlPFC samples, with the greatest difference observed in *POU3F1* and *SULF2* coexpression (**Supplementary Fig. 29B**, **Fig. 4G**). Additionally, dACC samples had higher Spearman’s correlation between each pair of VEN markers compared to the dlPFC samples (**Supplementary Fig. 29C,D**). To further characterize coexpression, we calculated the average proportion of expressors for one gene that are also expressors for another gene, which shows higher co-expression for dACC samples compared to dlPFC samples, with greater differences for *POU3F1* and *SULF2* co-expression with the remaining two VEN markers (**Fig. 4G**). The RNAScope images show a high degree of overlap of the 3 VEN markers in the dACC, but not in the dlPFC (**Fig. 4G**).

While VENs are not present in the rodent cortex, they do show molecular homology to an ET excitatory neuron type in the mouse cortex (Hodge et al. 2020). Aligning cell types across species based on gene expression profiles can facilitate predictions about functional properties and circuit connectivity of cell types in the human brain. This information is important since these properties cannot be directly studied in postmortem human brain tissue. To extend our findings across species, we integrated our data with a mouse dataset that combined viral retrograde labeling with molecular profiling to define gene expression profiles for cells based on their axonal projection targets (Zhou et al. 2023). We first showed that NMF38 and NMF61 map to cell types defined as L5 IT and L5 ET (according to the Allen Brain subclassification), respectively, in the mouse cortex (**Fig. 4H**). We next showed that these two NMF patterns map to different cell types based on their long-range projection target - including pons, superior colliculus, thalamus, and ventral tegmental area for NMF61/L5 ET and primary auditory cortex, posterior parietal cortex, primary somatosensory cortex, and primary visual cortex in NMF38/L5 IT (**Fig. 4I**). These projection targets are in line with the localization of NMF38 to the superficial aspect of L5 versus the deep aspect of L5 for NMF61 (**Fig. 4D**). Given that these two NMF patterns find highly similar molecular features between human and mouse, particularly for L5 IT and ET neurons, these data facilitate predictions about putative axonal targets of spatially-localized L5 dACC neurons, including VENs.

### 2.5 Identification of unique spatio-molecular features in the human dACC

We first looked at molecular features across the dACC data-driven spatial domains to understand spatio-molecular features that define its laminar organization. To do this, we identified top marker genes for each of the dACC data-driven domains using the pseudobulked counts matrix generated in **Section 2.1**. We then used an enrichment model to identify genes that are differentially expressed between one data-driven spatial domain compared to all other spatial domains by fitting a linear mixed effects model on the pseudobulked counts matrix. We visualized volcano plots for each spatial domain (**Supplementary Fig. 30A-G**) and extracted the top 50 markers (**Supplementary Table 5**). We identified several genes that were enriched in dACC spatial domains that were not previously established as laminar marker genes in the dlPFC including *RSPO2* (L2), *DRD5* (L5), and *KCTD8, ADRA2A*, and *CRHBP* (L6). Visualization of these genes across the dACC and dlPFC spatial domains showed that these genes are enriched in specific dACC layers, but not in the corresponding dlPFC layers (**Fig. 5E, Supplementary Fig. 31)**. More specifically, for the L6 markers, *KCTD8* localizes to L6a while *ADRA2A* and *CRHBP* localize to L6b (**Fig. 5E, Supplementary Fig. 31**).

**Fig. 5.**
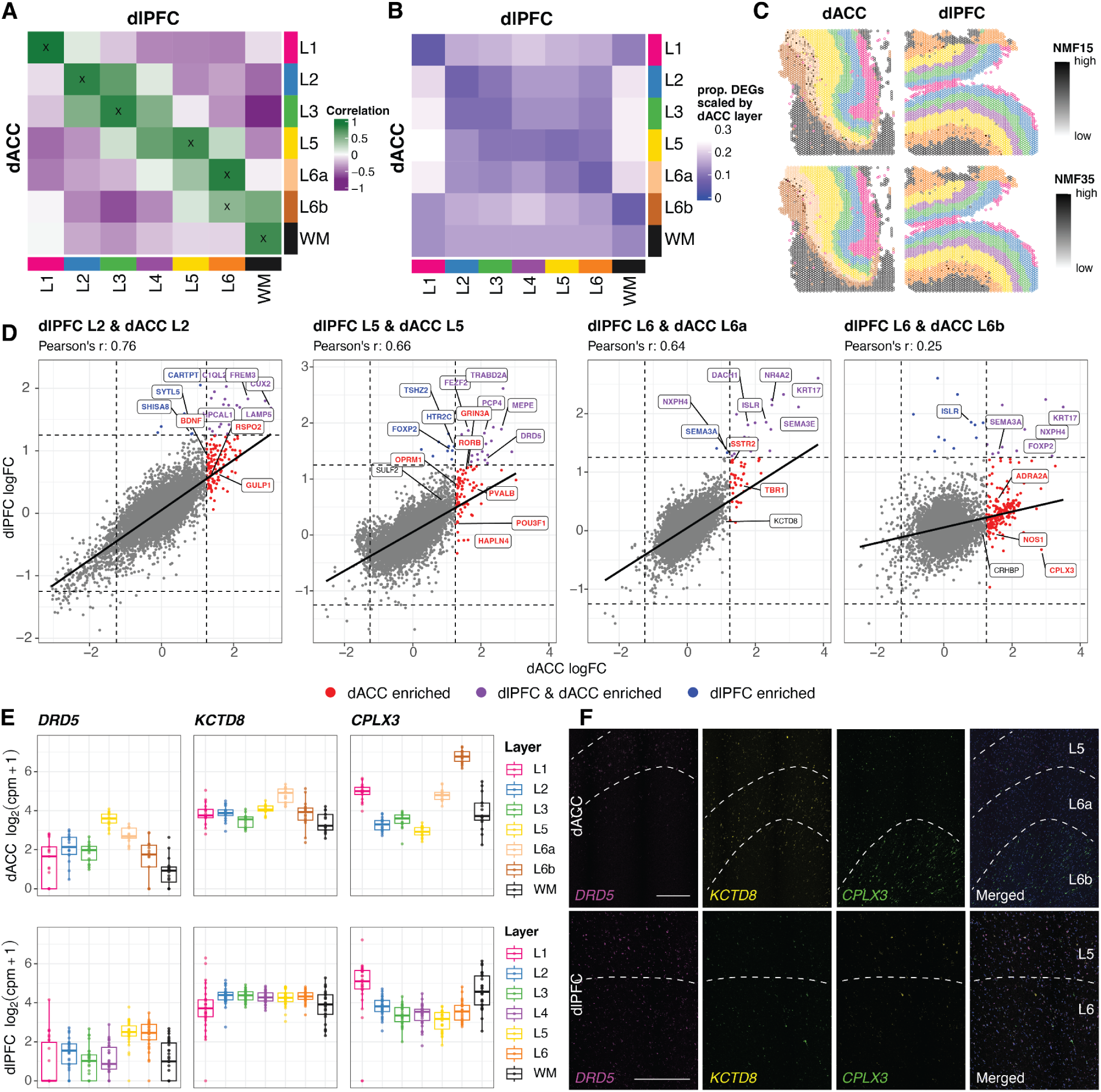
Within-donor differences in gene expression patterns across individual spatial domains of the dACC and dlPFC. (**A**) Pseudobulked sample-level comparison: `spatialLIBD` spatial registration heatmap displays Pearson’s correlation values between the top 100 marker genes in each dlPFC SRT spatial domain and in each dACC SRT spatial domain. The *x*-axis displays dlPFC spatial domains: L1, L2, L3, L4, L5, L6, and white matter (WM). The *y*-axis displays the dACC spatial domains described in (Fig. 2C). The black “X” represents high confidence. (**B**) Pseudobulked gene-level comparison: the heatmap displays the number of DEGs between each spatial domain in dACC and dlPFC, scaled by the proportion of DEGs across each dACC spatial domain. The *x*-axis displays dlPFC spatial domains: L1, L2, L3, L4, L 5, L6, and WM. The *y*-axis displays the dACC spatial domains described in (Fig. 2C). (**C**) TOP ROW: Spot plot of dACC Visium capture area from donor Br8667 (sample ID: V12Y31-080_B1) with spots colored by the dACC spatial domains. Spot plot of dlPFC Visium capture area from donor Br8667 (sample ID: Br8667_mid) with spots colored by the dlPFC spatial domains. Fill represents NMF15 projected into the dlPFC and dACC SRT datasets. BOTTOM ROW: Spot plot of dACC Visium capture area from donor Br8667 (sample ID: V12Y31-080_B1) with spots colored by the dACC spatial domains. Spot plot of dlPFC Visium capture area from donor Br8667 (sample ID: Br8667_mid) with spots colored by the dlPFC spatial domains. Fill represents NMF35 projected into the dlPFC and dACC SRT datasets. (**D**) Each scatterplot shows a comparison of the log fold-change (logFC) between one spatial domain in dACC and dlPFC SRT spatial domains. Each point is a gene. The *x*-axis of the first plot shows the logFC from enrichment model pseudobulked DE testing comparing L2 to all other spatial domains for dACC. The *y*-axis of the first plot shows the logFC from enrichment model pseudobulked DE testing comparing L2 to all other spatial domains for dlPFC. The second plot compares L5 in the dACC to L5 in the dlPFC. The third plot compares L6a in the dACC to L6 in the dlPFC. The fourth plot compares L6b in the dACC to L6 in the dlPFC. The colors highlight genes that are classified as either dACC enriched (red), dlPFC enriched (blue), or both (purple), where the threshold is a logFC greater than 1.25. The number of genes in each category is as follows: L2 dACC vs. L2 dlPFC: 117 dACC enriched and 24 dlPFC and dACC enriched; L5 dACC vs. L5 dlPFC: 80 dACC enriched, 9 dlPFC enriched, 18 dlPFC and dACC enriched; L6a dACC vs. L6 dlPFC: 42 dACC enriched, 2 dlPFC enriched, 19 dlPFC and dACC enriched; L6b dACC vs. Lb dlPFC: 191 dACC enriched, 11 dlPFC enriched, 10 dlPFC and dACC enriched. (**E**) Boxplots comparing gene expression of *DRD5, KCTD8,* and *CPLX3* across pseudobulked spatial domains from dACC SRT data and dlPFC SRT data. The *y*-axis displays log_2_(counts per million + 1) expression (computed manually) for each spatial domain (*x*-axis), for dACC (top row) and dlPFC (bottom row). Color represents spatial domains for each dataset. (**F**) Multiplex RNAScope single molecule fluorescence *in situ* hybridization (smFISH) in dACC (top row) and dlPFC (bottom row) for *DRD5* (magenta)*, KCTD8* (yellow)*, CPLX3* (green), and merged of all channels with DAPI (blue). Approximate cortical layer boundaries are indicated by dashed lines. Scale bars 500 µm.

Next, we deployed two separate approaches to directly compare spatial and cell type-specific gene expression across dACC and dlPFC. We refer to the first method of comparison as the pseudobulked sample-level comparison. For the pseudobulked sample-level comparison, we computed the correlation between the top 100 marker genes using an enrichment model across all pairs of spatial domains between dACC and dlPFC (**Fig. 5A**). This analysis confirmed strong correlation between L1, L2, L3, L5, and WM domains between dACC and dlPFC. Assessing correlation to the dlPFC L4 spatial domain, we noted that it was moderately associated with dACC L3 and L5. dlPFC L6 was more highly correlated with dACC L6a compared to L6b. We refer to the second method of comparison as the pseudobulked gene-level comparison, which was designed to directly leverage the fact that we have data from both regions from individual donors. Specifically, we used a mixed-effect model (**Methods**) that uses a random effect to take into account samples that originated from the same donor. For the pseudobulked gene-level comparison, we calculated the total number of differentially expressed genes between each pair of spatial domains in the dACC and dlPFC, scaled by the total number of DEGs in each dACC spatial domain (**Fig. 5B**). The pseudobulked gene-level analysis showed a small proportion of DEGs between L1, L2, L3, L5, and WM domains between dACC and dlPFC, indicating a strong similarity between these domains. We found complementary results between the two levels of comparison, as dlPFC L4 had the highest minimum number of DEGs compared to all other dlPFC spatial domains. This result indicates that there was no singular dACC spatial domain that was unequivocally mapped to dlPFC L4. Also similar to the pseudobulked sample-level of comparison, dlPFC L6 had a smaller proportion of DEGs with dACC L6a compared to dACC L6b.

To further explore the spatial domain differences between dACC and dlPFC, we compared the log fold-change (logFC) values from the enrichment model DE results. L5 in dACC and L5 in dlPFC have a similar relationship to L6a in dACC and L6 in dlPFC, while L6b in dACC and L6 in dlPFC have a very low correlation (**Fig. 5D**). logFC plots comparing the regions highlight the more substantial differences in the deeper layers (L5 and L6) compared to higher similarity in the more superficial layers (L1, L2, and L3) (**Fig. 5D, Supplementary Fig. 32**). While correlation across the regions for L2 was strong, we noted several notable differences, including *BDNF* and *RSPO2*. Consistent with our data in **Fig. 2G,H**, this analysis provides additional molecular evidence for the lack of an analogous L4 layer in the dACC. As noted above, dlPFC L4 had the highest minimum number of DEGs compared to all other dlPFC spatial domains, showing that it did not have a direct pairing in the set of dACC spatial domains (**Fig. 5B**). Moreover, in the scatterplots, we note that at the gene level, *RORB* and *PVALB*, which are established as L4 markers in granular cortical areas, are enriched in dACC L5. We also noted that another class of dACC enriched genes in L5 are expressed in VENs (*POU3F1, HAPLN4)*, providing additional evidence for localization of molecularly-defined VENs in the dACC versus dlPFC. In L6, we noted enrichment of *SSTR2* and *TBR1* when comparing L6a dACC to L6 dlPFC. We also noted enrichment of *CPLX3*, *ADRA2A*, and *NOS1* when comparing L6b dACC to L6 dlPFC, highlighting significant differences in deeper layers across dACC and dlPFC. Furthermore, we showed moderate correlations when comparing dlPFC L4 with dACC L3 and dACC L5 (**Supplementary Fig. 33**).

Since L6a and L6b cluster as distinct spatial domains in the dACC, we further probed differences between these sub-layers. We first computed pairwise model DE genes between L6a and L6b in the dACC pseudobulked SRT data (**Supplementary Table 6**). Several established neuronal markers were more highly expressed in L6a, including *CCK*, *NPTXR*, and *NCAM2*, while several glial markers were more highly expressed in L6b, including *MBP, MOG*, and *GFAP* (**Supplementary Fig. 30H**). These data likely reflect the closer proximity of L6b to the boundary of the WM and the inclusion of a WM transition zone in the dACC. From the dACC snRNA-seq NMF factorization, we identified two NMF patterns associated with L6 cell types (NMF15 and NMF35). NMF15 was associated with L6 CT neurons, while NMF35 was associated with L6b neurons (**Fig. 3D**). Directly comparing these dlPFC and dACC cell types revealed that both the L6 CT and L6b cell types were associated with the dlPFC Excitatory L6 cell type (**Supplementary Fig. 27**). Similar to our approach in **Section 2.5**, we projected these two patterns into the dACC and dlPFC SRT datasets (**Fig. 5C**). Spot plots localizing NMF35 in the dACC revealed that this pattern was localized to the boundary of L6a and L6b, bleeding into both regions, while NMF15 was restricted to L6a. In contrast, the dlPFC spot plots showed that NMF15 is localized to L6, while NMF35 is only very sparsely expressed in L6. The data reveals that the gene expression patterns present in L6b of dACC are not strongly expressed in dlPFC.

To extend our results and provide orthogonal validation for selected findings from these analyses, we performed multiplex smFISH for *DRD5*, *KCTD8*, and *CPLX3* in dACC and dlPFC tissue sections derived from the same brain donor. *DRD5*, *KCTD8*, and *CPLX3* were identified as relatively specific markers for dACC L5, L6a, and L6b domains, respectively. In dACC, we confirmed robust and specific expression in the expected layers for all three markers. However, in dlPFC, while *KCTD8* and *CPLX3* were expressed in the deep layers, expression was not as restricted compared to the dACC. For *DRD5,* expression in dlPFC was not robust, nor localized, to L5 (**Fig. 5F**).

### 2.6 Spatial domain and cell type mapping and inference of clinical and functional datasets

dACC dysfunction has been implicated in a wide variety of neuropsychiatric disorders. To evaluate heritability across disorders and their relevant behavioral traits, we used stratified linkage disequilibrium score regression (s-LDSC). We used the z-score of per-SNP heritability to calculate FDR-adjusted *p*-values and quantify the statistical significance of the enrichment for each trait within spatial domain and cell type marker genes (**Fig. 6A,B**). We found that two spatial domains and five cell types were significantly associated (FDR < 0.05) with at least one trait. We found that the L3 spatial domain was associated with bipolar disorder, while the L6a spatial domain and L5 IT and L6 CT excitatory neurons were associated with intelligence and education years. Inhibitory *VIP*+ neurons were associated with depression, further confirming GABAergic dysregulation in major depressive disorder (Luscher et al. 2011), and as expected, microglia were positively associated with Alzheimer’s disease (Hansen et al. 2018).

**Fig. 6.**
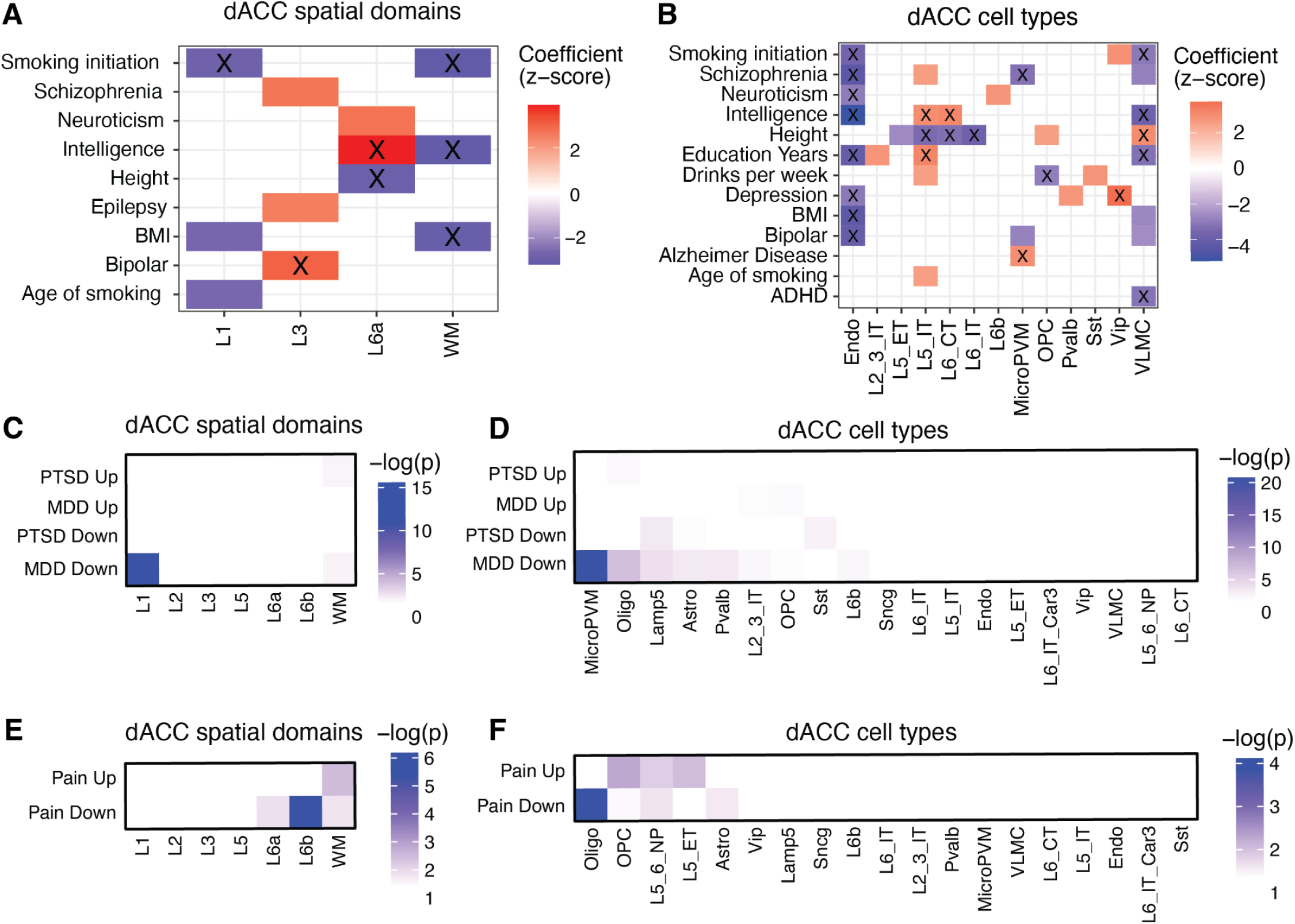
Stratified linkage disequilibrium score regression (s-LDSC) identifies differential disease risk heritability across cell types and spatial domains, and functional enrichment identifies differential enrichment across cell types and spatial domains. (**A**) Plot showing stratified linkage disequilibrium score regression (s-LDSC) coefficient *z*-scores for heritability of various polygenic traits (*y*-axis) across dACC SRT spatial domains (*x*-axis). Coefficients found to be significantly different from the baseline model following FDR correction (FDR < 0.1) are shown. Results are colored by s-LDSC regression coefficient *z*-score. The black “X” indicates stronger statistical significance (FDR < 0.05). (**B**) Same as (**A**) for dACC snRNA-seq cell types. (**C**) Heatmap showing statistical significance from Fisher’s exact test of psychiatric-related DEGs from a previous human dACC bulk RNA-seq study (Jaffe et al. 2022) (*x*-axis): MDD down: downregulated in major depressive disorder; PTSD down: downregulated in post-traumatic stress disorder; MDD up: upregulated in major depressive disorder; PTSD up: upregulated in post-traumatic stress disorder; to dACC spatial domains (*y*-axis). (**D**) Same as (**C**) for dACC snRNA-seq cell types. (**E**) Heatmap showing statistical significance from Fisher’s exact test of pain-related DEGs from a previous mouse ACC bulk RNA-seq study (Becker et al. 2023) (*x*-axis): Pain down: downregulated in pain; Pain up: upregulated in pain; to dACC spatial domains (*y*-axis). (**F**) Same as (**E**) for dACC snRNA-seq cell types.

As a complementary approach to heritability analyses, we next leveraged transcriptional signatures from case-control bulk RNA-sequencing of the human dACC (Jaffe et al. 2022) to test whether genes differentially expressed in PTSD or MDD were enriched within specific spatial domains and cell types using a two-sided Fisher’s exact test (**Fig. 6C,D**). Downregulated MDD DEGS showed significant enrichment (*p* < 0.05) in both L1 and WM, while upregulated PTSD DEGs were primarily associated with WM. Among dACC cell types, downregulated MDD DEGs were enriched in non-neuronal populations, including microglia, oligodendrocytes, astrocytes, and OPCs, as well as *LAMP5*+ and *PVALB+* inhibitory neurons. Downregulated PTSD DEGS were also associated with astrocytes and *LAMP5*+ and *SST+* neurons. In contrast, upregulated MDD DEGs were associated with L2/3 IT neurons and OPCs, while upregulated PTSD DEGs were specifically enriched in oligodendrocytes. Together, these findings reinforce a central role for both GABAergic dysfunction and dysregulation of glial and immune-related processes in the pathophysiology of MDD and PTSD, with convergent evidence implicating *LAMP5*, *PVALB*+, and *SST*+ inhibitory neurons in disease-related transcriptional changes within the dACC.

Given the prominent role of the dACC in pain perception and processing (Lieberman and Eisenberger 2015), we next asked whether transcriptional signatures associated with chronic pain states map onto specific spatial domains and cell types in the dACC. We used a bulk RNA-sequencing dataset that analyzed gene expression changes in the ACC in a mouse model of repeated activation of ACC innervation from the basolateral amygdala that is associated with behaviors linked to neuropathic pain and depression (Becker et al. 2023). Using the same enrichment framework as above, we assessed whether genes differentially expressed in the ACC pain model mapped onto human dACC spatial domains and cell types. We found that downregulated genes were concentrated in the deep-layer domains L6a and L6b, whereas upregulated genes were selectively enriched in WM. At the cellular level, we found that downregulated genes were most strongly associated with oligodendrocytes, L5/6 NP neurons, and astrocytes, whereas upregulated genes were associated with OPCs, L5 ET, and L5/6 NP neurons (**Fig. 6E**). These findings implicate deep-layer excitatory neurons in the dACC with chronic pain-related circuit dysfunction (Bhattacherjee et al. 2023), consistent with their known projection patterns to subcortical structures critical to sensory processing, affect, and motor control (Lui et al. 2021). More broadly, this analysis underscores the utility of our newly created spatial and snRNA-seq atlases as a valuable tool for linking circuit-level dysfunction to molecular and cellular pathology in neuropsychiatric disorders.

### 2.7 Data Access and Visualization

`SpatialExperiment` and `SingleCellExperiment` R objects used for analysis are hosted at a public GLOBUS endpoint (https://research.libd.org/globus/). FASTQ raw data, as well as `CellRanger` and `SpaceRanger` output files, can be found through GEO accessions GSE296731 and GSE296789, respectively. Single-nucleus RNA-sequencing data can be accessed through an `iSEE` app (Rue-Albrecht et al. 2018) and SRT data can be accessed through the `spatialLIBD` app (Pardo et al. 2022) at https://research.libd.org/spatialdACC/#interactive-websites. A website has been created for the project at research.libd.org/spatialdACC/ with links at the bottom for `Samui Browser` visualizations (Sriworarat et al. 2023) along with `iSEE` and `spatialLIBD` apps. The project code is available on GitHub at https://github.com/LieberInstitute/spatialdACC and has been archived at Zenodo (2025).

## 3 Discussion

The ACC controls a wide variety of functions related to emotional regulation and cognition. Individual ACC subdivisions, which include the dACC, as well as the more ventral sgACC and pgACC (2009; Stevens et al. 2011; van Heukelum et al. 2020), have specialized roles across these broad functions. The diversity in cell type composition and topography of efferent and afferent connections across the subdivisions is hypothesized to underlie these individual differences in function. In the context of cognition, the dACC is specifically implicated in response selection, conflict monitoring, and error detection, particularly in the context of reward processing (Devinsky et al. 1995; Vogt et al. 1995; Carter et al. 1998, 2009; Heilbronner and Hayden 2016). The dACC is also strongly linked to processing of attention. In the context of emotional regulation, the dACC is specifically implicated in regulation of pain and fear expression. To investigate spatio-molecular features of the dACC that could reflect the region’s unique structure and function, we generated paired snRNA-seq and SRT data from adjacent dACC tissue sections. To facilitate comparative molecular neuroanatomy analyses that could provide supporting evidence for the molecular rationale underlying evolutionary specializations in the dACC, data was generated in the same ten donors, and using the same experimental protocols and design, as in our previous molecular profiling study of the human dlPFC (Huuki-Myers et al. 2024).

Using unsupervised clustering of the dACC SRT data, we isolated seven unique spatial domains, which we annotated as L1, L2, L3, L5, L6a, L6b, and WM. While the agranular organization of the dACC has long been appreciated from a cytoarchitectural perspective, the molecular basis for this organization is less well understood. While molecular differences that support ACC agranularity were previously investigated in snRNA-seq data from sgACC and dlPFC of paired donors (Kim et al. 2023), SRT information that provides gene expression within the precise X-Y coordinates of the profiled tissue versus relying on inferred positioning is critical for laminar context. Moreover, snRNA-seq fails to capture extranuclear transcripts, and given the expanded amount of neuropil in the human brain (Spocter et al. 2012), cytoplasmic transcripts are likely to significantly contribute to overall molecular laminar identity. We used the dlPFC SRT and snRNA-seq data to determine that expression of several genes that are canonically enriched in L4, including *RORB* and *PVALB,* are substantial in both L4 and L5 spatial domains and corresponding cell types. However, in the dACC, expression of these genes is limited to the L5 spatial domain and L5 cell types. Expression of these markers is not enriched in the L5-adjacent L3 spatial domain nor in L3 cell types, confirming that the spatial clustering for dACC with no L4 spatial domain accurately reflects the laminar structure of the region. Ability to query the integrated snRNA-seq and SRT allowed us to provide compelling molecular evidence for the absence of L4 in dACC in the human brain.

VENs are an understudied cell type that are present in a selected set of large-brained animals and are unique to only a handful of cortical regions (Hodge et al. 2020). VENs warrant further exploration given that their loss is associated with schizophrenia (Brüne et al. 2010), suicidal psychosis (Brüne et al. 2011), and autism spectrum disorder (Santos et al. 2011). We identified an NMF pattern from the snRNA-seq data that corresponded to VENs. Projection of this pattern into the SRT data enabled us to better characterize the spatial localization of VENs, which we mapped to deep L5 in the dACC. Both data modalities were necessary for this discovery, as the differential expression of SRT spatial domains would aggregate together all cell types and sublayer patterns within L5 and compare it against the gene expression of the other spatial domains. In addition to discovering the sub-localization, we were able to infer connectivity patterns of VENs using mice models. The ACC is highly connected to both the prefrontal cortex and to subcortical regions within the limbic system, and the topography of connections to these areas differs across subdivisions of the ACC (Stevens et al. 2011; van Heukelum et al. 2020).

Substantial inter-individual variability in gene expression across unique brain donors is a significant caveat of postmortem human tissue studies that complicates comparative analyses across cortical regions, which are relatively similar in molecular and cellular structure. To leverage the fact that the dACC data was generated in the same donors as our previous dlPFC study (Huuki-Myers et al. 2024), we deployed comparative, within-donor comparisons of the two brain regions, which revealed greater divergence between regions in the deeper layers compared to the superficial layers. Specifically, we noted substantial differences in L5 molecular composition, with some being driven by gene expression patterns in deep L5, corresponding to VENs, which were not present in the paired dlPFC tissue. We also noted substantial differences in the molecular composition of L6 - both in spatial localization of known L6 marker genes, and also identifying both L6a and L6b markers in the dACC that are entirely divergent from the dlPFC. We also note that the dACC, dlPFC, and hippocampus are highly connected in the human brain, and that similar cross-region comparison computational strategies can be adapted to assess molecular coherence across connected regions to investigate cellular communication patterns. We note that the data generated in our human hippocampus snRNA-seq and SRT data resource also used the same ten neurotypical donors used to generate the data here (Thompson et al. 2024). To make this dACC data resource easily accessible for such types of future analyses and data analysis methods development or benchmarking, we made all raw and processed data freely available through GEO and several interactive web applications.

## 4 Methods

The font Courier New is used when referring to software and code.

### 4.1 Postmortem human tissue samples

Postmortem human brain tissue from the Lieber Institute for Brain Development (LIBD) Human Brain and Tissue Repository was collected through the following sites and protocols at the time of autopsy with informed consent from the legal next of kin: the Office of the Chief Medical Examiner of the State of Maryland, under the Maryland Department of Health IRB protocol #12–24, the Departments of Pathology at Western Michigan University Homer Stryker MD School of Medicine, at the University of North Dakota School of Medicine and Health Sciences, and the County of Santa Clara Medical Examiner-Coroner Office in San Jose, CA, all under the WCG protocol #20111080. Demographics for the 10 neurotypical control donors of European ancestry are listed in **Supplementary Table 1**. Details of tissue acquisition, handling, processing, dissection, clinical characterization, diagnoses, neuropathological examinations, and quality control measures have been described previously (Lipska et al. 2006). Fresh frozen coronal slabs at the level of the anterior striatum (caudate nucleus and putamen) were rapidly sub-dissected with a hand-held dental drill, perpendicular to the pial surface, targeting the dACC Brodmann Areas (BA) 33 and 24. Dissected blocks contained the following landmarks: corpus callosum ventrally, the anterior portion of the cingulate sulcus dorsally, white matter of the forebrain laterally, and the interhemispheric fissure medially. Brain blocks were approximately 8 × 8 × 5 mm in dimensions, were collected across the callosal sulcus to preserve the integrity of layer 1, and encompassed complete laminar structure, spanning all cortical layers of BA 33 and 24 and white matter. Tissue blocks were stored at −80°C in sealed cryogenic bags until cryosectioning.

### 4.2 Tissue processing and anatomical validation

Following block dissection, smaller tissue blocks were embedded in cold OCT and flash frozen in isopentane on dry ice and stored at −80°C in sealed cryogenic bags. Larger tissue blocks were flash frozen on dry ice and stored at −80°C in sealed cryogenic bags. At the time of cryosectioning, tissue blocks were mounted onto round chucks with OCT, acclimated to the cryostat (Leica CM3050s) at −14°C, ∼50 µm of tissue was trimmed from the block to achieve a flat surface, and several 10 µm sections were collected for quality control (RNAScope, H&E). Three anatomical validation measures were implemented: 1) visual inspection of the blocks to assess inclusion of the callosal sulcus, white and grey matter; 2) H&E staining to assess cellular integrity of cortical lamina and the white matter; and 3) multiplex RNAScope single molecule fluorescence *in situ* hybridization (smFISH) to ensure presence of molecular markers for cortical layers and the white matter (**Supplementary Fig. 2**). H&E staining was performed as previously described and images were acquired using an Aperio CS2 slide scanner (Leica) equipped with a 20x/0.75NA objective and a 2x doubler. RNAScope smFISH was performed as previously described (Maynard et al. 2020), and images were acquired using the Nikon AX-R confocal microscope, equipped with a 2x/20x NA objective or a Vectra Polaris slide scanner (Akoya Biosciences). Probes for established marker genes (ACD Bio) were used to identify cortical laminae in the human cortex(Maynard et al. 2021), including *SLC17A7* (Cat No. 415611) indicative of all present layers; *RELN* (Cat No. 413051) indicative of layer 1; and *MBP* (Cat No. 411051) indicative of white matter. Subsequent to the anatomical validation experiments to verify the presence of all cortical layers, blocks were again acclimated to the cryostat, mounted onto chucks, and scored with a razor to isolate Brodmann Area 33 and 24 and white matter in 6.5 × 6.5 mm squares. Adjacent 10 µm tissue sections were mounted onto the Visium Spatial Gene Expression Slide (catalog no. 2000233, 10x Genomics) and onto glass slides for further anatomical validation (RNAScope). Four independent donors were processed on any individual Visium slide, with a total of n=17 capture areas processed. For some donors, biological replicates (2-3 additional capture areas per donor) were generated.

### 4.3 Spatially-resolved transcriptomics (SRT) data generation

Scored 10 µm sections from each of the 10 dACC tissue blocks (**Supplementary Fig. 2**), were collected onto a 10x Visium Gene Expression slide, and processed according to manufacturer’s instructions (protocol number CG000239, Rev G, 10X Genomics) as previously described (Maynard et al. 2021; Huuki-Myers et al. 2024). Samples were processed with 4 different donors mounted on each Visium slide. Three of the donors were processed 3 times (3 spatially-adjacent replicates each), one of the donors was processed twice (2 replicates) and the remaining 6 donors were processed once (1 replicate each) (**Supplementary Table 1**). Prior to each Visium assay, H&E staining was performed (protocol CG000160, revision B, 10x Genomics) and high-resolution, brightfield images were acquired on a Leica CS2 slide scanner with a 20x/0.75NA objective and a 2x doubler. Following imaging, tissue sections were permeabilized for 18 minutes, reverse transcription was performed directly on the Visium slide and cDNA was collected from the slide, followed by library construction. The resulting libraries were quality controlled and sequenced on Illumina Nova-Seq 6000 following manufacturer’s instructions using a loading concentration of 300 pM at a minimum depth of 60,000 reads per Visium spot. Samples were sequenced to a median depth of 275,278,056 reads, corresponding to a median 67,211 of mean reads per spot, a median 2,257 of median unique molecular indices (UMIs) per spot, and median of 1,380 median genes per spot.

### 4.4 SRT data processing and analysis

#### Visium H&E image processing

Sample slide images were first processed using `VistoSeg` (Tippani et al. 2023). `VistoSeg` was used to divide the Visium sample slides into individual images using VistoSeg’s `splitSlide` function. This takes one large image and separates them into distinct capture areas, one for each area on the slide, labeled at A1, B1, C1, and D1. These individual capture area images were used as one of the inputs for `SpaceRanger` (10x Genomics).

#### Visium raw data processing

The individual images from `VistoSeg` were then aligned with the slide capture areas in the `Loupe Browser` (10x Genomics). This allows for the alignment of the image with the spots captured on the slide. Sample slides were then processed using the `SpaceRanger` (10x Genomics) version 1.3.1 which takes the json output from the `Loupe Browser`, the sample image, and associated FASTQ files in order to generate spatial feature counts for a sample. The `SpaceRanger` counts were input into the `spatialLIBD` Bioconductor package (Pardo et al. 2022) to create a `SpatialExperiment` object (Righelli et al. 2022).

#### Visium data quality control

We removed genes with zero UMI counts across all spots, spots with all zero counts, and spots outside of the tissue defined by the Loupe Browser. We used the `addPerCellQC()` function from the `scuttle` Bioconductor package to generate spot-level quality control metrics, including library size defined as the total number of UMIs per spot, number of detected genes, and mitochondrial expression rate (McCarthy et al. 2017). We used the `isOutlier()` function from the `scuttle` Bioconductor package to explore spots with low library size and/or spots with low numbers of detected genes based on a 3x median-absolute-deviation (MAD) threshold (McCarthy et al. 2017). These low quality spots were enriched in white matter (**Supplementary Fig. 3**). To prevent not removing spots just from the white matter, we used an extreme filter to remove spots with less than 20 detected genes and/or spots with less than 20 total UMI counts. The threshold of 20 was chosen to conservatively remove only spots that we were confident had no expression. The final spots that were discarded are in **Supplementary Fig. 4**. Across many tissue sections, the gradient of mitochondrial expression rate followed gradients in histology, so we did not use this metric to remove spots (**Supplementary Fig. 5**). We removed one spot with 100% mitochondrial expression rate. We calculated the log_2_-transformed normalized expression matrix after removing spots using `logNormCounts()` from the `scater` Bioconductor package (McCarthy et al. 2017).

#### Feature selection, spatially variable genes

We used `nnSVG`, which implements nearest neighbor Gaussian processes, to detect spatially variable genes (SVGs) (Weber et al. 2023). We ran `nnSVG` within each sample and then combined the gene sets. Within each sample, we removed lowly expressed genes using `filter_genes()` to retain genes with at least 3 counts in at least 0.5% of spatial locations. Then, we re-calculated the log_2_-transformed normalized expression matrix for each sample after removing spots using `logNormCounts()` from the `scater` Bioconductor package (McCarthy et al. 2017). We used the `nnSVG()` function from the `nnSVG` Bioconductor package on each tissue section to rank the genes in terms of spatial variance. Then, we combined the gene sets following the recommendations in the `nnSVG` tutorial by calculating the mean of the ranks of each gene across all the samples to generate an overall ranking and then calculating the number of samples where each gene is within the top 1,000 SVGs for that sample. Using the second metric, we created a table of 1,487 “replicated” genes that were highly ranked in at least two samples. Finally, the genes were ranked within the set of “replicated” genes using the average overall ranking. The gene list is available in **Supplementary Table 2**.

#### Unsupervised clustering of spatial transcriptomics data

Guided by the set of SVGs, we used `PRECAST` to integrate the samples and identify spatial domains (Liu et al. 2023). We converted from the `SpatialExperiment` object to a `Seurat` object to a `PRECAST` object using `CreateSeuratObject()` from the `Seurat` CRAN package (Hao et al. 2024) and `CreatePRECASTObject()` from the `PRECAST` CRAN package. To prepare for `PRECAST` model fitting, we added an adjacency matrix using `AddAdjList()` and parameter settings using `AddParSetting()`with the default arguments. We used the `PRECAST()` function to identify clusters with the number of clusters ranging from 5 to 20. An example of `PRECAST` annotations with 5 to 20 spatial domains is shown in one sample, V12N28-334_C1 (**Supplementary Fig. 6**).

#### Evaluation of unsupervised clustering of spatial transcriptomics data

We used two metrics to evaluate the `nnSVG`-guided `PRECAST` spatial domains to help inform which *k* value to choose for downstream analyses (**Supplementary Fig. 7**). First, we calculated the purity of each domain for each gene, defined as the proportion of observations in its neighborhood from a different domain, with the function `neighborPurity()` from the `bluster` Bioconductor package (Lun 2024). We group the data by cluster and use the average purity of genes per cluster. The cluster purity metric showed similar purity from *k*=5 to 8 clusters, with a drop in purity after 9 clusters. Second, we used the metric H_+_, which measures the discordance of unsupervised clusters as an internal validity metric (Dyjack et al. 2023). The H_+_ metric showed a decrease in discordance from *k*=5 to 7 clusters, with similar discordance values from 6 to 10.

#### Spatial domain annotation

Histology-driven spatial domains of the data were generated and annotated by manual classification of spots into domains using the `Samui Browser`, a web-based interactive tool for gene expression visualization overlaid with histology (Sriworarat et al. 2023). For spot classification, one sample from each donor was annotated based on anatomical features on the H&E images and expression of established gene markers for cortical layers (e.g. *AQP4* for L1, *HPCAL1* for L2/3, *PCP4* for L5, *KRT17* for L6) and *MBP* for white matter. The difference between white matter of the forebrain adjacent to L6 vs. white matter of the corpus callosum was annotated entirely based on histology and the H&E images because the same white matter marker gene, *MBP*, was used to annotate both domains due to its abundant expression. The final histology-driven annotations are shown in **Supplementary Fig. 9**. We also annotated the unsupervised spatial domains that were generated by `PRECAST` (*k*=9). We chose *k*=9 because this resolution generated contiguous spatial domains that resembled L1, L2, L3 and L5, based on established marker genes in the literature and in manually annotated layers of the human cortex (*RELN* for L1, *LAMP5* for L2, *ADCYAP* for L3, *PCP4* for L5 and *NR4A2* for L6). This resolution (*k*=9) also produced three domains with established markers of WM (MOBP). However, one of these three domains was defined by low library size and they did not correspond to any anatomically-distinct WM regions and were not identified during RNAScope anatomical validation. (**Supplementary Fig. 8**). We removed spots from this spatial domain from further downstream analysis. The remaining two WM domains were collapsed based on similarities in marker genes for WM. To increase confidence in annotation of the final domains, we performed spatial registration to show the similarity between the domains that were generated on manual classification of spots and the data-driven `PRECAST` domains. We utilized the DE enrichment model *t*-statistics (see next sections “Pseudobulking” and “Pseudobulking differential expression analysis”) from all data types for the spatial registration. To calculate a correlation matrix between the statistics, we used `layer_stat_cor()` from the `spatialLIBD` Bioconductor package (Pardo et al. 2022), specifying that the function keeps the top 100 marker genes (`top_n`). We used `layer_stat_cor_plot()` from the same package to visualize the correlation matrix as a heatmap.

#### Pseudobulking

In order to detect differentially expressed genes (DEGs) between the spatial domains, we pseudobulked the data, similar to (Maynard et al. 2021), where we aggregated total UMI counts across all spots within a spatial domain and within a tissue section. We used `registration_pseudobulk()` from the `spatialLIBD` Bioconductor package (Pardo et al. 2022) to pseudobulk the spatial domains and calculate a matrix approximate to log_2_(CPM + 1) via `cpm()` from `edgeR` of the pseudobulked counts matrix. This resulted in a counts matrix with 13,576 genes and 119 samples. We visualized the percent variance explained and the first two principal components computed from the log_2_-transformed normalized pseudobulked matrix using `getVarianceExplained()`, `plotExplanatoryVariables()`, and `plotPCA()` functions from the `scater` Bioconductor package (McCarthy et al. 2017) (**Supplementary Fig. 12**). There was 1 white matter pseudobulked sample with an extremely large PC2 value of 339 (sample V12N28-332_B1), so we removed this prior to downstream analyses.

#### Pseudobulking differential expression analysis

We conducted differential expression (DE) analysis using the pseudobulked samples via enrichment statistics to test if one cluster has greater expression than the rest, similar to (Thompson et al. 2024). We used the functions wrapped into `registration_wrapper()` from the `spatialLIBD` Bioconductor package to compute gene enrichment t-statistics. We utilized the false discovery rates and log_2_ fold-change values from the gene enrichment tests to create volcano plots using `EnhancedVolcano()` from the `EnhancedVolcano` Bioconductor package (Blighe et al. 2018) with the absolute log_2_ fold-change cut-off as 1.5 and the statistical significance cut-off as − log_10_(0.05) (**Supplementary Fig. 30**). We used `sig_genes_extract()` from the `spatialLIBD` Bioconductor package to extract the top 50 significant genes from each spatial domain (**Supplementary Table 5**).

To compute pairwise DE genes between L6a and L6b, we used the same functions to extract the pairwise DE genes and visualize the results in a volcano plot (**Supplementary Fig. 30H**).

### 4.5 snRNA-seq data generation

#### snRNA-seq data collection and sequencing

Single-nucleus RNA-sequencing (snRNA-seq) was performed using 2-3 100μm cryosections collected from each donor brain block with 10x Genomics Chromium Single Cell Gene Expression V3 technology.

Approximately 70-100mg of tissue was collected from each donor, placed in a pre-chilled 2mL microcentrifuge tube (Eppendorf Protein LoBind Tube, Cat #22431102), and stored at −80°C until the time of experiment. To isolate nuclei, cryosections were combined with chilled Nuclei EZ Lysis Buffer (MilliporeSigma #NUC101) into a glass dounce. Sections were homogenized using 10-20 strokes with both loose and tight-fitting pre-chilled pestles. Homogenates were filtered through 70 μm mesh strainers and centrifuged at 500g for 5 minutes at 4°C using a benchtop centrifuge. Nuclei pellets were resuspended in fresh EZ lysis buffer, centrifuged again, and resuspended in wash/resuspension buffer (1x PBS, 1% BSA, 0.2U/μL RNase Inhibitor). Final nuclei were washed in wash/resuspension buffer and centrifuged a total of 3 times. Alexa Fluor 488-conjugated anti-NeuN antibody (MilliporeSigma cat. #MAB377X) diluted 1:1000 in nuclei stain buffer (1x PBS, 3% BSA, 0.2U/μL RNase Inhibitor) was used to label nuclei by incubating at 4°C with continuous rotation for 1 hour. Proceeding NeuN labeling, nuclei were washed once in stain buffer, centrifuged, and resuspended in wash/resuspension buffer. Nuclei were labeled with propidium iodide (PI) at 1:500 in wash/resuspension buffer and subsequently filtered through a 35μm cell strainer. Fluorescent activated nuclear sorting (FANS) was performed using a Bio-Rad S3e Cell Sorter at the Lieber Institute for Brain Development. Gating criteria were selected for whole, singlet nuclei (by forward/side scatter), G0/G1 nuclei (by PI fluorescence), and neuronal nuclei (by Alexa Fluor 488 fluorescence). First, 9,000 nuclei were sorted based on PI+ fluorescence to include both neuronal and non-neuronal nuclei from each donor. Second, 9,000 additional nuclei were sorted into a separate tube based on both PI+ and NeuN+ fluorescence to facilitate enrichment of neurons. This resulted in a final N=20 for snRNA-seq (1 PI+ and 1 PI+NeuN+ sample for all 10 donors) with a total of 18,000 sorted nuclei per donor. Samples were collected over multiple rounds, each containing 3-4 donors for 6-8 samples per round. All samples were sorted into reverse transcription reagents from the 10x Genomics Single Cell 3′ Reagents kit (without enzyme). Reverse transcription enzyme and water were added to bring the reaction to full volume. cDNA synthesis and subsequent library generation were performed according to the manufacturer’s instructions for the Chromium Next GEM Single Cell 3’ v3.1 (dual-index) kit (CG000315, revision E, 10x Genomics). Samples were sequenced on a NovaSeq 6000 (Illumina) at the Johns Hopkins University Single Cell and Transcriptomics Sequencing Core.

### 4.6 snRNA-seq data analysis

#### snRNA-seq data processing and quality control

We removed empty droplets as the first step of quality control for the snRNA-seq data. We used `barcodeRanks()` from the `DropletUtils` Bioconductor package (Griffiths et al. 2018; Lun et al. 2019) to incorporate unique sample-informed thresholds for each sample. We used the barcode rank statistics to find cliff and knee points for each sample (**Supplementary Fig. 14**). To identify empty droplets, we used `emptyDrops()` from the `DropletUtils` Bioconductor package with 25,000 iterations (`niters`) and the knee points as lower bound on the total UMI count (`lower`) for each sample.

We removed genes with zero UMI counts across all nuclei and nuclei with all zero counts. Then, we wanted to remove low quality nuclei using doublet detection, mitochondrial expression rate, and library size before moving on to the next step in the analysis pipeline. We used the `addPerCellQC()` function from the `scuttle` Bioconductor package to generate nuclei-level quality control metrics, including library size, number of detected genes, and mitochondrial expression rate (McCarthy et al. 2017). We used the `isOutlier()` function from the `scuttle` Bioconductor package to explore low library size nuclei, nuclei with low numbers of detected genes, and nuclei with high mitochondrial expression rate based on a 3x median-absolute-deviation (MAD) threshold (McCarthy et al. 2017). We discarded these low quality nuclei (**Supplementary Fig. 15**, all low sum nuclei were also low detected nuclei). We used `scDblFinder()` from the `scDblFinder` Bioconductor package (Germain et al. 2021) to remove doublets from the dataset (**Supplementary Fig. 16**).

#### snRNA-seq feature selection and dimensionality reduction

We used the `devianceFeatureSelection()` function from the `glmpca` CRAN package (Townes et al. 2019) to calculate the Poisson deviance on the counts matrix, taking into account the donor batch variable. We used the highest Poisson deviance to select the top 2,000 features (**Supplementary Fig. 18**). To approximate GLM-PCA for dimensionality reduction, we used the `nullResiduals()` function from the `glmpca` CRAN package with Poisson likelihoods on Pearson residuals and then ran PCA using the `runPCA()` function from the `scater` Bioconductor package (McCarthy et al. 2017).

#### snRNA-seq clustering and cell type annotation

On the GLM-PCA reduced dimensions, we used `RunHarmony()` from the `harmony` CRAN package (Korsunsky et al. 2019) for batch correction. The `harmony` batch corrected data was plotted using UMAP to ensure the samples were not separated out and clustered together (**Supplementary Fig. 18**). For cell type annotation, we used the `Azimuth Human Motor Cortex` reference data to predict cell types (Bakken et al. 2021). We used the function `RunAzimuth()` from the `Azimuth` R package, which uses the `Seurat` data organization tool (Hao et al. 2021, 2024). This annotation resulted in 20 cell types, of which we removed the Sst+ Chodl+ GABAergic neuron (“Sst Chodl”) cell type due to its small proportion (n=34) (**Supplementary Fig. 17**). The Sst Chodl cell type is included in the snRNA-seq `iSEE` app (Rue-Albrecht et al. 2018) and publicly available R `SingleCellExperiment` object (Amezquita et al. 2020). We used a UMAP plot to visualize the separation of the final 19 cell types (**Fig. 3A**) using `ggplot2` (Wickham 2016).

#### Pseudobulked DE analysis

Similar to the SRT DE analysis, we conducted DE analysis using pseudobulked samples via enrichment statistics to test if one cell type has greater expression than the rest. We examined the pseudobulked samples with PCA to ensure well-mixed data before proceeding with the DE analysis (**Supplementary Fig. 19**). We used `registration_wrapper()` from the `spatialLIBD` Bioconductor package to compute gene enrichment t-statistics (Pardo et al. 2022). We utilized the false discovery rates and log_2_ fold-change values from the gene enrichment tests to create volcano plots using `EnhancedVolcano()` from the `EnhancedVolcano` Bioconductor package (Blighe et al. 2018) with the absolute log_2_ fold-change cut-off as 1.5 and the statistical significance cut-off as − log10(0.05) (**Supplementary Fig. 20**, **Supplementary Fig. 21**). We used `sig_genes_extract()` from the `spatialLIBD` Bioconductor package to extract the top 30 significant genes from each cell type (**Supplementary Table 3**). We visualized the log_2_-normalized expression of a couple of the top marker genes for each layer-specific cell type using the `ComplexHeatmap` Bioconductor package (Gu et al. 2016) (**Fig. 3**).

#### Spatial registration

We utilized the DE enrichment model t-statistics from both data types for spatial registration. To calculate a correlation matrix between the statistics, we used `layer_stat_cor()` from the `spatialLIBD` Bioconductor package (Pardo et al. 2022), specifying that the function keeps the top 100 marker genes (`top_n`). We used `layer_stat_cor_plot()` from the same package to visualize the correlation matrix as a heatmap (**Fig. 3C**).

### 4.7 Dorsolateral prefrontal cortex (dlPFC) data

#### Dorsolateral prefrontal cortex (dlPFC) SRT dataset

We leveraged dlPFC SRT data from the same ten donors to facilitate cross-region comparisons between the dACC and dlPFC. The dlPFC SRT data was downloaded using `fetch_data()` from the `spatialLIBD` Bioconductor package with the option `“spatialDLPFC_Visium“` (Pardo et al. 2022). This dataset contains 30 samples of dlPFC, where each donor has three samples. We relabeled the `BayesSpace_harmony_09` annotation to reflect spatial domain annotations. Specifically, we associated 2 to Layer 1, 3-Layer 2, 5-Layer 3, 8-Layer 4, 4-Layer 5, 7-Layer 6, 6-WM, 9-WM, and removed 1 which corresponded to meninges.

#### Dorsolateral prefrontal cortex (dlPFC) snRNA-seq dataset

Additionally, we leveraged dlPFC snRNA-seq data from the same ten donors to facilitate cross-region comparisons between the dACC and dlPFC. The dlPFC snRNA-seq data was downloaded using `fetch_data()` from the `spatialLIBD` Bioconductor package with the option `“spatialDLPFC_snRNAseq“` (Pardo et al. 2022). This dataset contains 19 tissue blocks from 10 donors from dlPFC. We used the `cellType_layer` annotations for our comparisons. We used the same functions to compute pseudobulked DE marker genes for each cell type compared to all other cell types and used the same spatial registration process to compute the correlation between dlPFC cell type markers and dACC snRNA-seq cell type markers (**Supplementary Fig. 26**).

### 4.8 Non-negative matrix factorization

#### NMF factorization

We used the log_2_-transformed normalized expression of the snRNA-seq dataset to run NMF. To decide on the number of ranks, we used the `cross_validate_nmf()` function from the `singlet` R package (DeBruine 2022b). We determined the optimal rank to be *k*=75 (**Supplementary Fig. 23**) and then used the `nmf()` function from the `RcppML` R package to factorize the expression matrix (DeBruine 2022a).

#### NMF annotation

To remove NMF patterns associated with technical variables, we created a heatmap to visualize the association of each pattern with various technical variables, including brain donor, quality control metrics, and sex of brain donor. For continuous variables, such as mitochondrial percentage, sum UMI counts, and detected UMI counts, we computed the Pearson correlation with each NMF pattern’s weights. For categorical variables, we created dummy variables of either 0 or 1 to represent exclusion/inclusion in the category and then computed the correlation with each NMF pattern’s weights. The NMF patterns found to be associated with these variables were not used for further analyses: NMF29 and NMF31 for sex; NMF62 for mitochondrial percentage; NMF4, NMF8, and NMF12 for sum/detected UMI counts; and NMF6, NMF22, and NMF64 for brain donor (**Supplementary Fig. 25**).

The NMF patterns were aggregated across cell types, and the mean of each NMF pattern within each cell type is displayed in a heatmap (**Supplementary Fig. 24**). For each NMF pattern, the top 10 genes contributing to the loadings matrix (`w` matrix in the `RcppNL::nmf()` output) were chosen as the marker gene set (**Supplementary Table 4**). We evaluated these marker genes to assure that they matched with the associated cell types for the NMF patterns as shown in the heatmap. The top NMF pattern for each cell type was selected for downstream analyses.

#### NMF pattern transfer to dACC and dlPFC SRT datasets

The annotated patterns learned from the dACC snRNA-seq were projected into various datasets using a similar procedure, first used in the human hippocampus study (Thompson et al. 2024). First, we describe transfer into the dACC SRT gene expression data. The loadings matrix and the gene expression matrix from the dACC SRT data were subset to the genes present in both matrices and ordered to match each other. Then, we used the `project()` function from the `RccpML` R package to project the loadings into the log_2_-transformed normalized dACC SRT expression data (DeBruine 2022a). The resulting patterns correspond to the same patterns annotated from the dACC snRNA-seq data. This process was used to transfer the patterns into other datasets, namely dlPFC SRT data and dlPFC snRNA-seq data.

#### NMF pattern transfer to mouse single-nucleus methylation seqeuncing (snmC-seq) dataset

The annotated patterns learned from the dACC snRNA-seq were projected into a mouse model linking axon projections to snmC-seq in neurons. The snmC-seq data was downloaded from GEO (accession code GSE230782) and reproduced a workflow used in a previous study to process the data (Thompson et al. 2024). We subset the dataset to include only ACC neurons and then log-transformed and negated the extracted CH gene body counts, approximating gene expression with the inverse of non-CpG cytosine methylation. To convert into human morphology, we matched mouse genes with human orthologs and removed those without matches. This matrix is analogous to the dACC SRT gene expression data in the previous section. The loadings matrix and this matrix were subset to the genes present in both matrices and ordered to match each other. Then, we used the `project()` function from the `RccpML` R package to project the loadings into the final matrix (DeBruine 2022a).

#### NMF pattern differential expression

We identified differentially expressed genes between two NMF patterns using the following procedure. In the dACC SRT dataset, we used NMF38 and NMF61 transferred from the dACC snRNA-seq NMF patterns. Spots that had a nonzero entry in the patterns matrix for NMF38 were classified as expressing NMF38 (n=4,191), and spots that had a nonzero entry in the patterns matrix for NMF61 were classified as expressing NMF61 (n=1,022) (**Supplementary Fig. 28**). We used a similar DE framework as above to find pairwise DE genes and corresponding t-statistics between the two classifications with `registration_stats_pairwise()` from the `spatialLIBD` Bioconductor package (Pardo et al. 2022).

### 4.9 Cross-region comparisons

#### von Economo Neurons (VENs)

We examined the distribution of von Economo neurons (VENs) in the dACC and compared this to the dlPFC. We used `make_escheR()` from the `escheR` Bioconductor package (Guo et al. 2023) to visualize spot plots of NMF38 and NMF61 expression in the dACC and dlPFC SRT data. We also created sample-average boxplots of NMF38 and NMF61 expression to compare the SRT and snRNA-seq data from both regions. Briefly, we calculated the average NMF38 and NMF61 expression for each donor within the dACC SRT Layer 5 and the dlPFC SRT Layer 5 and visualized these data points with boxplots. We also calculated the average NMF38 expression for each donor within the dACC snRNA-seq L5 IT cell type and the dlPFC snRNA-seq Excitatory L5 and visualized these data points with boxplots. Finally, we calculated the average NMF61 expression for each donor within the dACC snRNA-seq L5 ET cell type and the dlPFC snRNA-seq Excitatory L5 and visualized these data points with boxplots.

#### RNAScope Validation

Three smFISH experiments were performed on tissue sections from postmortem human dACC and dlPFC from two donors (Br6432 and Br8325) using RNAScope Multiplex Fluorescent Reagent Kit v2 (Advanced Cell Diagnostics) and 4-Plex Ancillary Kit as previously described (Maynard et al. 2020). Tissue was cryosectioned at ∼10 μm on a Leica cryostat from the same dACC tissue blocks used for Visium and independent dlPFC tissue blocks from the same donor. The following probe panels were used: 1) Hs-*POU3F1* (assigned Opal dye 570), Hs-*SULF2* (assigned Opal dye 620), Hs-*GABRQ* (assigned Opal dye 690), and Hs-*PCP4* (assigned Opal dye 520) using catalog no. 483181-C4, 502241-C3, 483171, 446111-C2, respectively (**Fig. 4G** and **Supplementary Fig. 29**); 2) Hs-*PCP4* (assigned Opal dye 620), Hs-*RORB* (assigned Opal dye 520), Hs-*ADCYAP1* (assigned Opal dye 690 using catalog no. 446111-C4, 1273361-C3, 446061-C2, 582501, respectively (**Fig. 2K,L**); 3) Hs-*KCTD8* (assigned Opal dye 570), Hs-*DRD5*-O1 (assigned Opal dye 690), Hs-*CPLX3* (assigned Opal dye 520) using catalog no. 563921-C3, 437391-C4, 818241, 487681-C2, respectively (**Fig. 5F**). Images were acquired with the same settings across tissue sections using a Nikon AX-R confocal microscope, equipped with a Nikon APO lambda D 20x / 0.80 objective NA objective.

#### VENs RNAScope quantification

Images from RNAScope experiment 1 (*POU3F1*, *SULF2*, *GABRQ*, and *PCP4*) were quantified with `HALO` (Indica labs) as previously described (Mulvey et al. 2024). `HALO` settings files can be found on GitHub under processed-data, then under 24-HALO-Analyses (https://github.com/LieberInstitute/spatialdACC/tree/main/processed-data/24-HALO-Analyses). Given the enrichment of VENs in L5, we restricted analysis to this layer using *PCP4* expression to set the boundaries of L5. Briefly, we performed *k*-means clustering at *k*=3 on both copy count and signal intensity for *PCP4* and defined cells positive for *PCP4* (*PCP4*+) as the top 2 clusters (3 minimum copy counts and minimum signal intensity of 9). Next, we spatially plotted *PCP4*+ cells and drew a polygon around them for each image using custom R scripts (Mulvey et al. 2024). For each cell (defined by DAPI nuclear IF stain with an additional 3um extrapolation for cytoplasm) within the *PCP4+* L5 polygon, we quantified the copy number of each gene and cell area. For each sample separately, we scaled the copy count by cell area divided by median cell area for that sample as recommended in (Atta et al. 2024). As copy counts may become unreliable estimates for highly expressed genes with saturated fluorescent signals, we used a Gaussian mixture model to call each cell as expressing or not expressing each gene within each sample, using `Mclust()` with *k*=2 from the `mclust` R package (Scrucca et al. 2023). The proportion of cells for each sample classified as expressing *POU3F1*, *SULF2*, and *GABRQ* is presented in **Supplementary Fig. 29A**. We also calculated the proportion of total cells in each sample that were called as expressed two of the three genes or all three genes (**Supplementary Fig. 29B**). Additionally, we calculated the Spearman’s correlation for each sample for each pair of genes. For example, the Spearman’s correlation between *POU3F1* and *SULF2* copy counts in the left region of Br8325 for dACC was 0.37 (**Supplementary Fig. 29C**). We summarized these correlations into one plot, as shown in **Supplementary Fig. 29D**. To further describe the coexpression, we visualized the average proportion of cells called as expressing one gene that are also called as expressing another gene (**Fig. 4G**). For example, to get the value of 0.45, we counted the total number of cells expressing *SULF2* in one dACC sample. Out of those cells, we found the number of cells also expressing *POU3F1* and calculated the proportion of *SULF2* cells that also expressed *POU3F1*. We took the average across the 3 dACC samples to get 0.45.

#### Pseudobulked sample-level dACC and dlPFC comparison

One way we compared the spatial domains between the dACC and the dlPFC was on the pseudobulked sample-level. We used `registration_wrapper()` from the `spatialLIBD` Bioconductor package to compute gene enrichment t-statistics for the dlPFC SRT dataset (Pardo et al. 2022). We utilized the DE enrichment model t-statistics from the dlPFC SRT and dACC SRT datasets for spatial registration. To calculate a correlation matrix between the statistics, we used `layer_stat_cor()` from the `spatialLIBD` Bioconductor package (Pardo et al. 2022), specifying that the function keeps the top 100 marker genes (`top_n`). We used `layer_stat_cor_plot()` from the same package to visualize the correlation matrix as a heatmap.

#### Pseudobulked gene-level dACC and dlPFC comparison

A second way we compared the spatial domains between the dACC and the dlPFC was on the pseudobulked gene-level. We combined the pseudobulked expression matrices from both regions into one matrix, indicating the region in the metadata with the variable “Tissue”. We only kept the genes that were in both matrices in the combined gene expression matrix. In addition to the “Tissue” variable, we also had “Donor” and “Layer” variables in the metadata for each pseudobulked sample. For each “Layer” and “Tissue” combination, we calculated the number of differentially expressed genes using the `dream` workflow (Hoffman and Roussos 2021), (Hoffman and Schadt 2016). We created a `DGEList` object from the combined pseudobulked expression matrix and normalized it using `calcNormFactors()` from the `edgeR` Bioconductor package (Robinson et al. 2010). We created a linear contrasts matrix with the format `“∼ Layer_Tissue + (1 | Donor)“`, where “Layer_Tissue” is a variable combining “Layer” & “Tissue” and “Donor” is a random effect. The fixed effect variable to be tested is thus “Layer_Tissue”. We used `voomWithDreamWeights()`, followed by `dream()` and `variancePartition::eBayes()` to estimate precision weights and regression coefficients and apply empirical Bayes shrinkage on the linear mixed model (Law et al. 2014; Hoffman and Roussos 2021). We counted a gene to be significantly DE if the absolute value of the z-score, the *p*-value transformed into a signed z-score, was larger than 1.645 and the absolute value of the log_2_-fold change was greater than 1.5. We visualized these results with a heatmap, where the number of DEGs for each comparison was scaled by the total number of DEGs for the dACC spatial domain in that comparison. To clarify, the number of DEGs between Layer 1 in dACC and Layer 3 in dlPFC was scaled by the total number of DEGs across all Layer 1 dACC comparisons to dlPFC spatial domains.

### 4.10 stratified linkage disequilibrium score regression (s-LDSC)

#### Genome annotation

To define the gene set for the dACC SRT data, we used a specificity score approach (Skene et al. 2018). We created a pseudobulked gene expression matrix using `aggregateAcrossCells()` from the `scuttle` Bioconductor package (McCarthy et al. 2017) and removed samples that had less than 50 spots. We used `filterByExpr(group = spatial domains)` from the `edgeR` Bioconductor package (Chen et al. 2025) to find an adequate expression cutoff and used that cutoff to remove genes that did not have sufficiently high expression across spatial domains. We also removed duplicated genes and any non-protein coding genes. We then computed normalized CPM values for each gene in each cluster using `cpm()` from the `edgeR` Bioconductor package (Chen et al. 2025). The gene set for each spatial domain was made up of the genes with the top 10% of expression. We added a 100Kb window upstream and downstream from each gene in the gene set for each spatial domain. This created the genome annotation for each spatial domain.

The same approach was used for the cell types in the dACC snRNA-seq data.

#### GWAS enrichment

We used the stratified linkage disequilibrium score regression (`s-LDSC`) approach from a previous study (Thompson et al. 2024) to evaluate the enrichment of heritability for a variety of traits. Following the workflow from the `LDSC` resource website (https://alkesgroup.broadinstitute.org/LDSCORE), we ran `s-LDSC` for each genome annotation with the baseline LD model v2.2. This model has 97 annotations to control for the LD between variants with other functional annotations in the genome. We used HapMap Project Phase 3 SNPs as regression SNPs, and 1000 Genomes SNPs of European ancestry samples as reference SNPs (International HapMap 3 Consortium et al. 2010; 1000 Genomes Project Consortium et al. 2015). To quantify enrichment, we used the z-score of per-SNP heritability metric. From the z-scores, we calculate adjusted p-values using the Benjamini & Hochberg procedure. To visualize the results, we included traits and domains with FDR < 0.1 and added an additional “X” if the FDR < 0.05.

### 4.11 Bulk data integration

#### Clinical enrichment of MDD and PTSD bulk RNA sequencing dataset

We utilized an existing bulk RNA sequencing dataset to test for clinical enrichment of MDD and PTSD associated DEGs within spatial domains and cell types. This dataset from Jaffe et al. (available in the online supplementary files) consists of DEGs associated with MDD and DEGs associated with PTSD compared to neurotypical donors from postmortem human brains within the dACC, dlPFC, and the amygdala (Jaffe et al. 2022). We first consider the tests of enrichment within dACC spatial domains using the dACC MDD DEGs. For the MDD DEGs, we chose the subset from the dACC (the columns were `dACC_logFC_MDD` and `dACC_adjPVal_MDD`). We kept only the genes overlapping between the results table from the bulk dataset and the enrichment model results table for the spatial domains. We further separated the MDD analysis into upregulated and downregulated DEGs. The gene set for MDD upregulated was based on an adjusted *p*-value threshold of 0.1 with positive log fold-change, while the gene set for MDD downregulated was based on an adjusted *p*-value threshold of 0.1 with negative log fold-change. The background gene set for each was the remaining genes in the results table. For each spatial domain, the gene set was based on an adjusted *p*-value of 0.05 and a log fold-change greater than 1. The background gene set for each spatial domain was the remaining genes in the enrichment model results table. We used Fisher’s exact test with the function `fisher.test(alternative = “two.sided”)` from the `stats` base R package to test for enrichment and present the *p*-values in a heatmap. The same process was used to test for clinical enrichment of the spatial domains in the PTSD DEGs gene sets (the columns were `dACC_logFC_PTSD` and `dACC_adjPVal_PTSD`). Finally, the same process was used to test for clinical enrichment of the snRNA-seq cell types with both the MDD and PTSD DEGs gene sets.

#### Pain functional enrichment from mouse ACC bulk RNA sequencing dataset

We utilized an existing bulk RNA sequencing dataset to test for functional enrichment of pain associated DEGs from mouse ACC within spatial domains and cell types. This dataset consists of DEGs associated with pain from mouse ACC (Becker et al. 2023). We downloaded the data from GEO (accession code GSE227159). We first considered the tests of enrichment for the spatial domains with the pain DEGs. To convert into human morphology, we matched mouse genes with human orthologs and removed those without matches. We kept only the genes overlapping between the results table from the mouse dataset and the enrichment model results table for the spatial domains. We further separated the pain functional enrichment analysis into upregulated and downregulated DEGs. The gene set for upregulated was based on an adjusted *p*-value threshold of 0.1 with positive log fold-change, while the gene set for downregulated was based on an adjusted *p*-value threshold of 0.1 with negative log fold-change. The background gene set for each was the remaining genes in the results table. For each spatial domain, the gene set was based on an adjusted *p*-value of 0.05 and a log fold-change greater than 1. The background gene set for each spatial domain was the remaining genes in the enrichment model results table. We used Fisher’s exact test with the function `fisher.test(alternative = “two.sided”)` from the `stats` base R package to test for enrichment and present the *p*-values in a heatmap. The same process was used to test for functional pain enrichment of the snRNA-seq cell types with both the upregulated and downregulated DEGs gene sets.

## 4.12 Code availability

All version numbers for corresponding software packages are available on GitHub at https://github.com/LieberInstitute/spatialdACC (2025).

## Acknowledgements

Portions of some figures were created with BioRender.com. We thank the LIBD neuropathology team, particularly James Tooke and Amy Deep-Soboslay, for curation of the brain samples and assistance with tissue dissections. We thank the staff and physicians at the brain donation sites, and the generosity of the brain donors and their families, without whom this work would not be possible. We thank members of the Martinowich, Maynard, Page, and Hicks labs for critical reading of the manuscript and feedback, particularly Ege Yalcinbas with support for feedback on smFISH imaging and analyses. We thank the Joint High Performance Computing Exchange (JHPCE) for providing computing resources for these analyses. Finally, we thank the families of Connie and Steve Lieber and Milton and Tamar Maltz for their generous support.

## Author Contributions

Conceived and Designed the Study: SCP, KRM, SCH, KM

Performed Experiments and Collected Data: SVB, MRV, AC, HRD, ADR, SHK, ST

Software: RAM, MT, SH, SCH

Formal Analysis: KS, MST, AC

Data Curation: KS, MST, RAM, MT, HRD, LCT, SH

Tissue Resources: JEK, TMH

Writing – original draft: KS, MST, SVB, KM

Writing – review & editing: KS, MST, SVB, SCP, KRM, SCH

Visualization: KS, MST, SVB, AC

Supervision: SCP, KRM, SCH, KM

Funding acquisition: KRM, SCH, KM

Project Administration: SCH, KM

## Supplementary Figures

**Supplementary Fig. 1.**
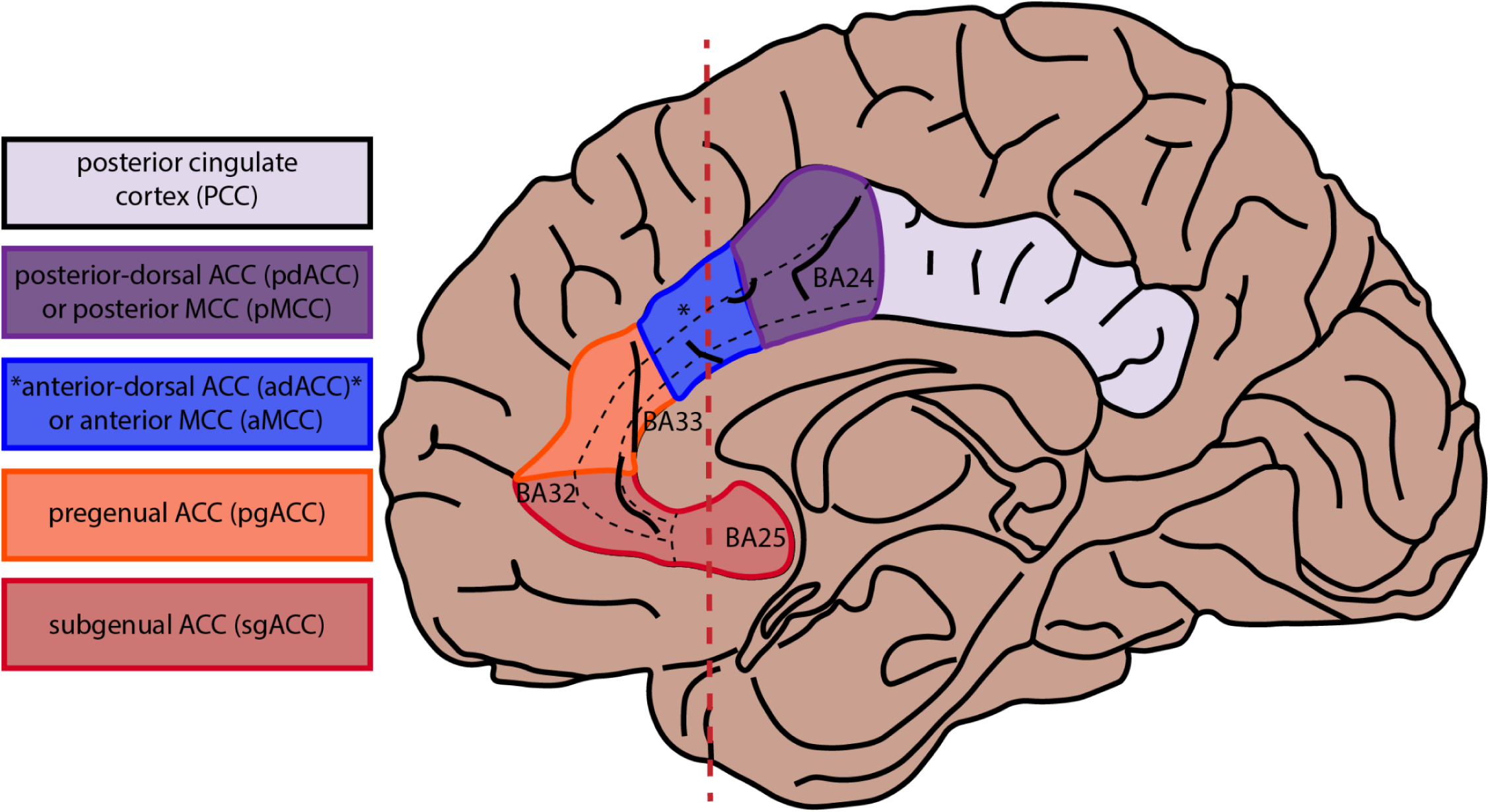
Region annotations across the ACC. Illustration of the midsagittal view of the human brain indicating five subdivisions of the cingulate cortex (CC): red - subgenual anterior CC (sgACC); orange - pregenual anterior CC (pgACC); blue - anterior-dorsal anterior CC (adACC) or anterior mid CC (aMCC); purple - posterior-dorsal anterior CC (pdACC) or posterior mid CC (pMCC); lavender - posterior CC (PCC). Approximate boundaries of Brodmann areas (BAs) 25, 33, 24, and 32 are indicated by black dashed lines across the CC. Red vertical dashed line indicates the level at which the coronal slabs were selected for this study, targeting the adACC/aMCC as the region of interest (blue*).

**Supplementary Fig. 2.**
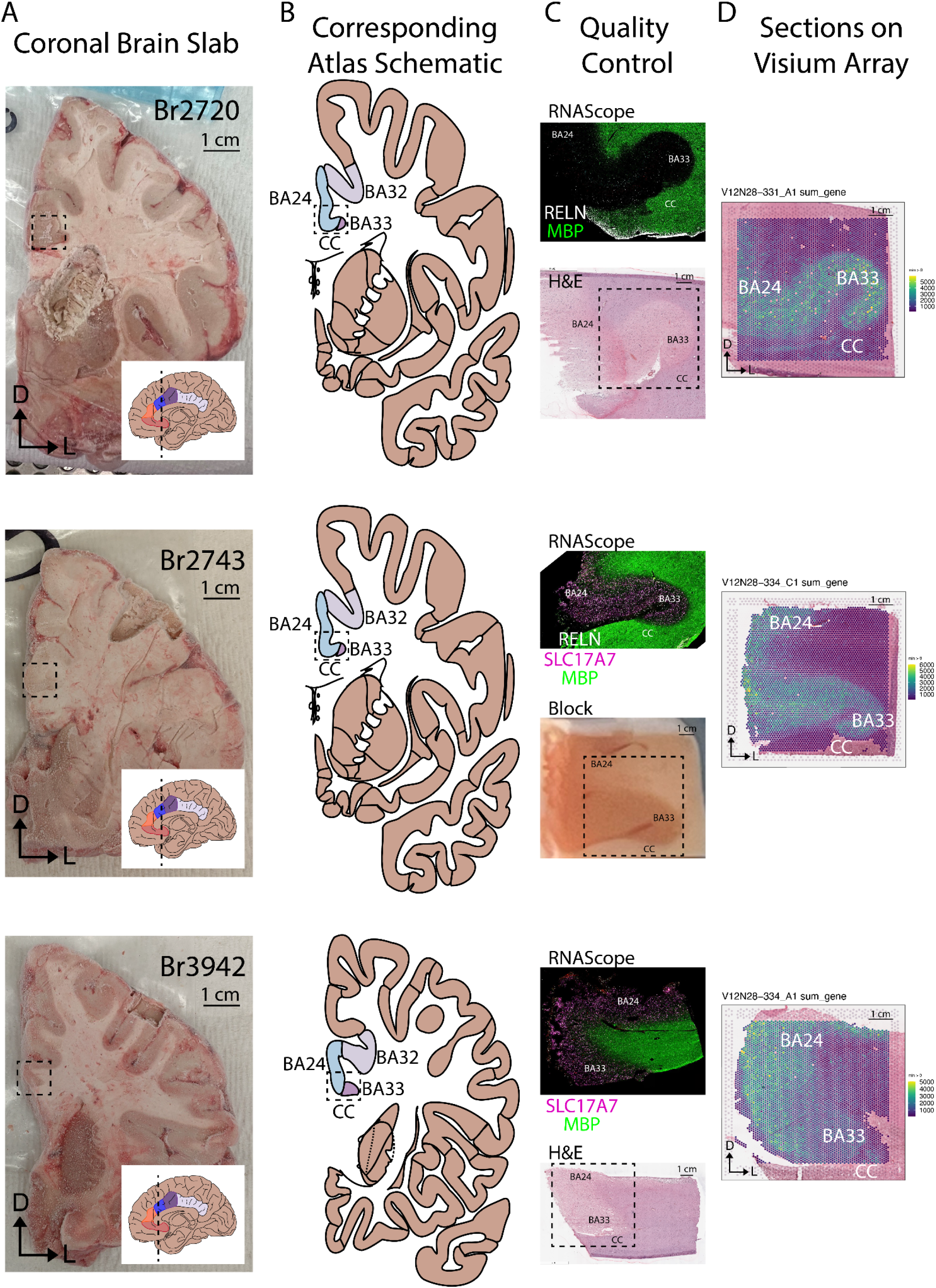
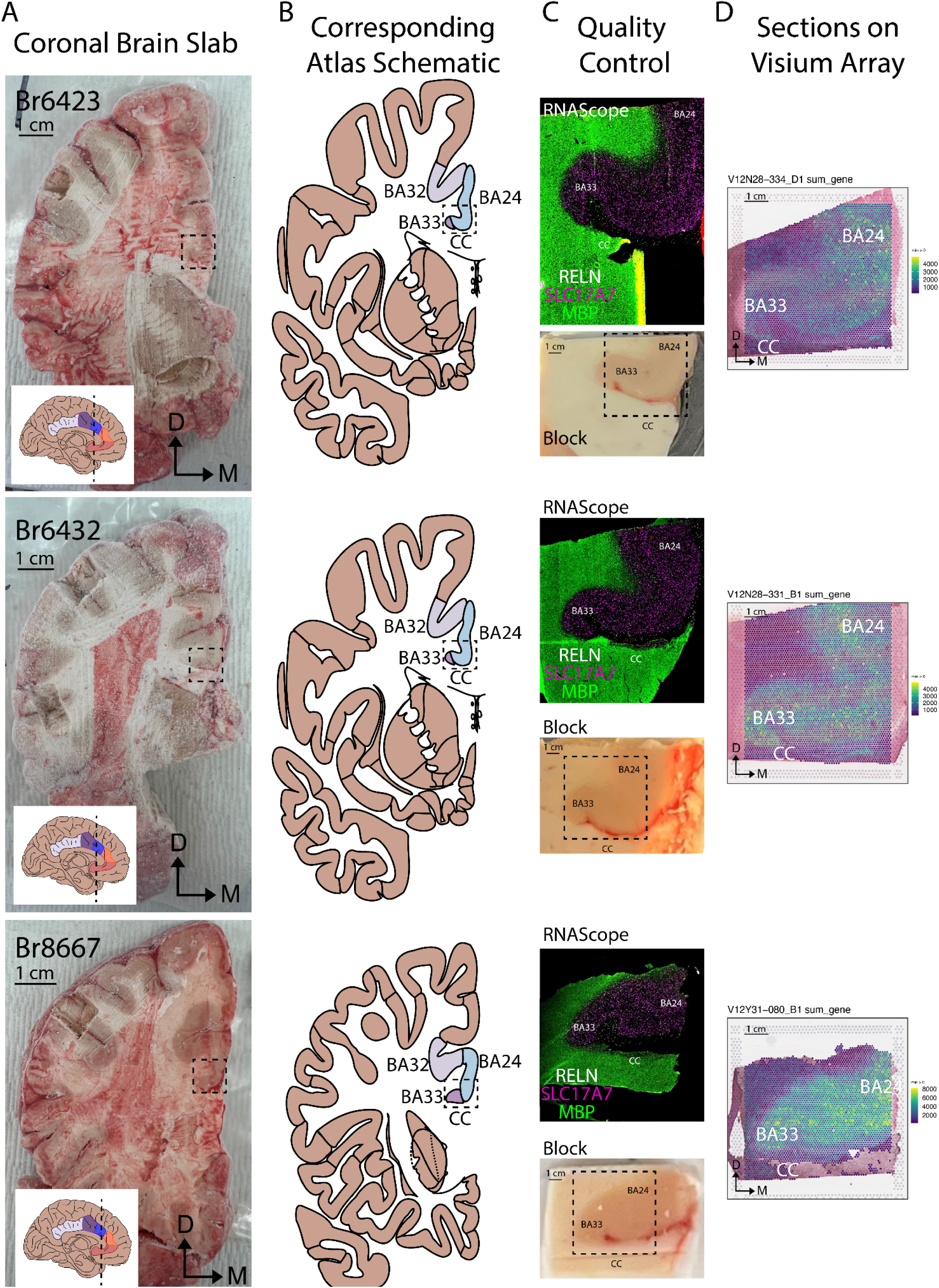
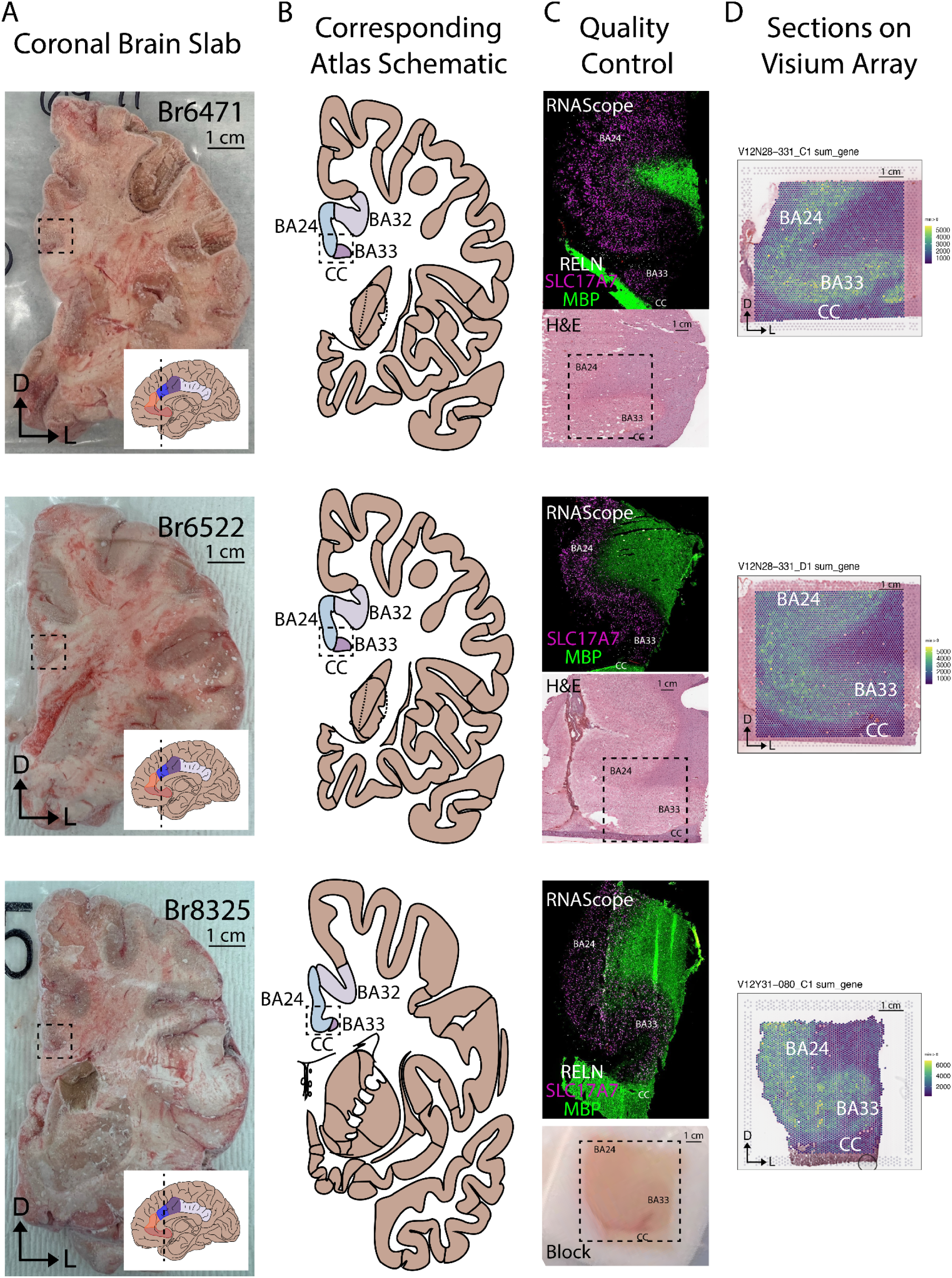
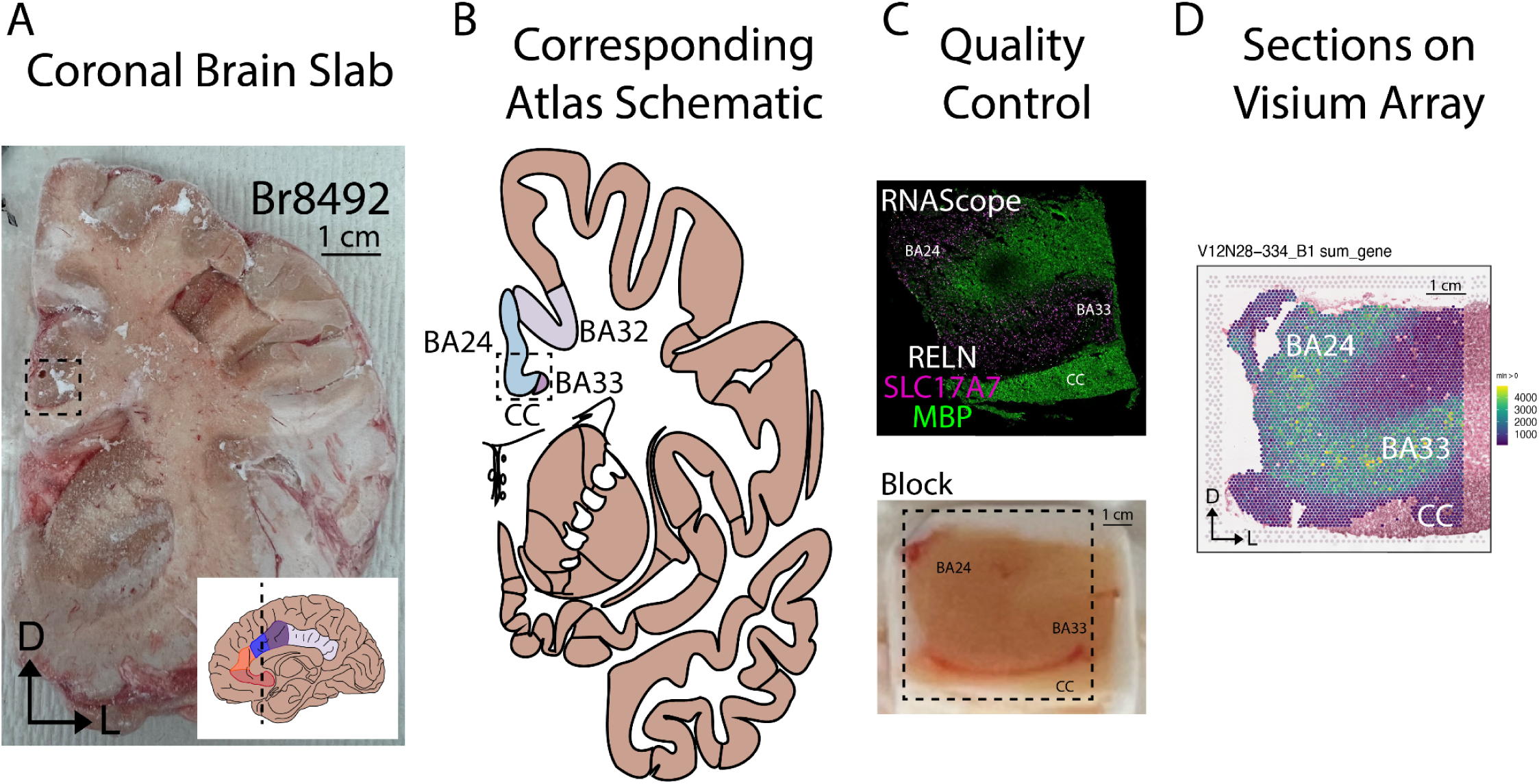
Neuroanatomical validation of dACC location across 10 donors. (**A**) Fresh-frozen coronal brain slabs from the n=10 neurotypical control donors. Dashed boxes indicate the location of the dissected blocks in (**C**). Inset depicts a midsagittal view of the brain with five subdivisions of the cingulate cortex (red - subgenual region; orange - pregenual region; blue - anterior-dorsal ACC or anterior MCC; purple - posterior-dorsal ACC or posterior MCC; lavender - posterior cingulate cortex). Dashed line across the midsagittal schematic indicates the level of the coronal slab. Neuroanatomical orientation is indicated by arrows: D - dorsal; L - lateral; M - medial. (**B**) Brain atlas schematic corresponding to the level of the coronal cut of the brain slab in (**A**). Brodmann areas (BAs), BA33, BA24, and BA32, which make up the cingulate cortex, are indicated. CC - corpus callosum. Dashed boxes indicate the location of the dissected blocks in (**C**). (**C**) Quality control experiment summary for each dissected brain block. RNAScope experiments with probes marking *MBP* (white matter), *SLC17A7* (excitatory neurons), and/or *RELN* (Layer 1), are presented in green, pink, and white, respectively. Images of H&E or unprocessed brain blocks with all relevant structures of the block are indicated: BA24, BA33, CC - corpus callosum. Dashed boxes indicate the area of tissue placed on the Visium array in (**D**). (**D**). Sections from each donor, as placed on Visium arrays, rotated to match the orientation of each slab. BAs are indicated; CC - corpus callosum. Neuroanatomical orientation is indicated by arrows: D - dorsal; L - lateral; M - medial.

**Supplementary Fig. 3.**
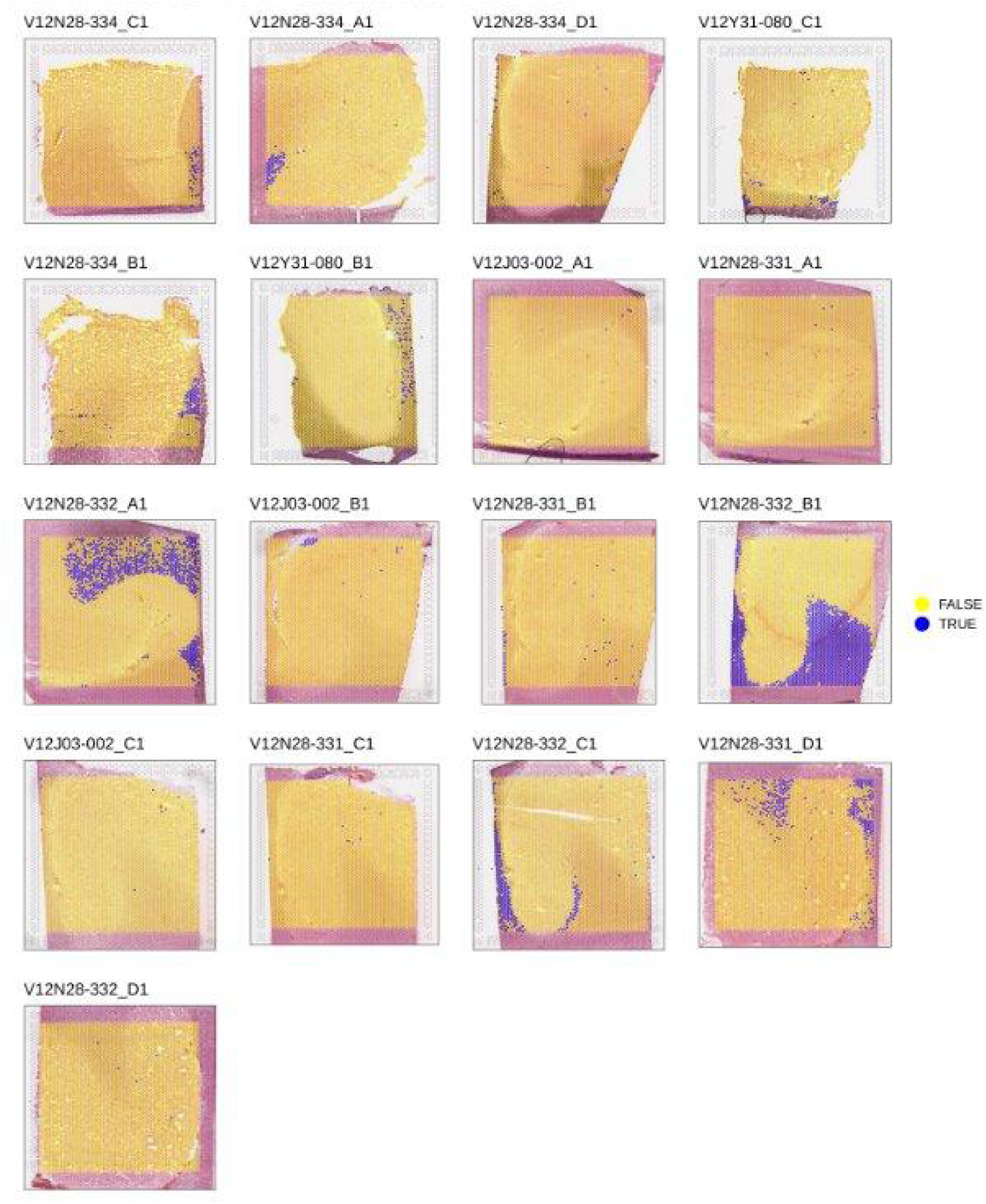
Spot plots per sample showing 3x median-absolute-deviation (MAD) threshold for spots with low library size and/or low number of detected genes. The spot plot is overlaid on the histology image. Blue color represents spots that are 3 MADs below the median library size and/or median number of detected genes for each sample. If the 3 MADs criteria was used to remove spots, then these spots would have been removed. As described in **Methods**, the criteria from **Supplementary Fig. 4** was used to remove spots instead.

**Supplementary Fig. 4.**
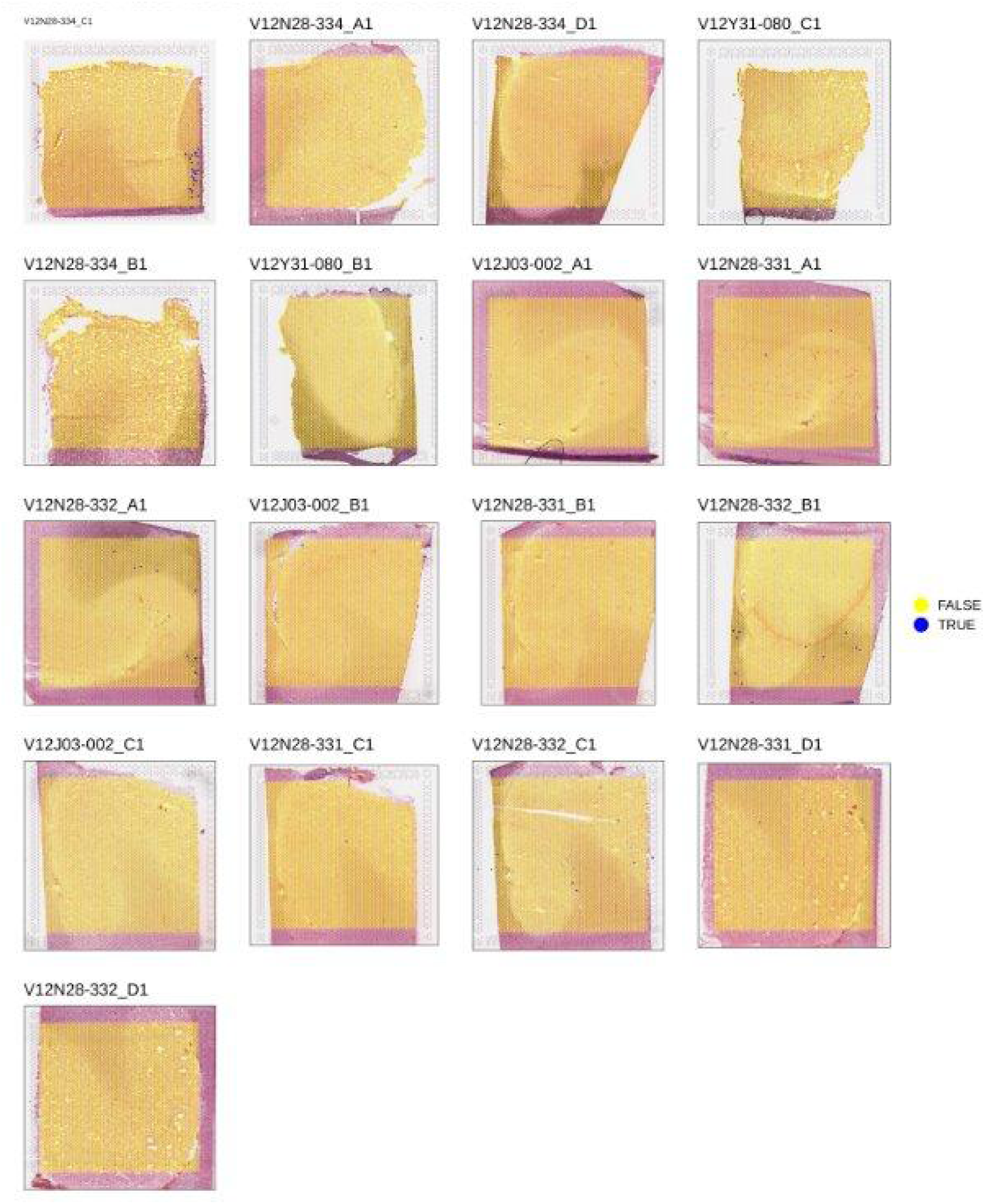
Spot plots per sample showing extreme filter for spots with library size less than 20 and/or number of detected genes less than 20. The spot plot is overlaid on the histology image. Blue color represents spots that have library size less than 20 and/or number of detected genes less than 20; these spots were removed before proceeding with downstream analyses.

**Supplementary Fig. 5.**
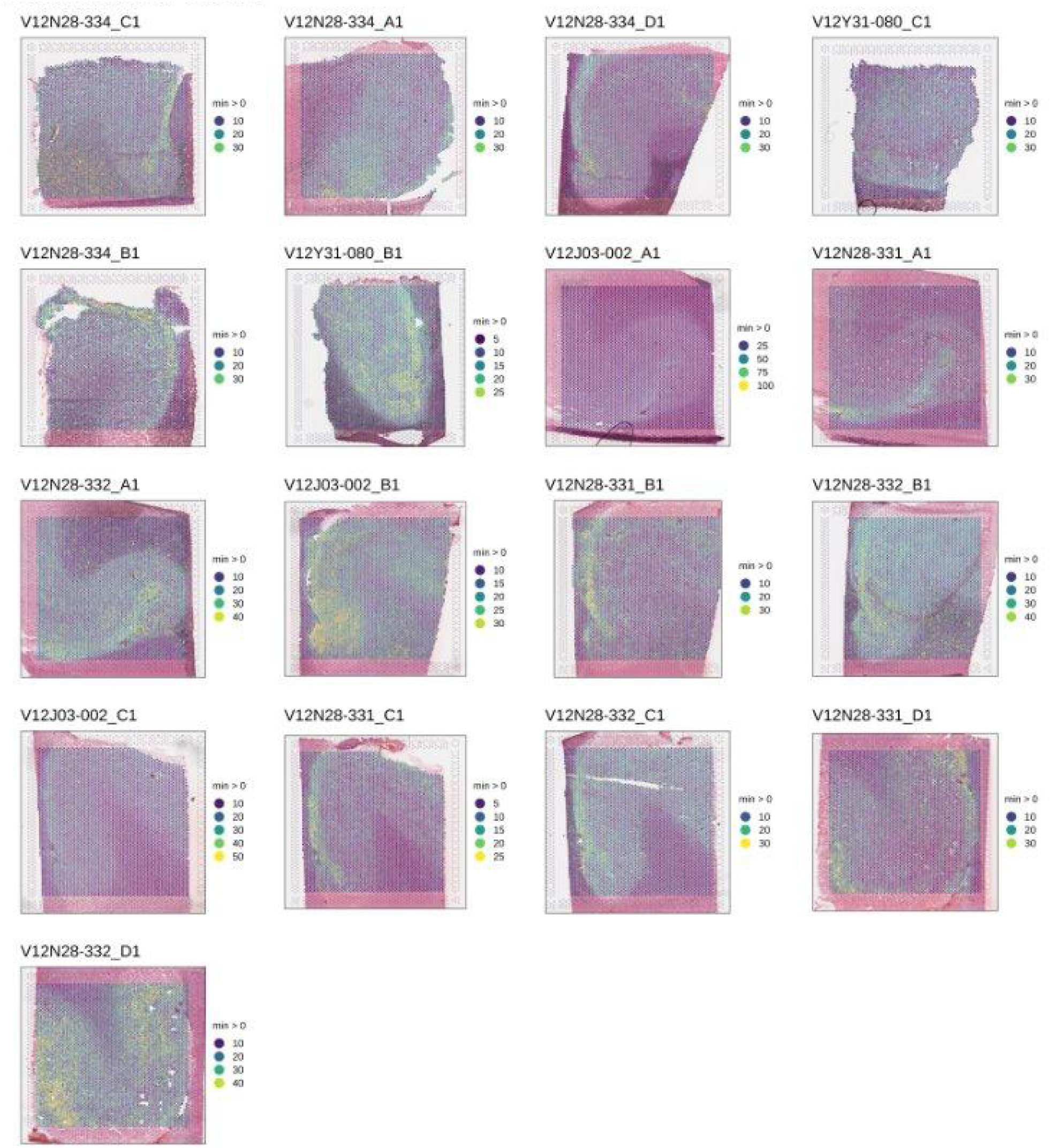
Spot plots per sample showing mitochondrial percentage for each spot. The spot plot is overlaid on the histology image. Color represents the mitochondrial percentage for each spot.

**Supplementary Fig. 6.**
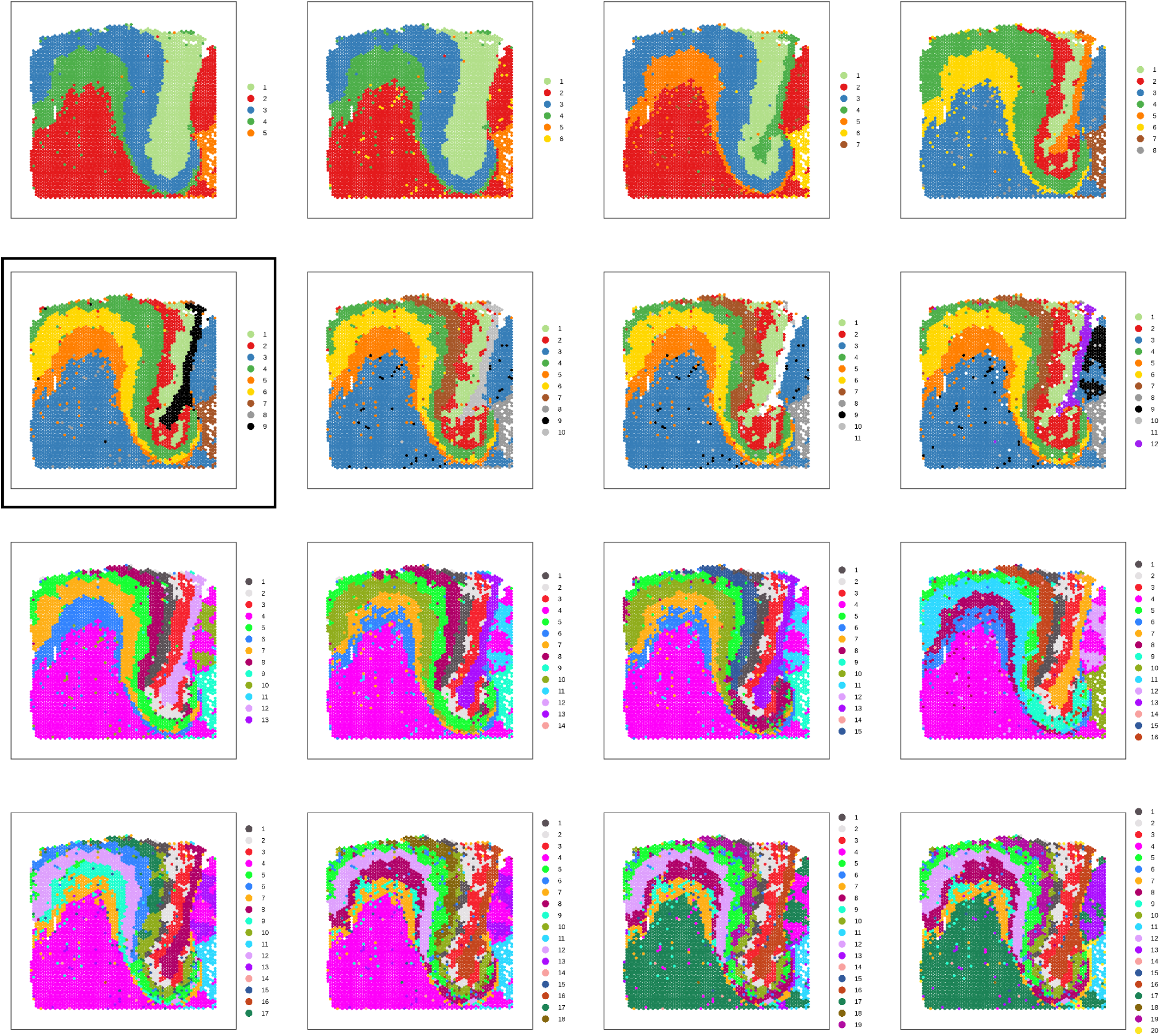
Spot plots of data-driven PRECAST clusters. Shown on one sample of the dACC SRT data (V12N28-334_C1), the number of clusters increases from 5 to 20 clusters with the descending rows. The `PRECAST` algorithm, guided by `nnSVG` spatially variable genes, was fit with the given number of clusters. Color represents the cluster label of each spot for the given `PRECAST` clustering algorithm. Note that the colors do not necessarily correspond to the same domain across clustering resolutions. The black box highlights the final clusters (*k*=9) selected for downstream analyses.

**Supplementary Fig. 7.**
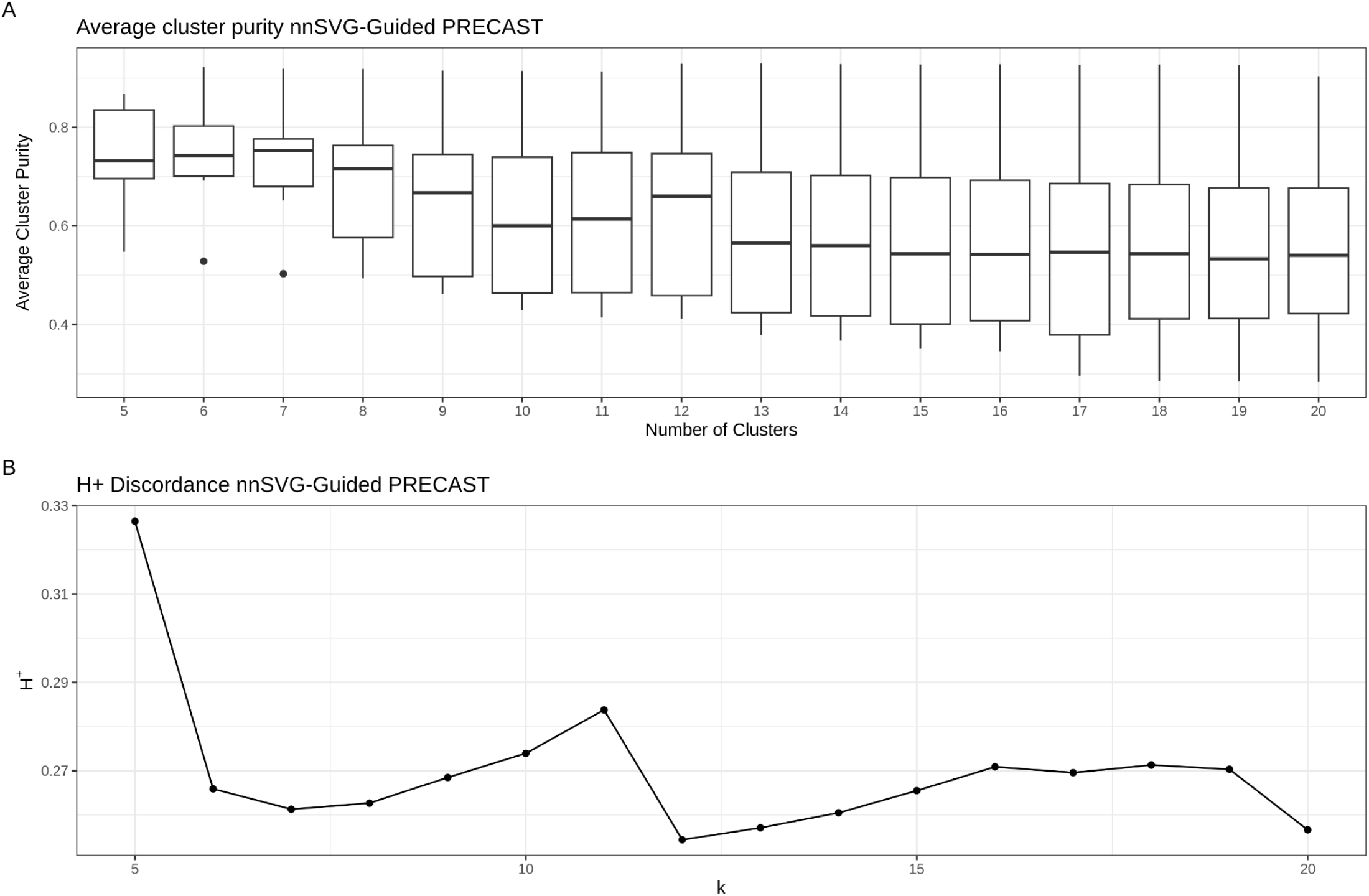
Metrics to evaluate nnSVG-guided PRECAST clusters of the dACC SRT data. (**A**) Boxplots of average cluster purity (*y*-axis) for each clustering algorithm from `PRECAST` with number of clusters ranging from 5 to 20 (*x*-axis). The purity of each cluster per gene is the proportion of observations in its neighborhood from a different cluster, and the average purity is the average of the purity scores across genes in a specific cluster. Higher cluster purity is desirable. (**B**) Line plot of the H_+_ discordance metric (*y*-axis) (computed with `fasthplus`) for each clustering algorithm from `PRECAST` with the number of clusters ranging from *k*=5 to 20 (*x*-axis). This metric measures the discordance of unsupervised clusters, and lower values are desirable.

**Supplementary Fig. 8.**
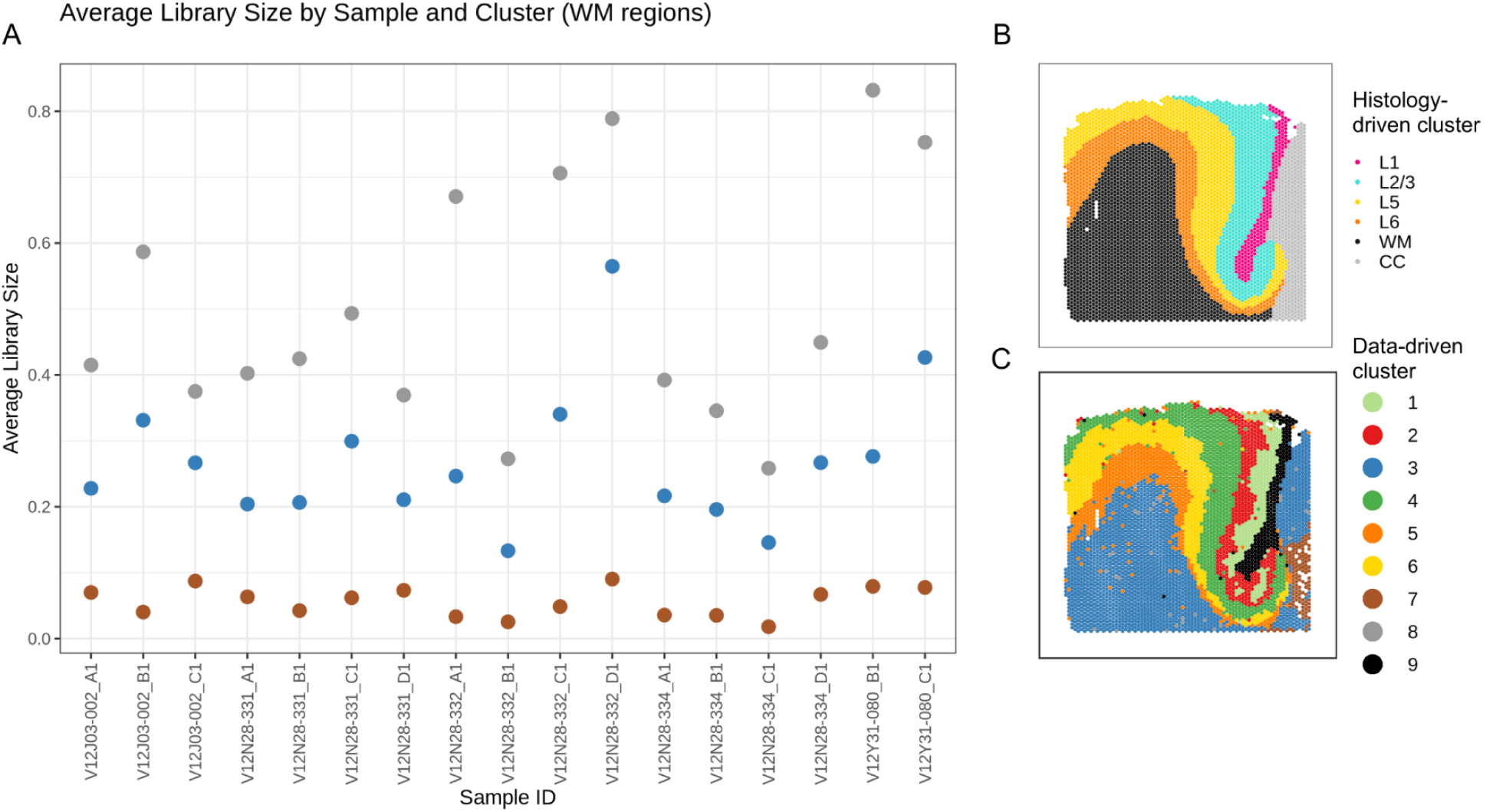
Average library size of white matter nnSVG-guided PRECAST clusters of the dACC SRT data. Dot plot of average library size (*y*-axis) per sample (*x*-axis) within `PRECAST` clustering algorithm *k*=9. Color represents cluster number, matching with *k*=9 in **Supplementary Fig. 6** (**C**), with an example from sample V12N28-334_C1. (**B**) Domains from sample V12N28-334_C1. Color represents the histology-driven manually annotated spatial domain of each spot.

**Supplementary Fig. 9.**
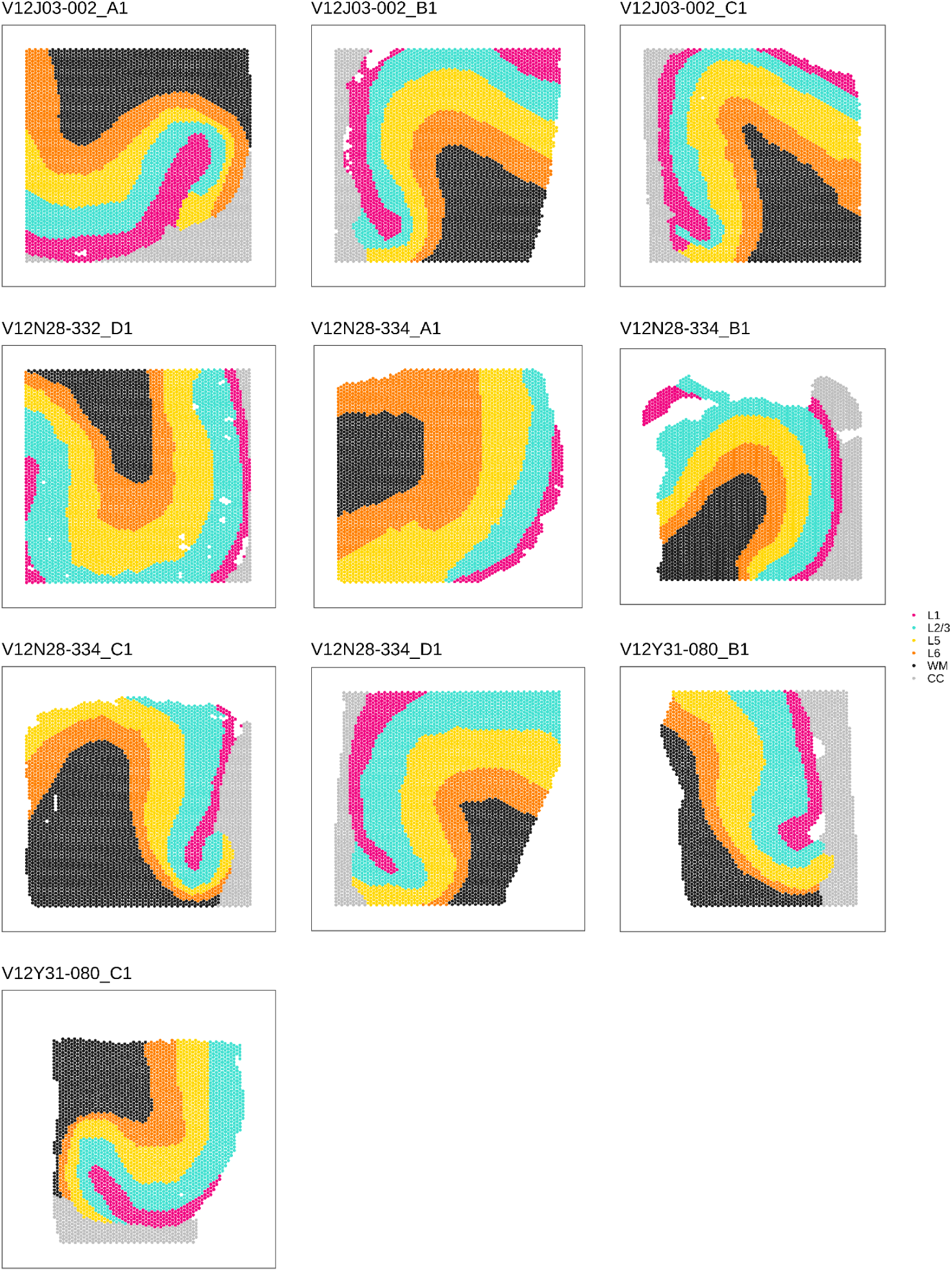
Spot plots showing histology-driven manual annotations in dACC SRT data. For ten samples of the dACC SRT data, one sample for each donor was manually annotated with `Samui Browser` (Sriworarat et al. 2023) based on anatomical features and laminar marker genes. Color represents the histology-driven manually annotated spatial domain of each spot.

**Supplementary Fig. 10.**
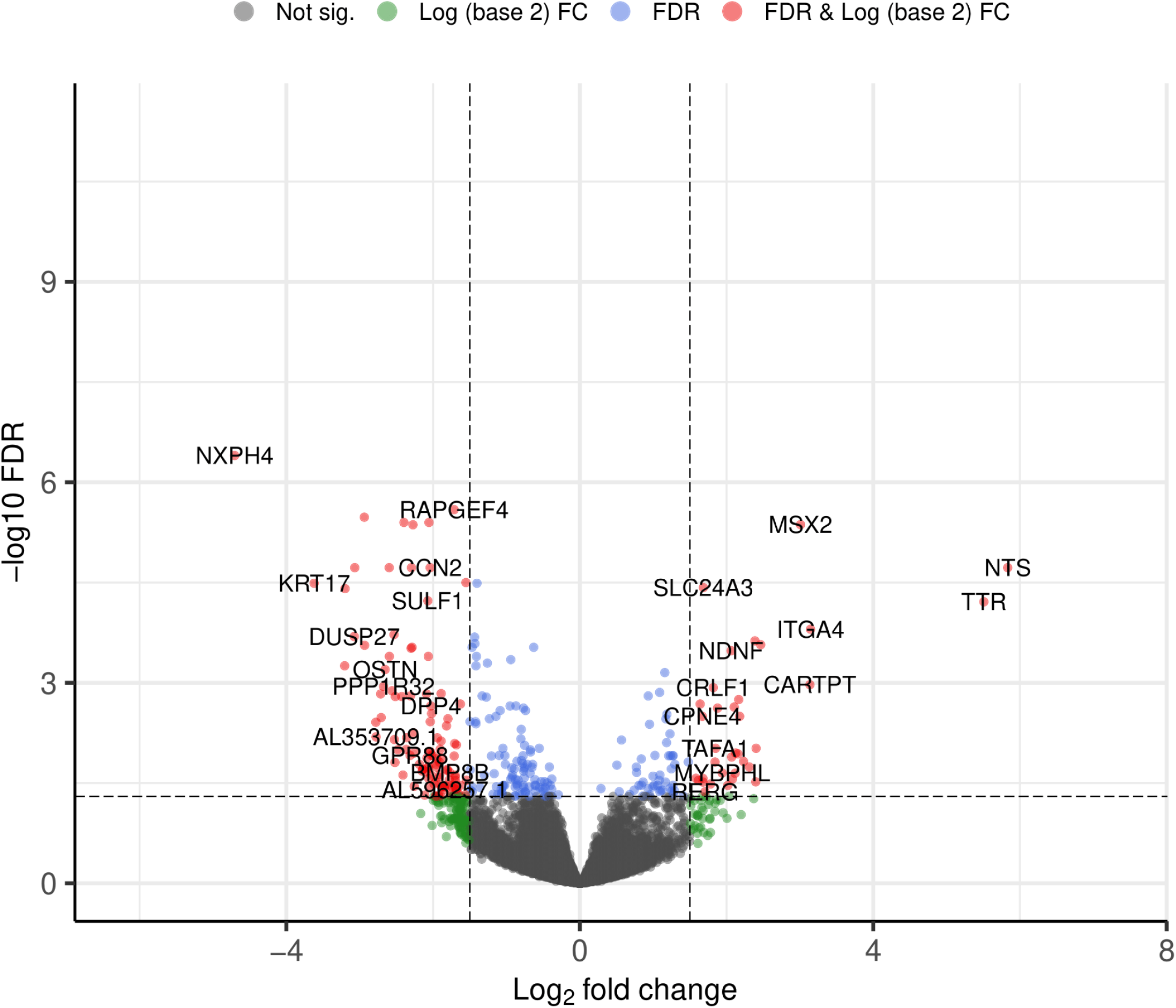
Differential expression of white matter domains from histology-driven manual annotations in dACC SRT data. `EnhancedVolcano` plot shows the DE results for the pairwise model pseudobulked test for the CC (corpus callosum) dACC SRT manual annotation compared to the WM (white matter) dACC SRT manual annotation. Each point is a gene with its log fold-change (logFC) (*x*-axis) and statistical significance (*y*-axis). Statistical significance is measured with negative log-transformation of FDR-adjusted *p*-values. Color indicates categorization of each gene; red represents statistically significant with FDR < 0.05 and absolute value of logFC > 1, blue represents statistically significant with FDR < 0.05 only, grey represents not statistically significant with FDR >= 0.05, and green represents absolute value of logFC > 1 only.

**Supplementary Fig. 11.**
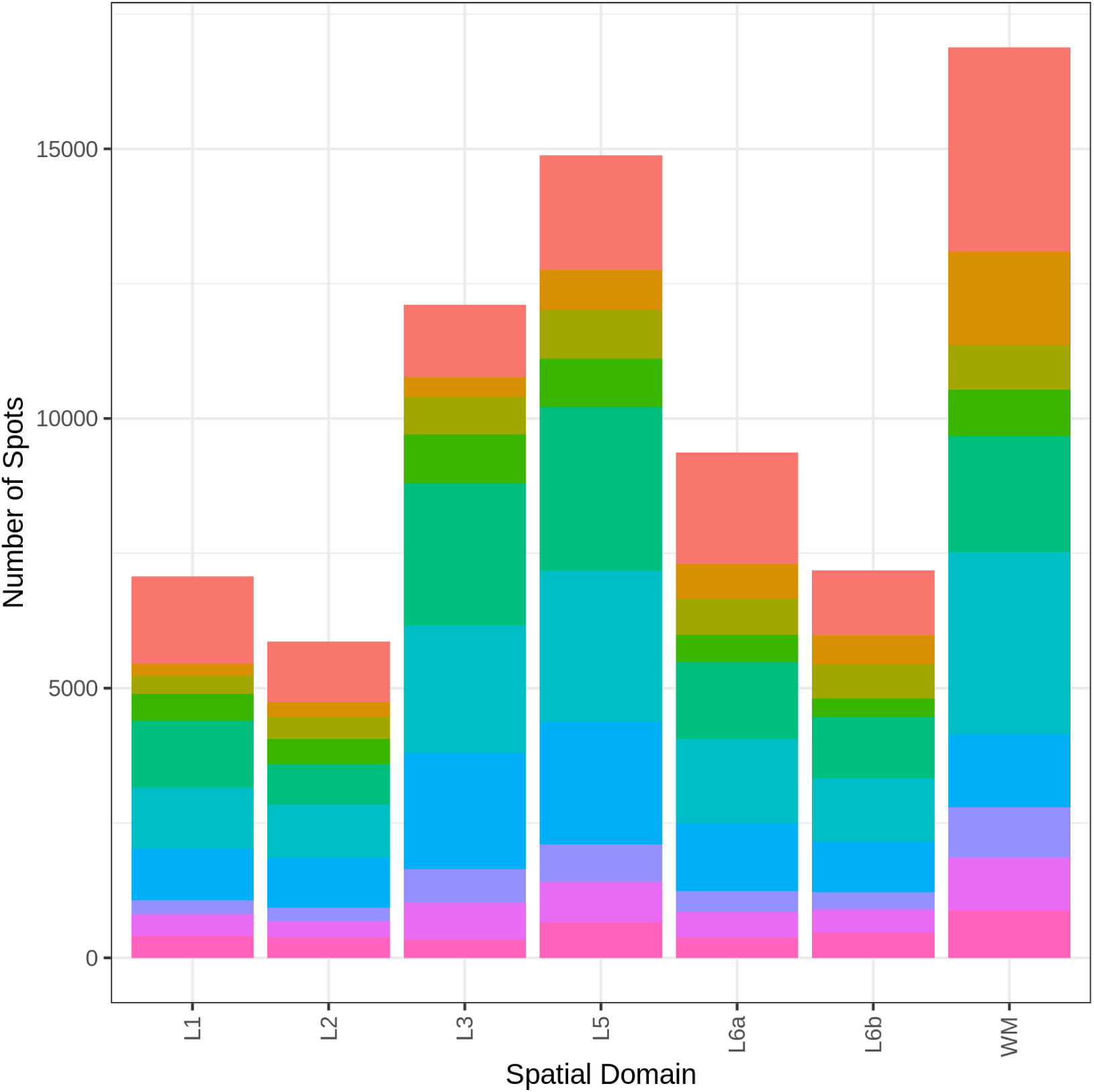
Brain donor composition of dACC SRT spatial domains. Barplot displays the number of spots (*y*-axis) in each spatial domain (*x*-axis) in the dACC SRT data. Color represents the proportion of spots in each spatial domain coming from each donor.

**Supplementary Fig. 12.**
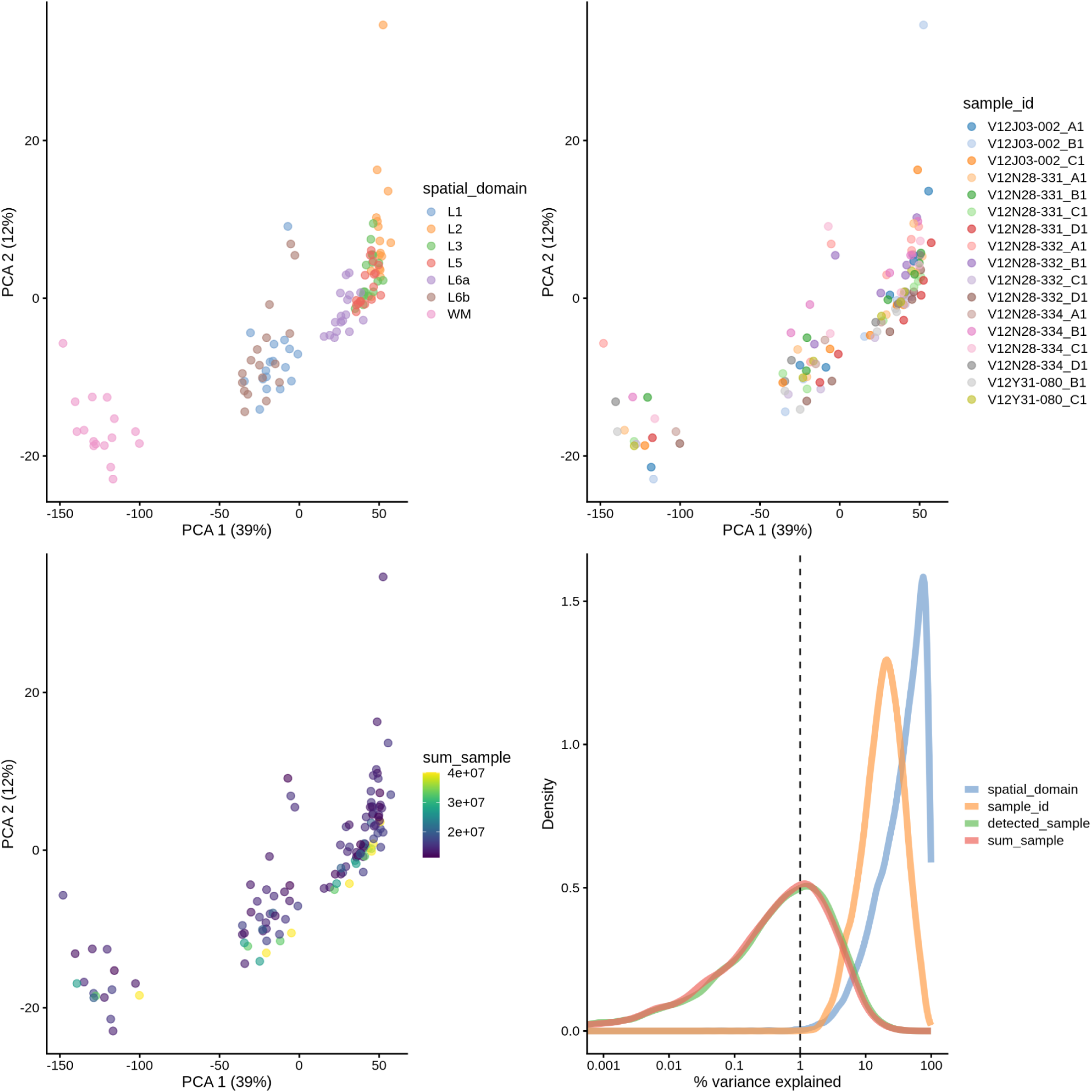
Principal components analysis (PCA) of pseudobulked dACC SRT data. The first three plots show the score plots of the first two (PCs) of the pseudobulked dACC SRT spatial domains. Each score plot is colored by spatial domain, sample id, and total UMI counts per sample, respectively. The fourth plot, made with scater (McCarthy et al. 2017), shows the percent variance explained by spatial domain, sample id, total UMI counts per sample, and total detected genes per sample.

**Supplementary Fig. 13.**
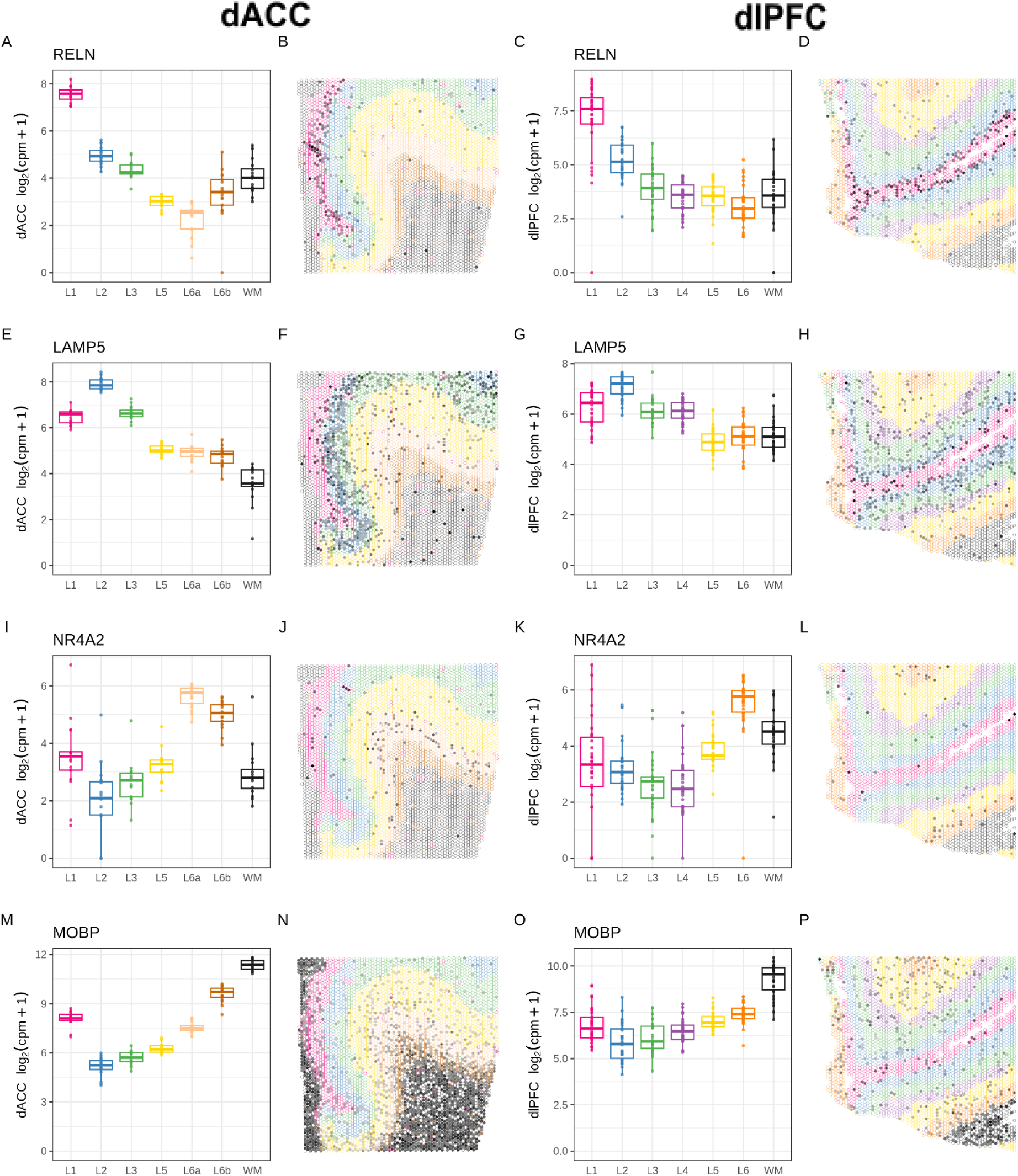
Spatial domain layer markers in paired dACC and dlPFC SRT data. Each row shows information for one gene, in order, *RELN, LAMP5, NR4A2, MOBP.* Each column displays a similar style of plot. First column (**A, E, I, M**): The *y*-axis displays log_2_(counts per million + 1) expression (computed manually) for each spatial domain (*x*-axis) in the pseudobulked dACC SRT data. Color represents the spatial domain. Second column (**B, F, J, N**): `escheR` spot plot of dACC Visium capture area from donor Br6432 (sample ID: V12N28-331_B1) with spots colored by the dACC spatial domains. Fill represents log_2_-normalized expression per spot. Third column (**C, G, K, O**): The *y*-axis displays log_2_(counts per million + 1) expression (computed manually) for each spatial domain (*x*-axis) in the pseudobulked dlPFC SRT data. Color represents the spatial domain. Fourth column (**D, H, L, P**): Spot plot of dlPFC Visium capture area from donor Br6432 (sample ID: Br6432_ant) with spots colored by the dlPFC spatial domains. Fill represents log_2_-normalized expression per spot.

**Supplementary Fig. 14.**
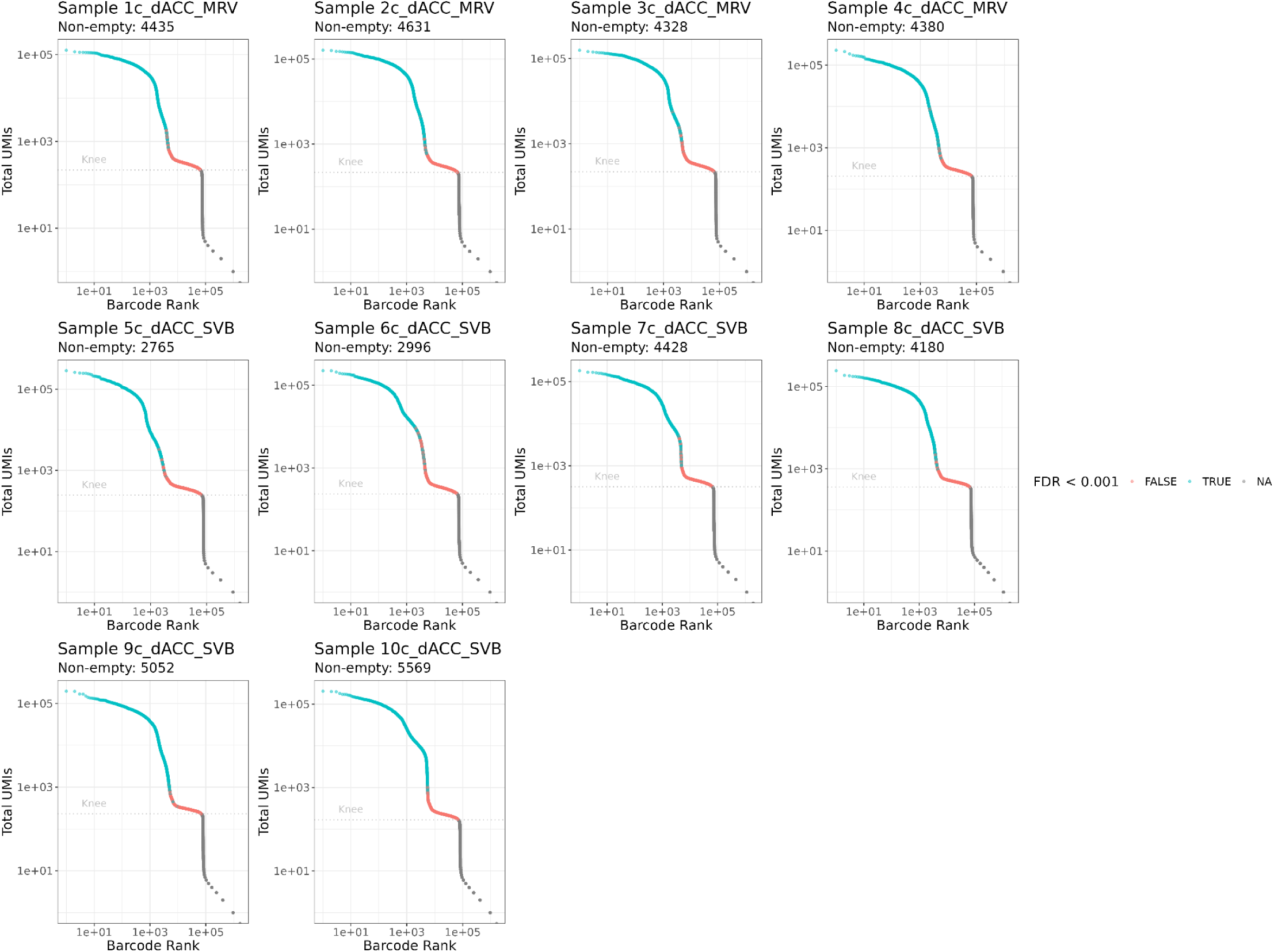
Empty droplets preprocessing of dACC snRNA-seq data. `DropletUtils` knee plots identify sample-specific UMI thresholds for empty droplets preprocessing in the dACC snRNA-seq data. Color represents quality based on an FDR cutoff of 0.001: blue indicates predicted real nuclei with FDR < 0.001 and red indicates predicted empty nuclei with FDR > 0.001. The horizontal dotted line represents the knee point used as an empty droplet cutoff. Each knee plot represents a different sample.

**Supplementary Fig. 15.**
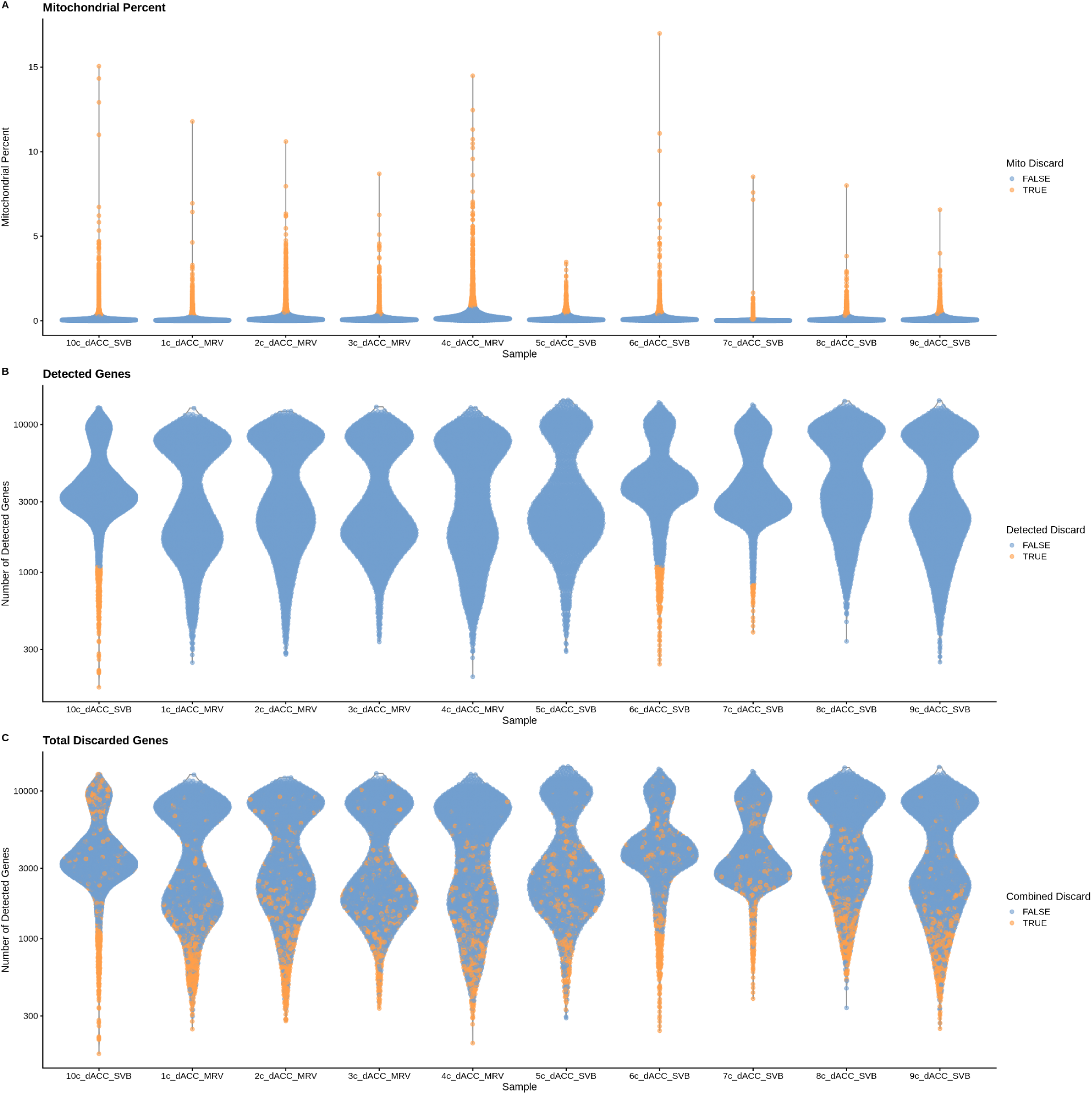
Quality control (QC) metrics for dACC snRNA-seq data. Color represents discarded status: yellow indicates nuclei that were dropped, while blue indicates nuclei used for downstream analyses. (**A**) Violin plot shows the mitochondrial percentage (*y*-axis) for each nucleus, separately plotted for each sample (*x*-axis). (**B**) Violin plot shows the number of detected genes (*y*-axis) for each nucleus, separately plotted for each sample (*x*-axis). (**C**) Violin plot shows the number of detected genes (*y*-axis) for each nucleus, separately plotted for each sample (*x*-axis). The color represents nuclei discarded based on mitochondrial percentage and number of detected genes. Thresholds were determined with scuttle (McCarthy et al. 2017).

**Supplementary Fig. 16.**
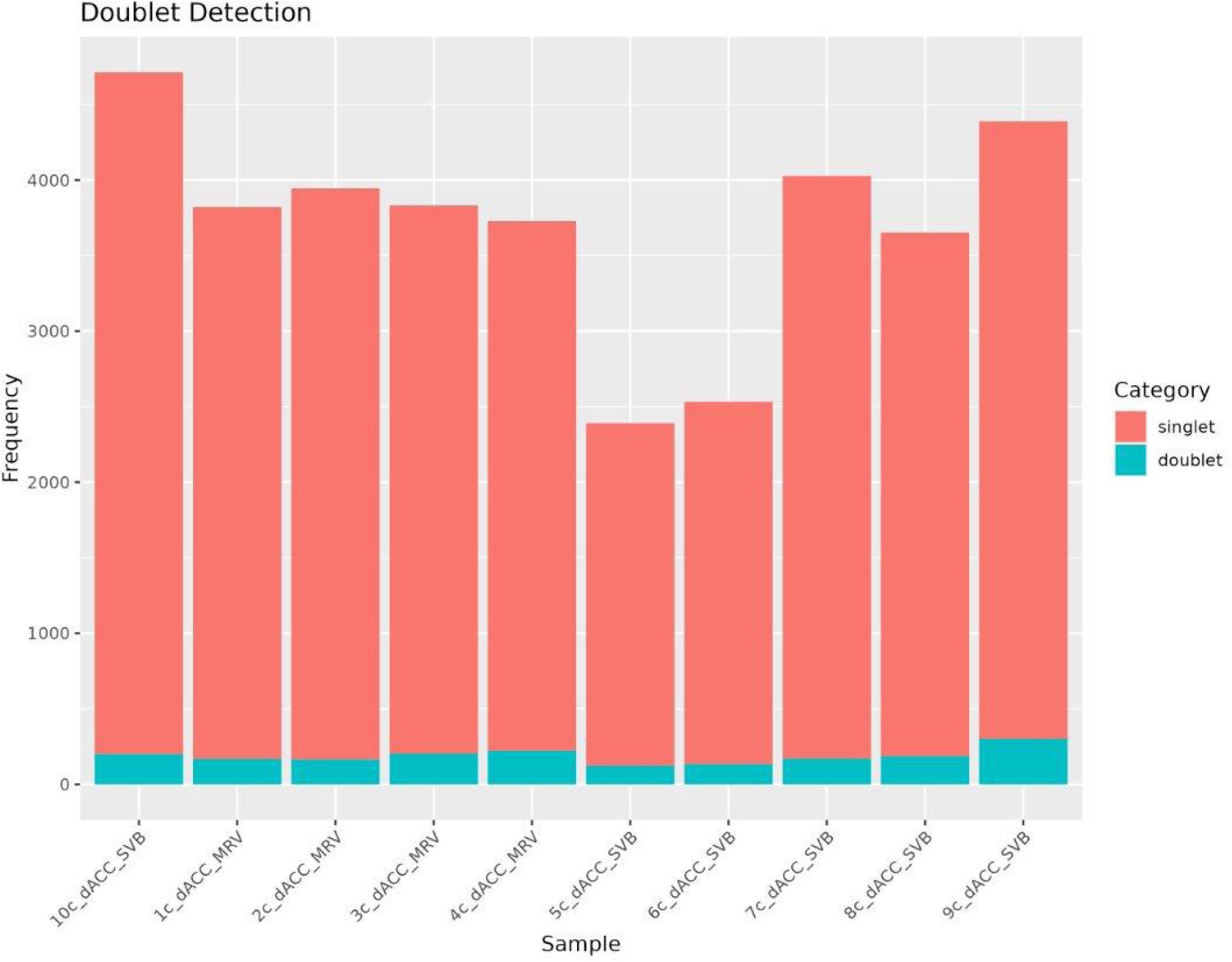
Doublet detection removes a small proportion of nuclei in dACC snRNA-seq data. Barplots quantify the frequency (*y*-axis) of doublets and singlets in each sample (*x*-axis) of the dACC snRNA-seq data. Color represents the proportion of singlets and doublets in each sample. Doublets were detected with scDblFinder (Germain et al. 2021).

**Supplementary Fig. 17.**
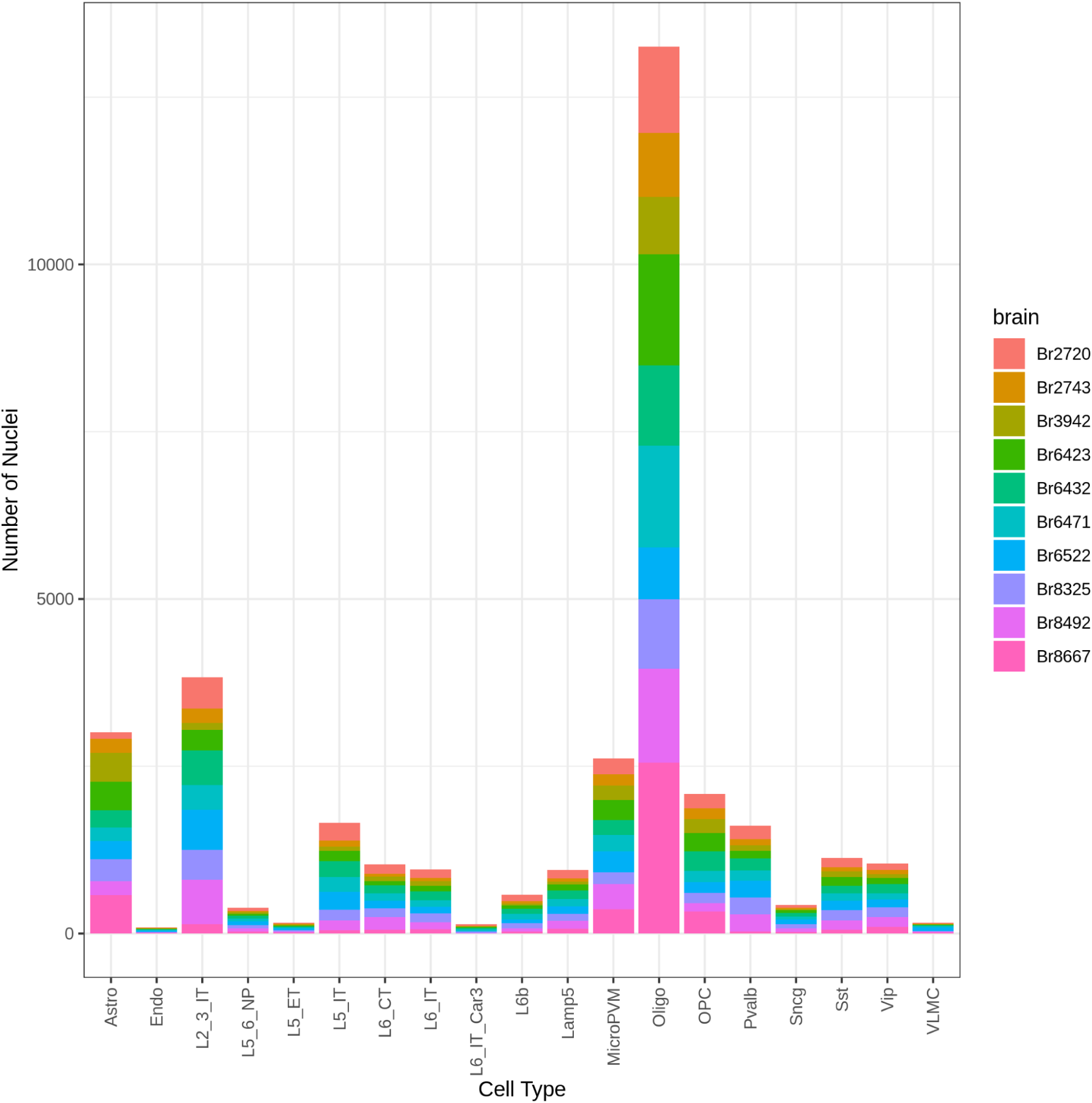
Brain donor composition of dACC snRNA-seq cell types. Barplot displays the number of nuclei (*y*-axis) in each cell type (*x*-axis) in the dACC snRNA-seq data. Color represents the proportion of nuclei in each cell type coming from each donor.

**Supplementary Fig. 18.**
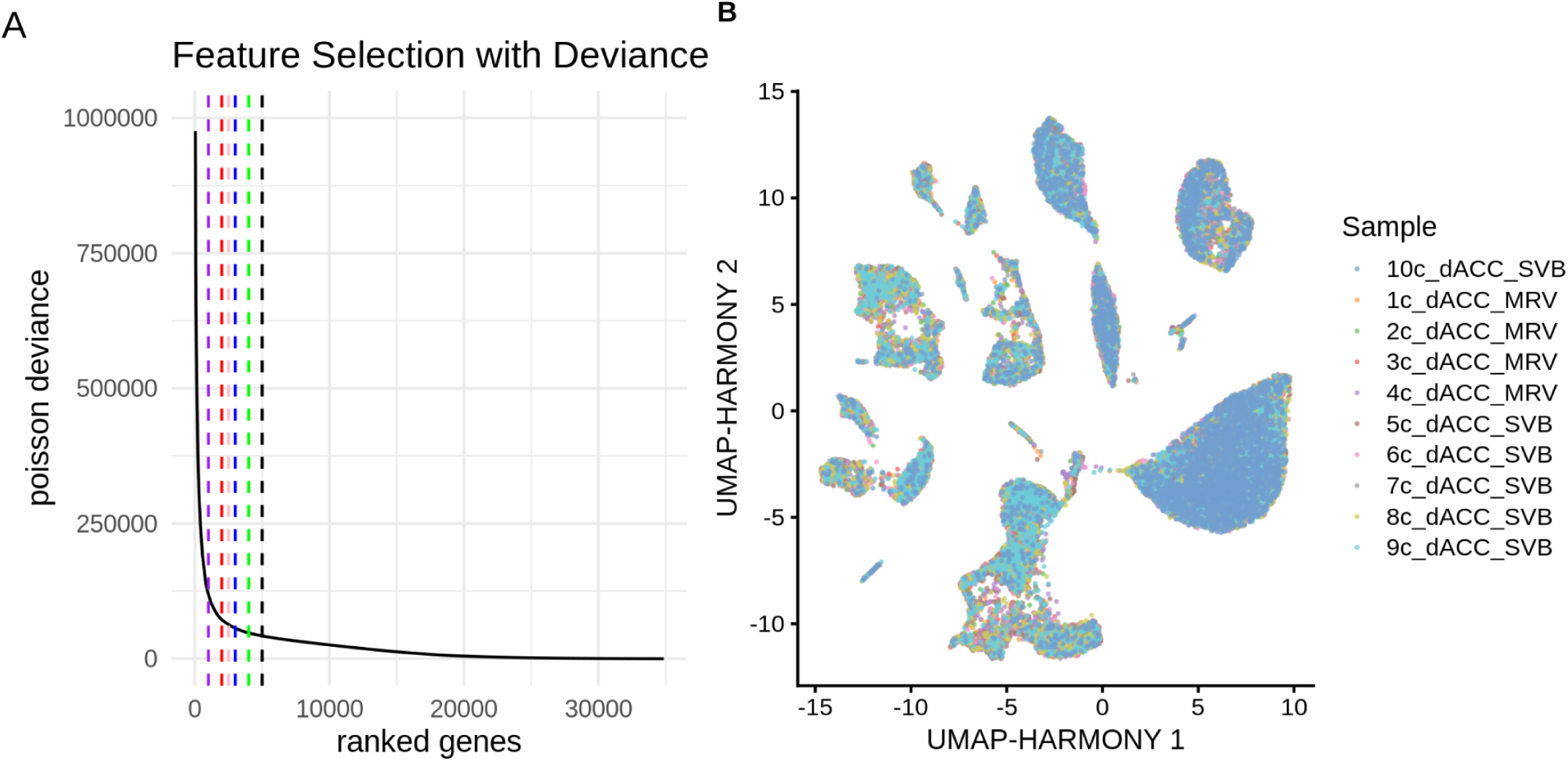
Dimension reduction of dACC snRNA-seq data. (**A**) The line plot shows a deviance feature selection intermediate figure. The Poisson deviance (*y*-axis) was calculated using the dACC snRNA-seq counts matrix and is plotted against the number of ranked genes (*x*-axis). The dashed lines mark 1,000, 2,000, 2,500, 3,000, 4,000, and 5,000 genes. (**B**) Uniform manifold approximation and projection (UMAP) representation of the dACC snRNA-seq dataset after Harmony batch correction. Color represents the sample for each nucleus.

**Supplementary Fig. 19.**
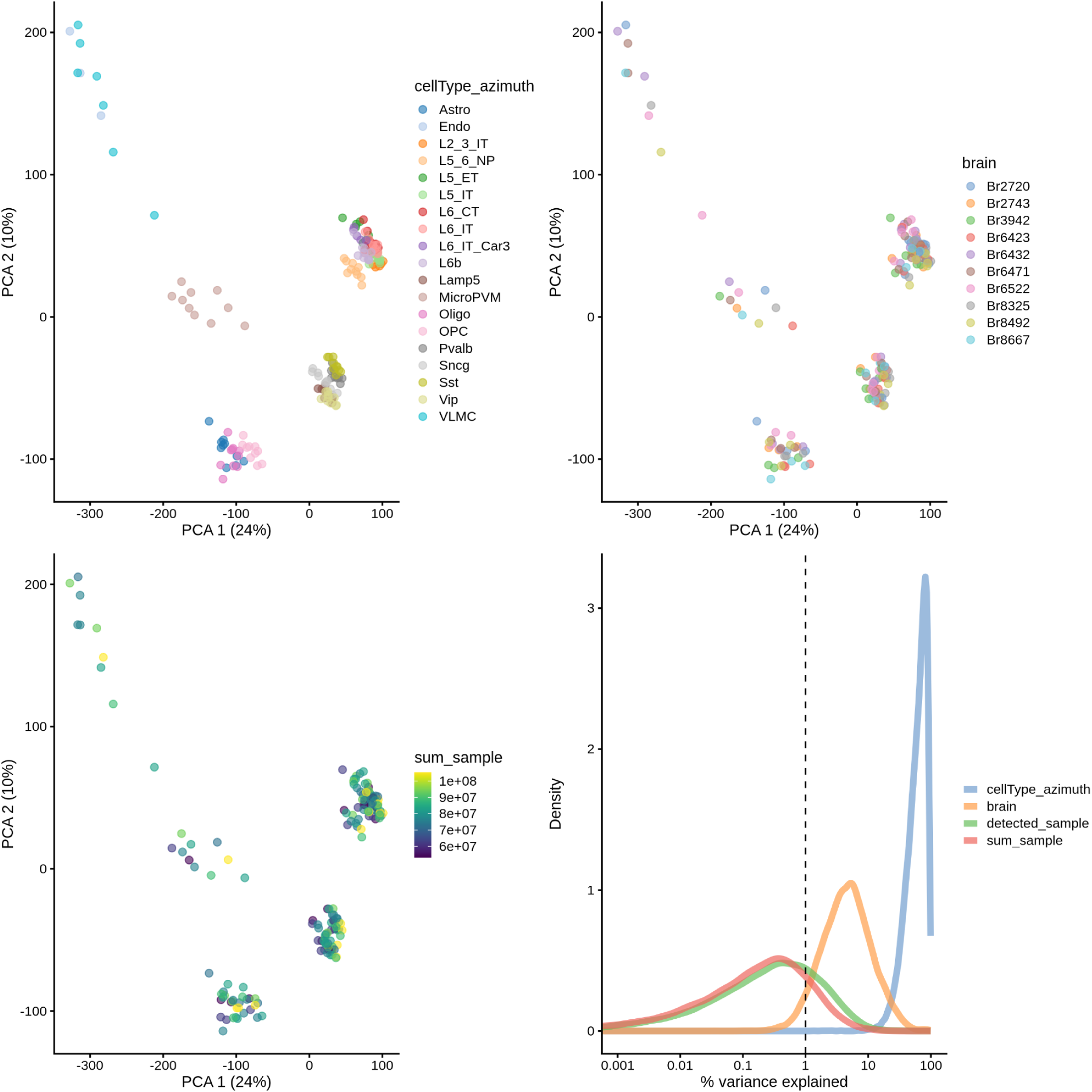
Principal components analysis (PCA) of pseudobulked dACC snRNA-seq data. The first three plots show the score plots of the first two (PCs) of the pseudobulked dACC snRNA-seq cell types. Each score plot is colored by cell type, brain donor, and total UMI counts per sample, respectively. The fourth plot, made with scater (McCarthy et al. 2017), shows the percent variance explained by cell type, brain donor, total UMI counts per sample, and total detected genes per sample.

**Supplementary Fig. 20.**
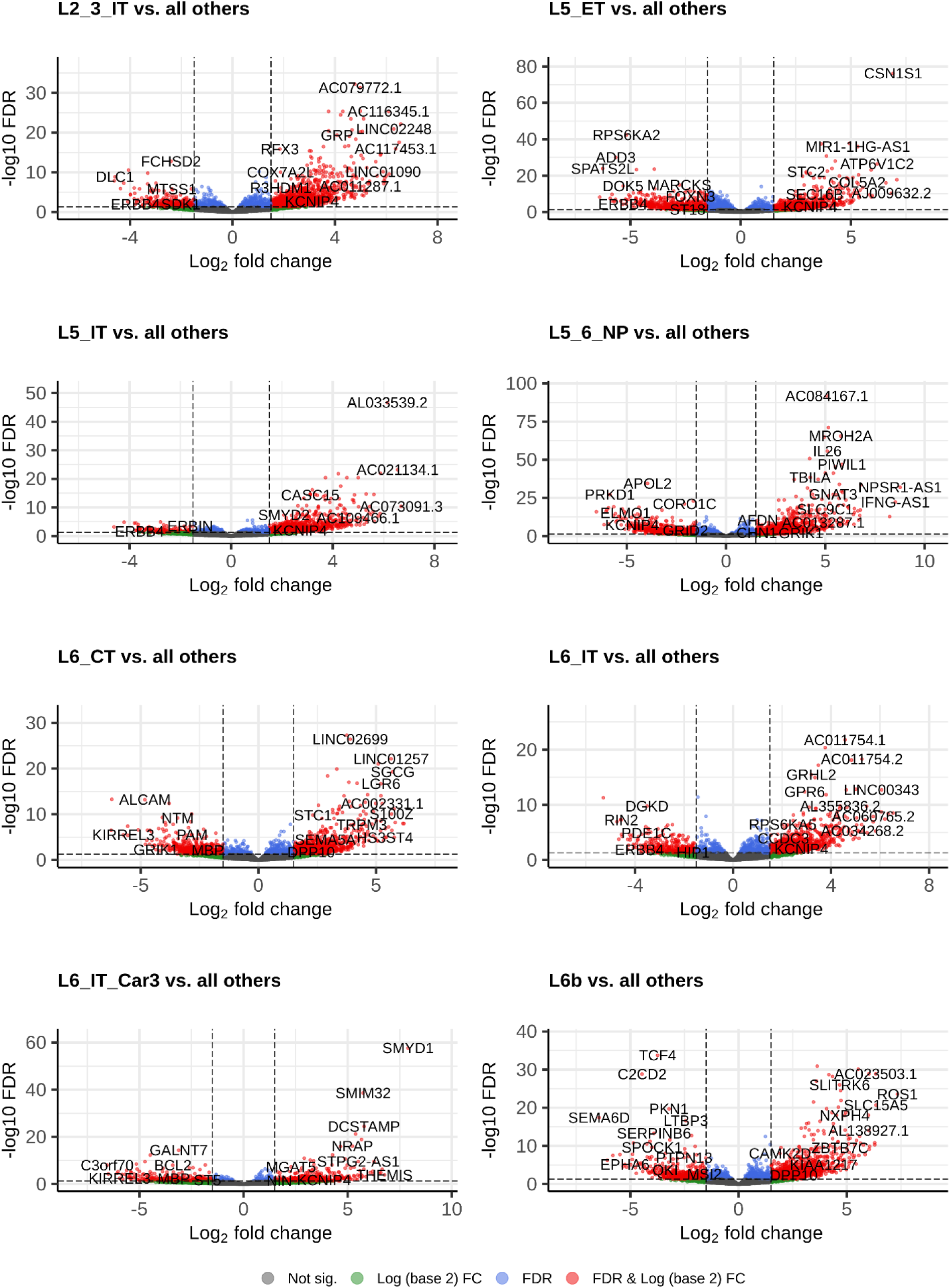
Differential expression (DE) analysis of pseudobulked dACC snRNA-seq layer-specific cell types. Each `EnhancedVolcano` plot shows the DE results for the enrichment model pseudobulked test for one dACC snRNA-seq cell type compared to all other dACC snRNA-seq cell types. Each point is a gene with its log fold-change (logFC) (*x*-axis) and statistical significance (*y*-axis). Statistical significance is measured with negative log-transformation of FDR-adjusted *p*-values. Color indicates categorization of each gene; red represents statistically significant with FDR < 0.05 and absolute value of logFC > 1, blue represents statistically significant with FDR < 0.05 only, and grey represents not statistically significant with FDR >= 0.05. These plots are a subset of only the layer-specific cell types in the dACC snRNA-seq data.

**Supplementary Fig. 21.**
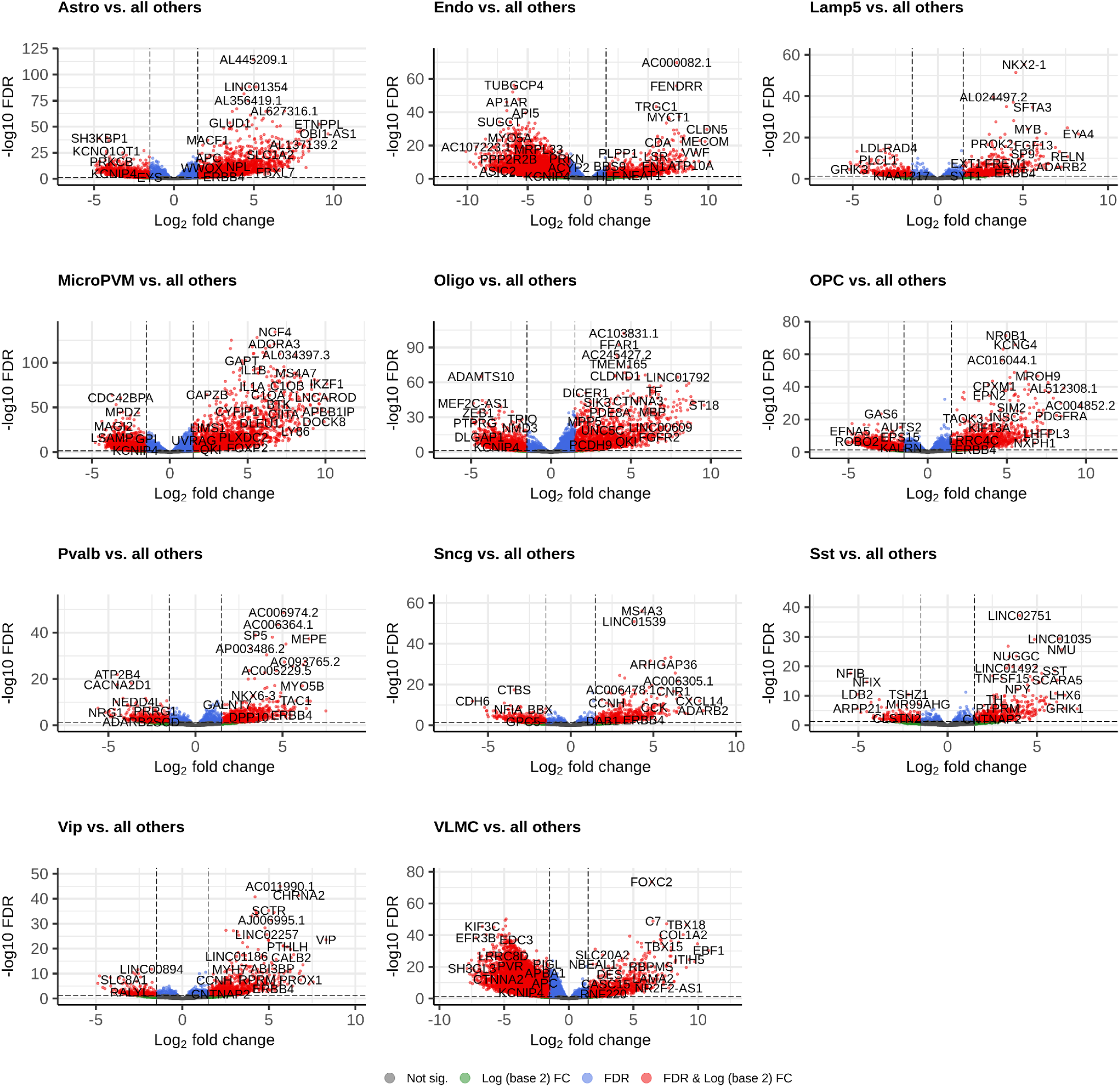
Differential expression (DE) analysis of pseudobulked dACC snRNA-seq non-layer-specific cell types. Each `EnhancedVolcano` plot shows the DE results for the enrichment model pseudobulked test for one dACC snRNA-seq cell type compared to all other dACC snRNA-seq cell types. Each point is a gene with its log fold-change (logFC) (*x*-axis) and statistical significance (*y*-axis). Statistical significance is measured with negative log-transformation of FDR-adjusted *p*-values. Color indicates categorization of each gene; red represents statistically significant with FDR < 0.05 and absolute value of logFC > 1, blue represents statistically significant with FDR < 0.05 only, and grey represents not statistically significant with FDR >= 0.05. These plots are a subset of only the non-layer-specific cell types in the dACC snRNA-seq data.

**Supplementary Fig. 22.**
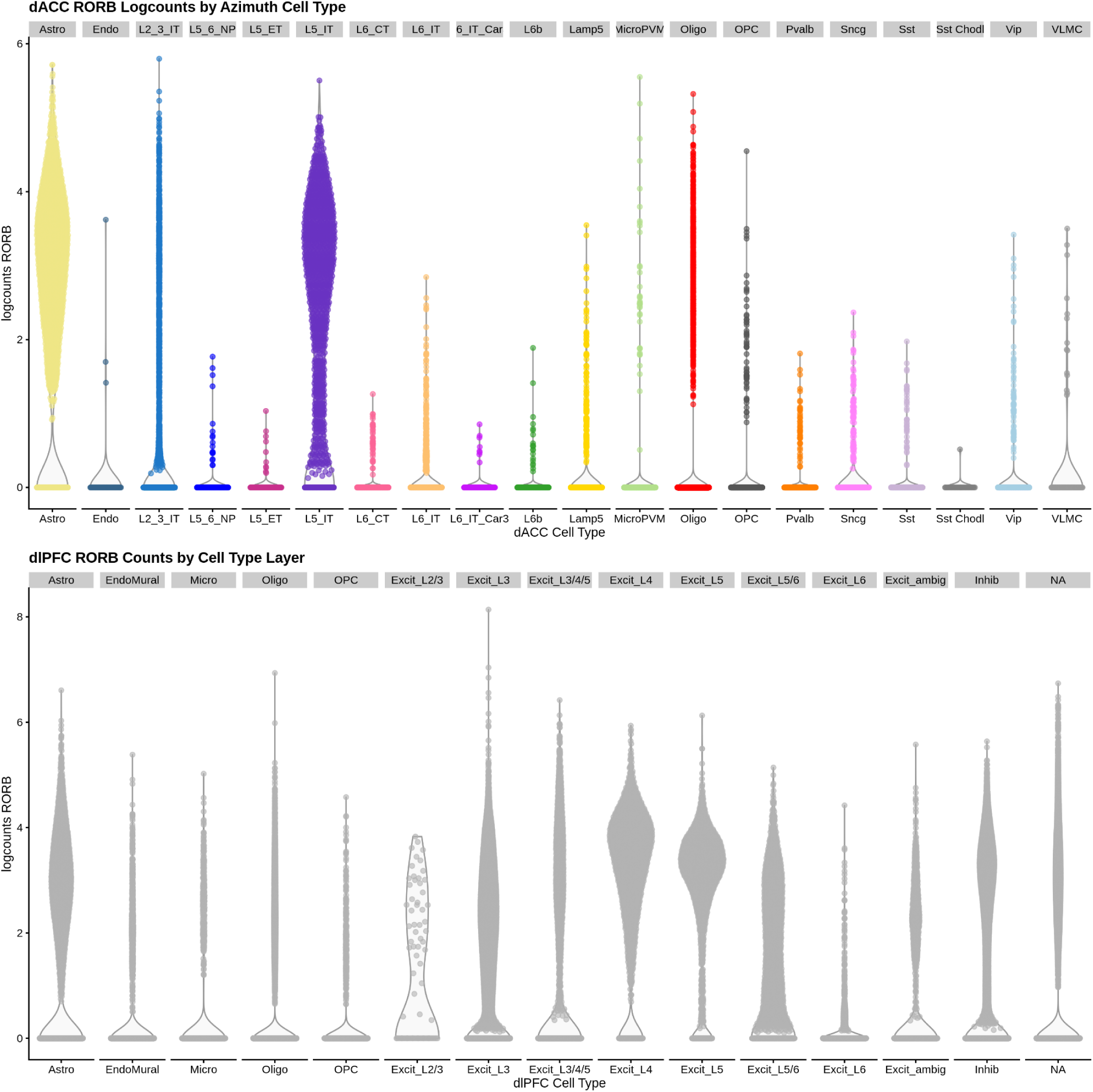
Expression of *RORB* in dlPFC and dACC snRNA-seq data. Top violin plots show log_2_-normalized expression (*y*-axis) of *RORB* across the `Azimuth` cell types (*x*-axis) in the dACC snRNA-seq data. Color represents cell type. Bottom violin plots show log_2_-normalized expression (*y*-axis) of *RORB* across the cell type layer annotations (*x*-axis) in the dlPFC snRNA-seq data.

**Supplementary Fig. 23.**
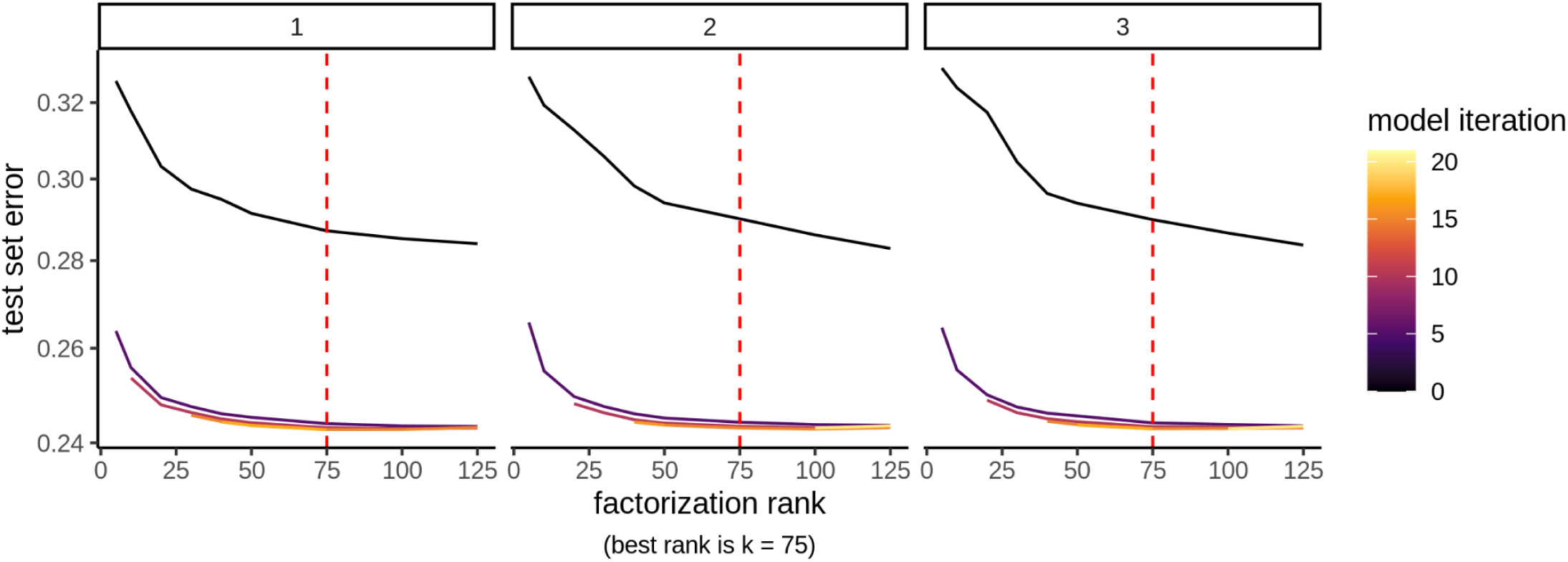
Non-negative matrix factorization (NMF) cross-validation on dACC snRNA-seq data indicates *k*=75 as optimal rank. Each line plot shows 1 replicate of NMF cross-validation on the log_2_-normalized dACC snRNA-seq data, which was computed on the ranks of 5, 10, 20, 30, 40, 50, 75, 100, and 125 (*x*-axis). The test set error is plotted on the *y*-axis. The cross-validation software `singlet` identified *k*=75 as the optimal rank to use for NMF factorization.

**Supplementary Fig. 24.**
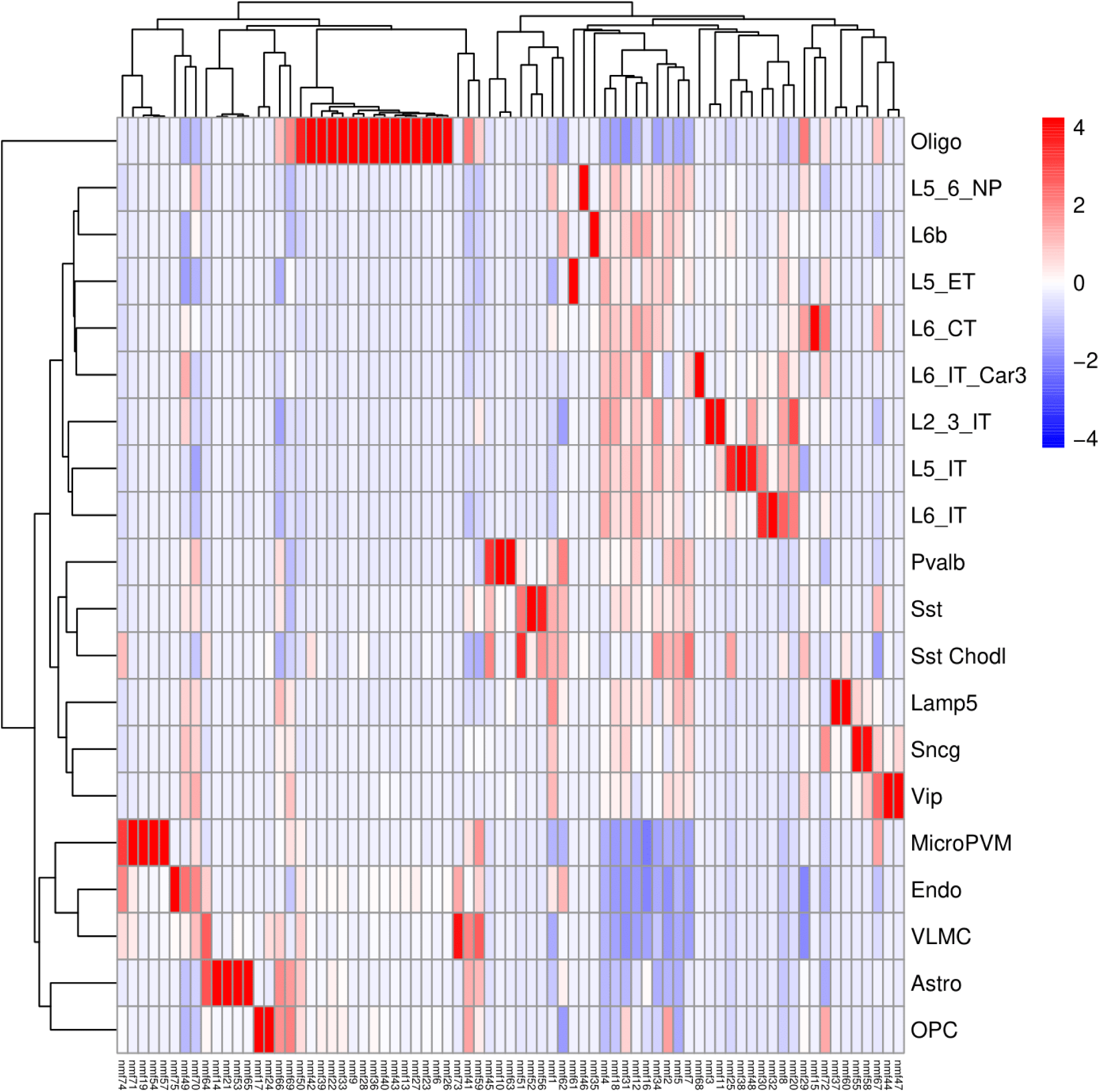
Non-negative matrix factorization (NMF) patterns of dACC snRNA-seq data associated with snRNA-seq cell types. Heatmap displays the correlation between the 75 NMF patterns from the dACC snRNA-seq data (*y*-axis) and snRNA-seq cell types (*x*-axis). The NMF patterns were aggregated across cell types, and the mean of each NMF pattern within each cell type is displayed.

**Supplementary Fig. 25.**
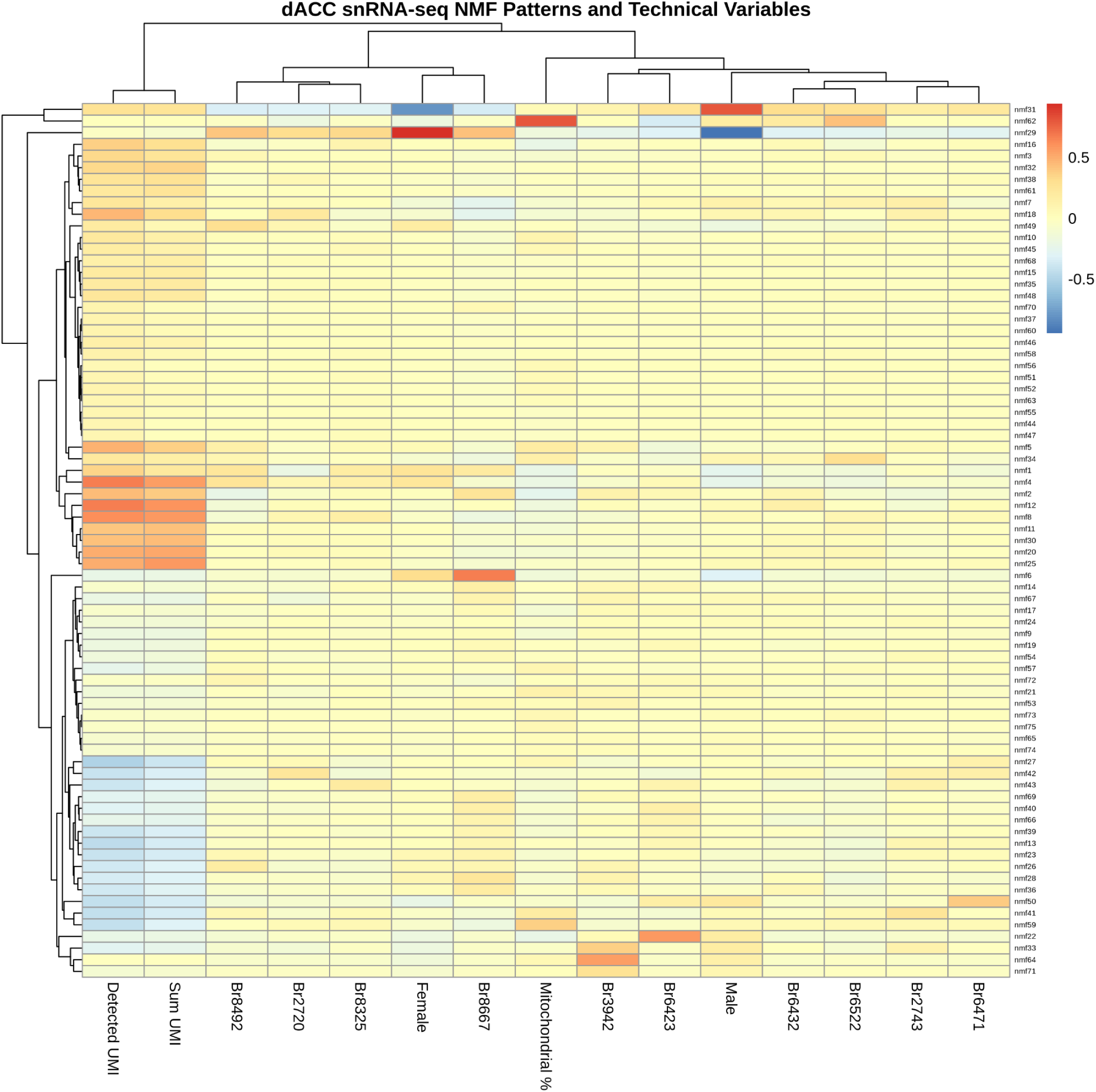
Non-negative matrix factorization (NMF) patterns of dACC snRNA-seq data associated with technical variables. Heatmap displays the correlation between the 75 NMF patterns from the dACC snRNA-seq data (*y*-axis) and technical variables related to brain donor, sex of brain donor (Male or Female), and quality control metrics (Detected UMI counts, Sum UMI counts, and Mitochondrial Percentage) (*x*-axis).

**Supplementary Fig. 26.**
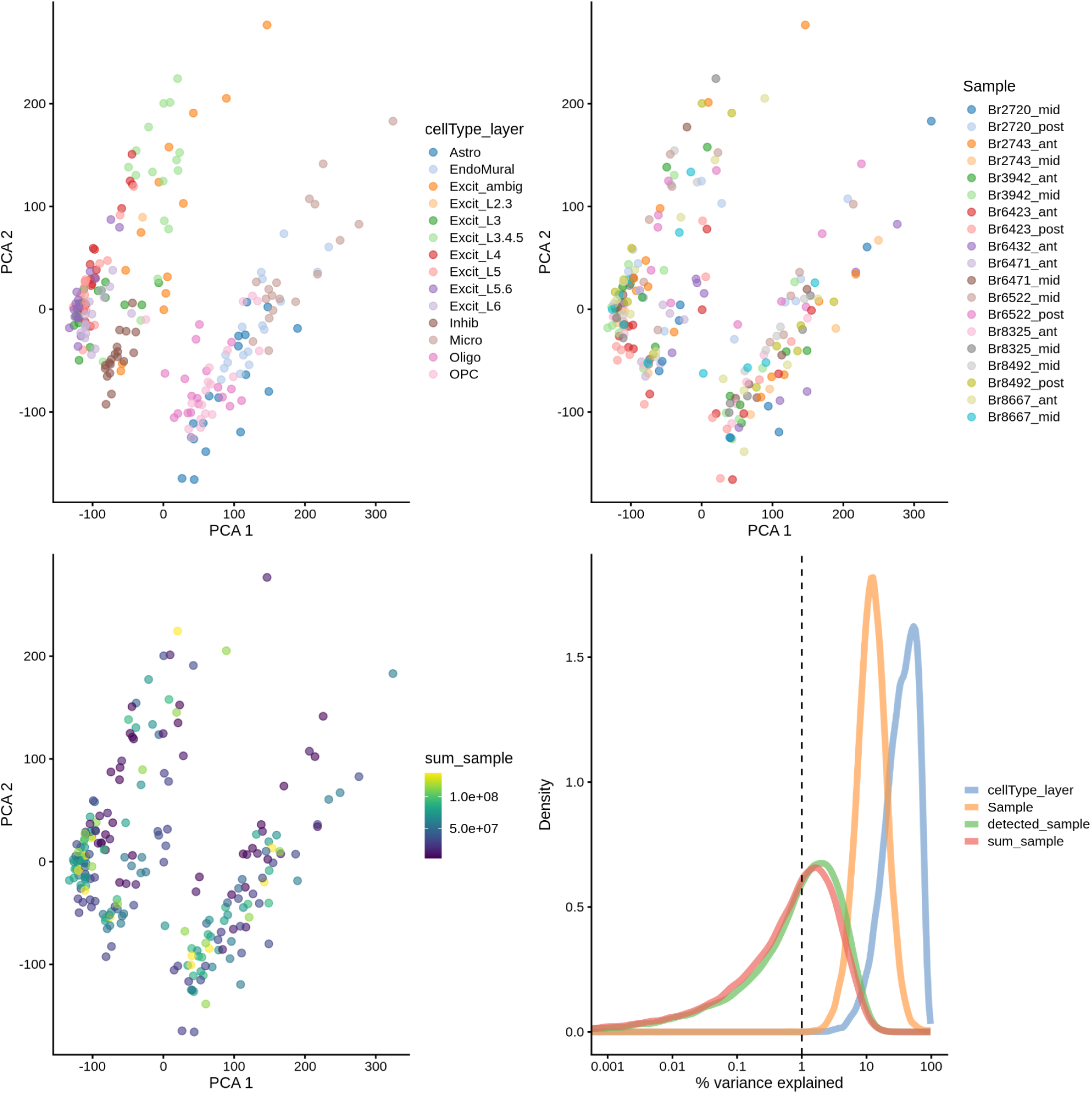
Principal components analysis (PCA) of pseudobulked dlPFC snRNA-seq data. The first three plots show the score plots of the first two (PCs) of the pseudobulked dlPFC snRNA-seq cell types. Each score plot is colored by cell type, sample id, and total UMI counts per sample, respectively. The fourth plot, made with scater (McCarthy et al. 2017), shows the percent variance explained by cell type, sample id, total UMI counts per sample, and total detected genes per sample.

**Supplementary Fig. 27.**
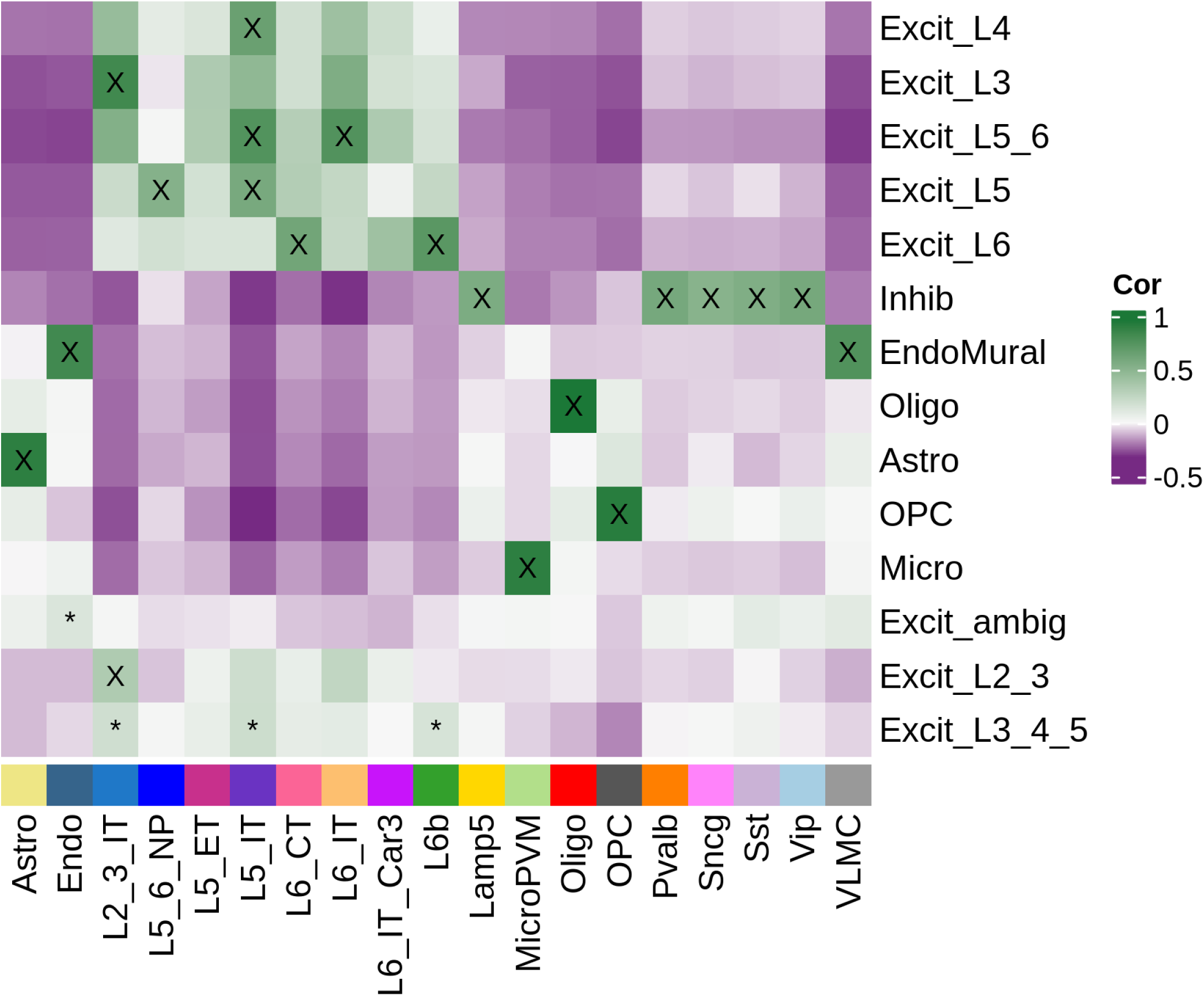
`spatialLIBD` spatial registration heatmap displays Pearson’s correlation values between the top 100 marker genes in each dlPFC snRNA-seq cell type and in each dACC snRNA-seq cell type. The *x*-axis displays the dACC snRNA-seq cell types described in (Figure 3A). The *y*-axis displays dlPFC snRNA-seq cell types: Astro: Astrocyte; EndoMural: endothelial and mural cells; Excit L2/3: Layer 2/3 excitatory neurons; Excit L3: Layer 3 excitatory neurons; Excit L3/4/5: Layer 3/4/5 excitatory neurons; Excit L4: Layer 4 excitatory neurons; Excit L5: Layer 5 excitatory neurons; Excit L5/6: Layer 5/6 excitatory neurons; Excit L6: Layer 6 excitatory neurons; Excit ambig: ambiguous excitatory neurons; Micro: microglia; Inhib: inhibitory neurons; Oligo: oligodendrocytes; OPC: oligodendrocyte precursor cell. The black “X” represents high confidence and the black asterisk represents poor confidence (`confidence_threshold` = 0.25).

**Supplementary Fig. 28.**
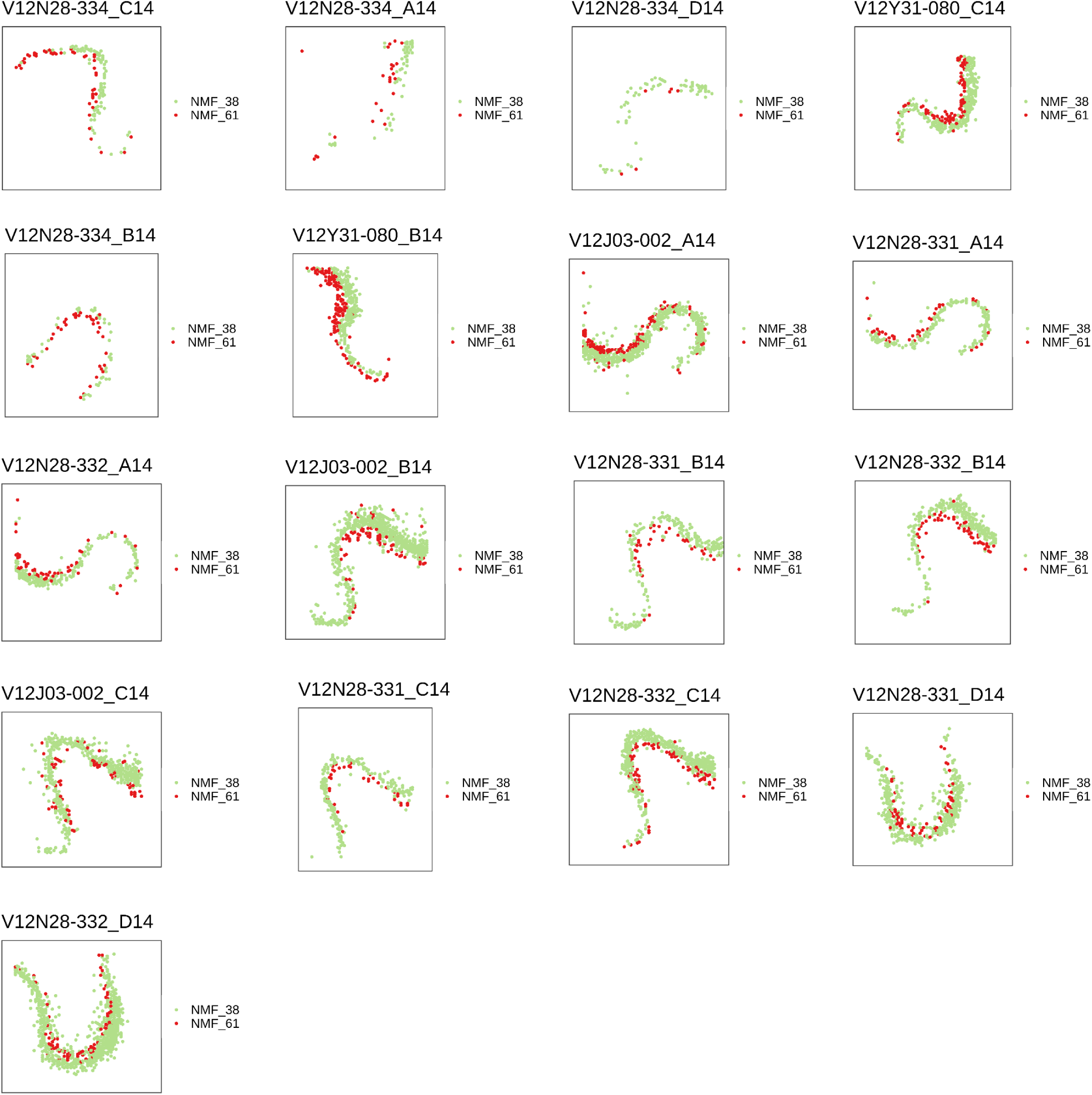
Spot plot to display NMF patterns 38 and 61 mathematically projected into the dACC SRT data. Each plot shows 1 sample of the dACC SRT data, with a subset of the spots from each sample that were predicted as either NMF38 or NMF61. Color represents spots that were predicted as either NMF38 (green) or NMF61 (red).

**Supplementary Fig. 29.**
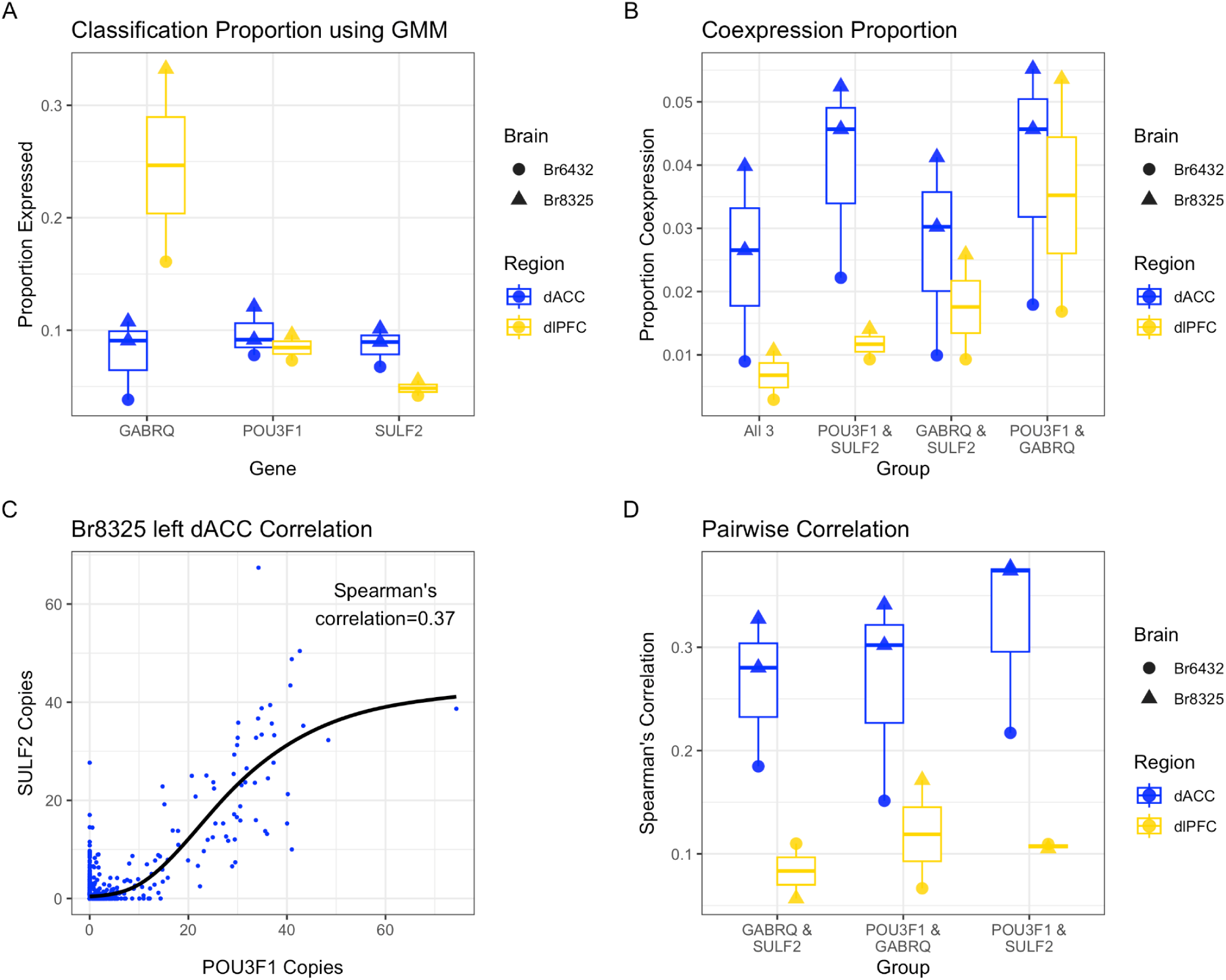
Analysis of RNAScope copy counts for VEN marker genes in dACC and dlPFC Layer 5 SRT data. (**A**) A Gaussian mixture model (GMM) with *k*=2 was used to classify the cells from each gene and sample as either expressing or not expressing the gene of interest. Boxplots display the proportion of cells (*y*-axis) from Layer 5 that were called as expressing each gene (*GABRQ*, *POU3F1*, and *SULF2*) (*x*-axis). Color indicates the region of the sample, either dACC or dlPFC. Shape indicates the brain donor of the sample, either Br6432 or Br8325. Note that there are two dACC samples from Br8325. (**B**) Boxplots display the proportion of cells (*y*-axis) from Layer 5 that were called as expressing two or three genes (*x*-axis). Color and shape same as (**A**). (**C**) Scatterplot displays the relationship between *POU3F1* copy count (*x*-axis) and *SULF2* copy count (*y*-axis) in dACC sample Br8325. A black loess smoothed line is fit to the data, and the Spearman’s correlation is displayed in the upper right corner. (**D**) Boxplots display Spearman’s correlation values (*y*-axis) for each pair of genes (*x*-axis). Color and shape same as (**A**).

**Supplementary Fig. 30.**
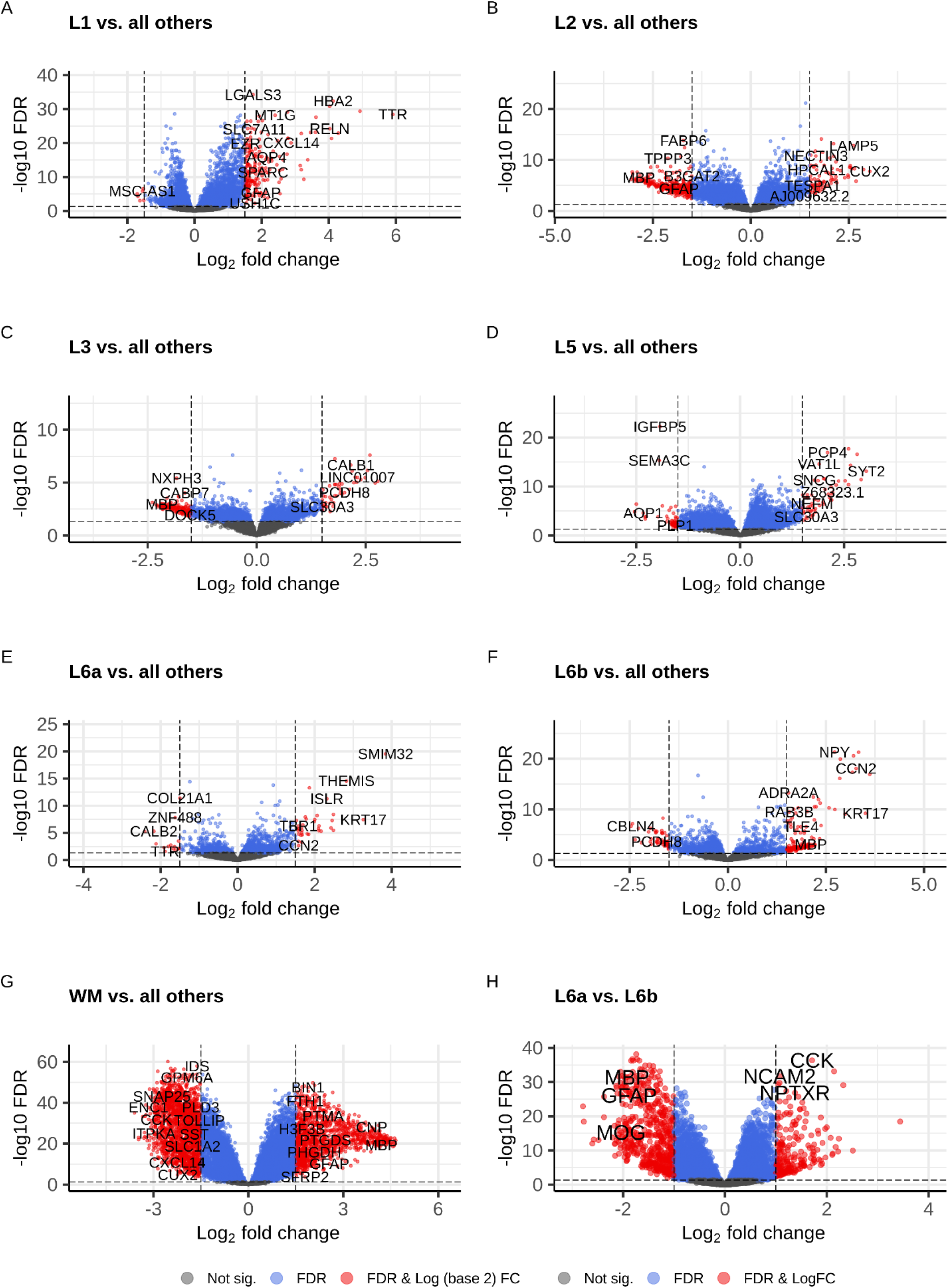
Differential expression (DE) analysis of pseudobulked dACC SRT data. **(A-G)** Each `EnhancedVolcano` plot shows the DE results for the enrichment model pseudobulked test for one dACC SRT spatial domain compared to all other dACC SRT spatial domains. Each point is a gene with its log fold-change (logFC) (*x*-axis) and statistical significance (*y*-axis). Statistical significance is measured with negative log-transformation of FDR-adjusted *p*-values. Color indicates categorization of each gene; red represents statistically significant with FDR < 0.05 and absolute value of logFC > 1, blue represents statistically significant with FDR < 0.05 only, and grey represents not statistically significant with FDR >= 0.05. **(H)** Similar to **(A-G)**, but the volcano plot shows DE results for the pairwise model pseudobulked test for dACC SRT spatial domain L6a compared to dACC SRT spatial domain L6b.

**Supplementary Fig. 31.**
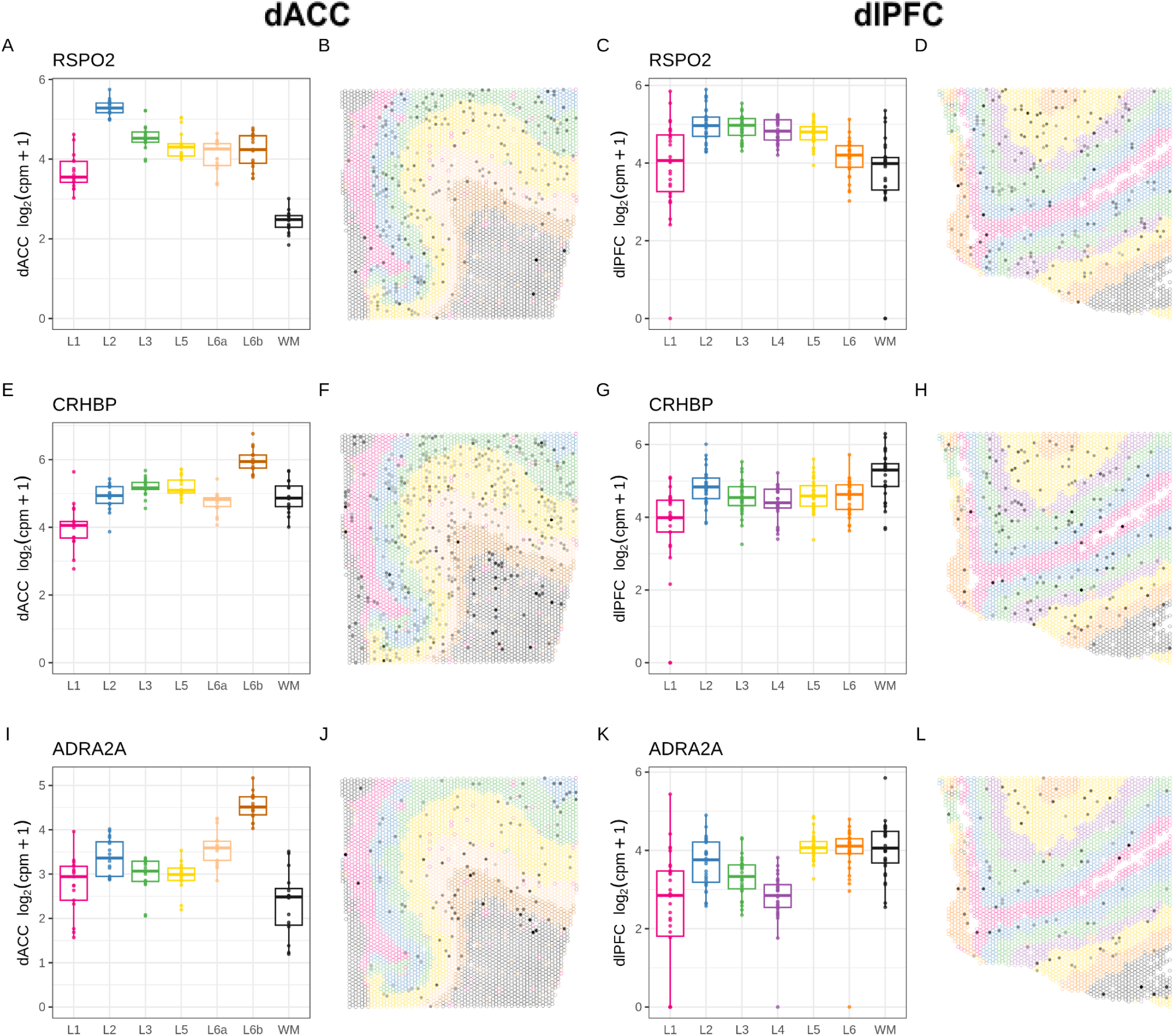
Spatial domain layer markers in dACC and dlPFC SRT data. Each row shows information for one gene, in order, *RSPO2, CRHBP, ADRA2A.* Each column displays a similar style of plot. First column (**A, E, I**): The *y*-axis displays log_2_(counts per million + 1) expression (computed manually) for each spatial domain (*x*-axis) in the pseudobulked dACC SRT data. Color represents the spatial domain. Second column (**B, F, J**): `escheR` spot plot of dACC Visium capture area from donor Br6432 (sample ID: V12N28-331_B1) with spots colored by the dACC spatial domains. Fill represents log_2_-normalized expression per spot. Third column (**C, G, K**): The *y*-axis displays log_2_(counts per million + 1) (computed manually) expression for each spatial domain (*x*-axis) in the pseudobulked dlPFC SRT data. Color represents the spatial domain. Fourth column (**D, H, L**): Spot plot of dlPFC Visium capture area from donor Br6432 (sample ID: Br6432_ant) with spots colored by the dlPFC spatial domains. Fill represents log_2_-normalized expression per spot.

**Supplementary Fig. 32.**
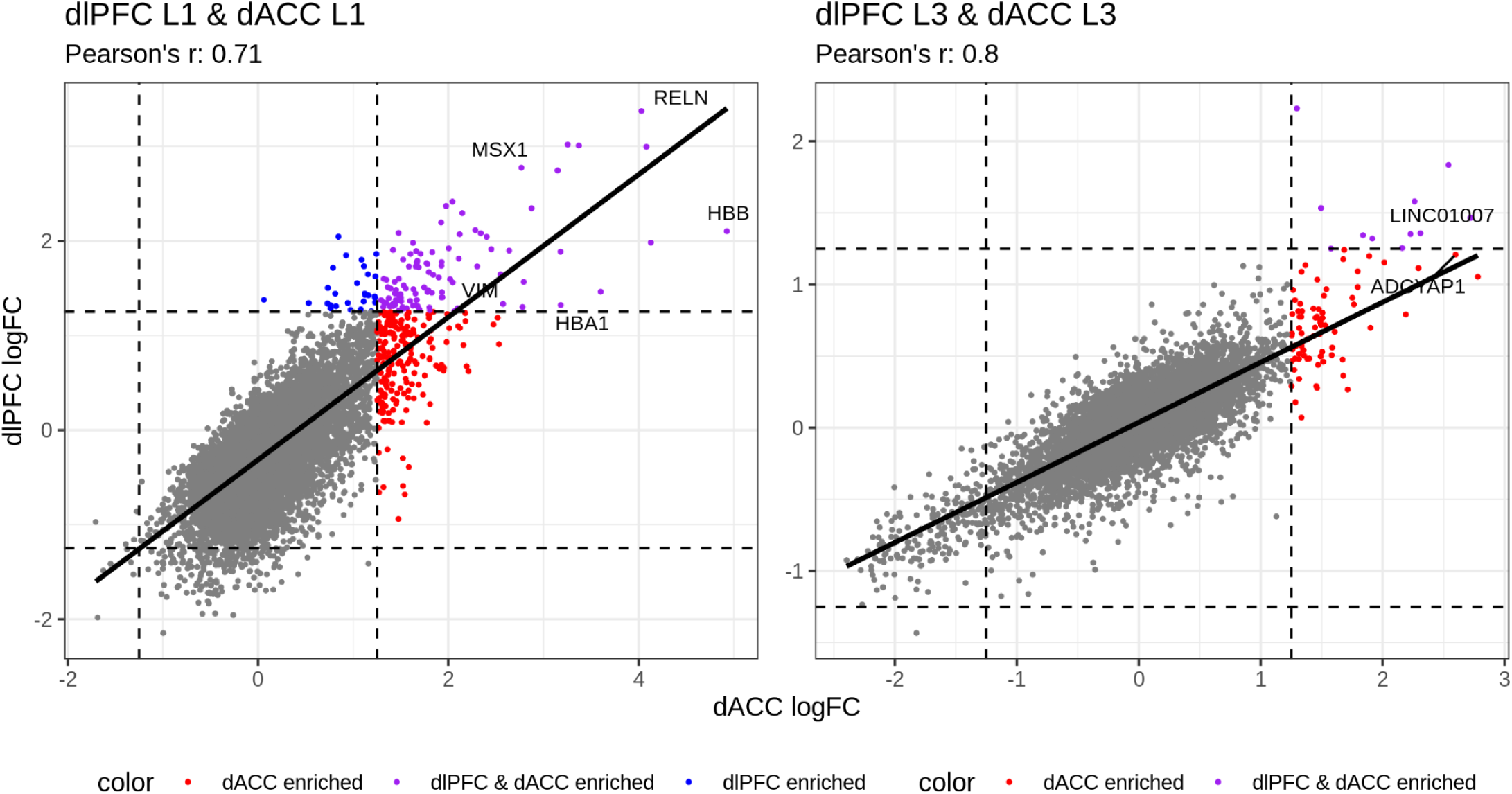
Scatterplots comparing pseudobulked differential expression (DE) results per gene between dACC and dlPFC spatial domains. Each scatterplot shows a comparison of the log fold-change (logFC) between one spatial domain in dACC and dlPFC (Huuki-Myers et al. 2024) SRT spatial domains. Each point is a gene. The *x*-axis of the first plot shows the logFC from enrichment model pseudobulked DE testing comparing Layer 1 to all other spatial domains for dACC, computing using `registration_wrapper()` (Pardo et al. 2022). The *y*-axis of the first plot shows the logFC from enrichment model pseudobulked DE testing comparing Layer 1 to all other spatial domains for dlPFC. The second plot compares L3 in the dACC to L3 in the dlPFC. The colors highlight genes that are classified as either dACC enriched (red), dlPFC enriched (blue), or both (purple), where the threshold is a logFC greater than 1.25. The number of genes in each category is as follows: L1) 197 dACC enriched & 28 dlPFC enriched & 101 dlPFC and dACC enriched and L3) 75 dACC enriched & 11 dlPFC and dACC enriched.

**Supplementary Fig. 33.**
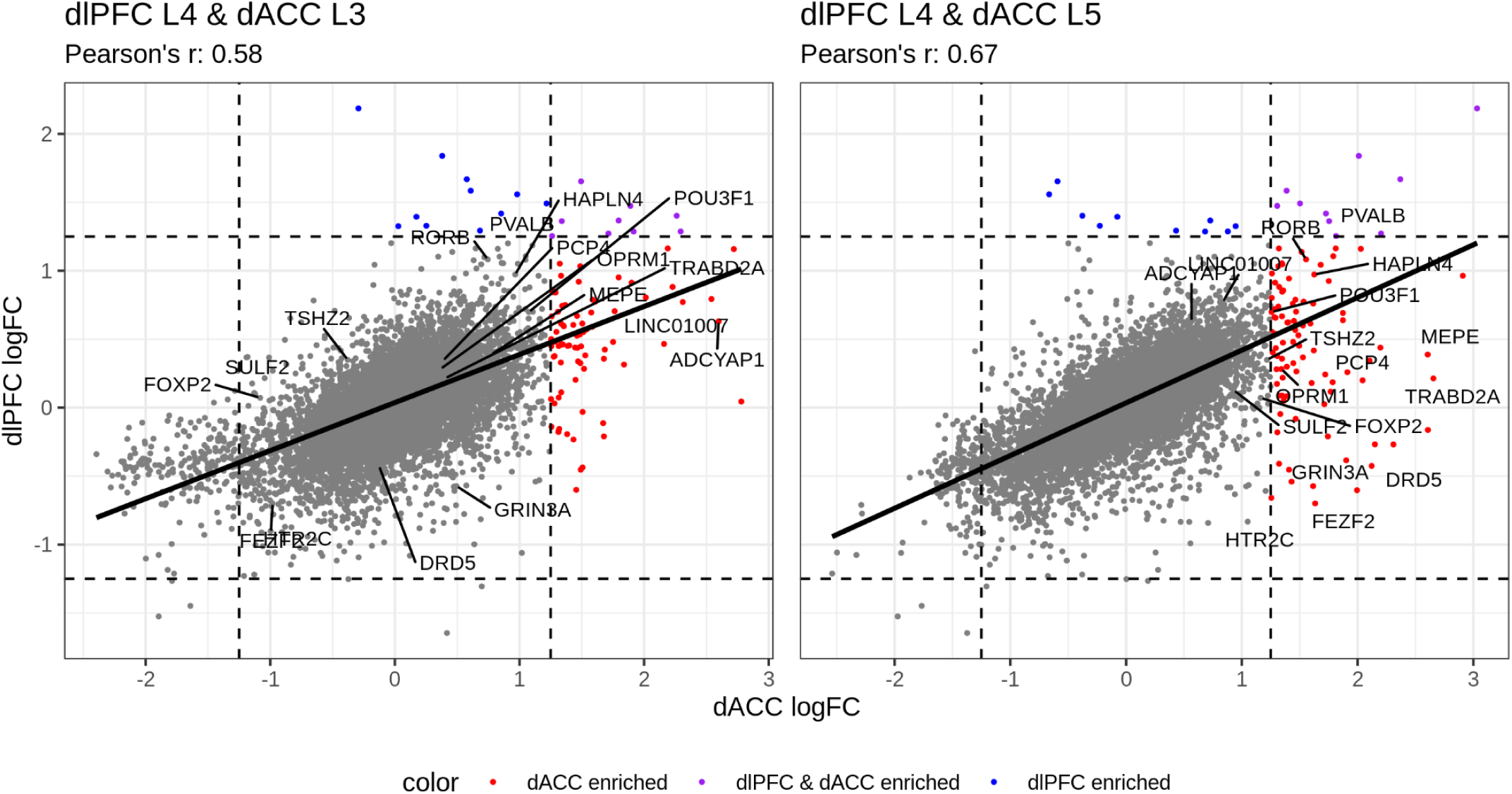
Scatterplots comparing pseudobulked differential expression (DE) results per gene between dACC and dlPFC spatial domains. Each scatterplot shows a comparison of the log fold-change (logFC) between one spatial domain in dACC and dlPFC (Huuki-Myers et al. 2024) SRT spatial domains. Each point is a gene. The *x*-axis of the first plot shows the logFC from enrichment model pseudobulked DE testing comparing Layer 3 to all other spatial domains for dACC, computing using `registration_wrapper()` (Pardo et al. 2022). The *y*-axis of the first plot shows the logFC from enrichment model pseudobulked DE testing comparing Layer 4 to all other spatial domains for dlPFC. The second plot compares L5 in the dACC to L4 in the dlPFC. The colors highlight genes that are classified as either dACC enriched (red), dlPFC enriched (blue), or both (purple), where the threshold is a logFC greater than 1.25. The number of genes in each category is as follows: L4 & L3) 77 dACC enriched & 11 dlPFC enriched & 9 dlPFC and dACC enriched and L4 & L5) 89 dACC enriched & 10 dlPFC enriched & 10 dlPFC and dACC enriched.

## Supplementary Tables

**Supplementary Table 1. Donor demographics.** Demographics for the 10 neurotypical control donors including age, sex, diagnosis, postmortem interval (PMI), screening RNA integrity number (RIN) in prefrontal cortex (PFC), each assay performed, and the number of replicates included (Visium H&E, Visium-SPG, snRNA-seq).

**Supplementary Table 2. nnSVG gene ranks.** As described in **Methods**, table contains results from nnSVG analysis that were used to identify spatially variable genes (SVGs) in the SRT data. Columns include ENSEMBL ID, gene name, overall gene rank, mean gene rank across all capture areas, and the number of capture areas for which each gene was highly ranked (within top 1000 most variable genes).

**Supplementary Table 3. Top 30 marker genes for snRNA-seq cell types.** For each gene, a linear mixed-effects model was fit with counts pseudobulked across spots within a cell type to identify differences in expression enriched in one cell type compared to all other cell types using Student’s *t*-test statistics.

**Supplementary Table 4. Top 10 marker genes for snRNA-seq NMF patterns.** For each NMF pattern, the top 10 genes contributing to the loadings matrix were selected as marker genes.

**Supplementary Table 5. Top 50 marker genes for SRT spatial domains.** For each gene, a linear mixed-effects model was fit with counts pseudobulked across spots within a spatial domain to identify differences in expression enriched in one domain compared to all other domains using Student’s *t*-test statistics.

**Supplementary Table 6. Top 50 pairwise markers comparing L6a and L6b SRT spatial domains.** For each gene, a linear mixed-effects model was fit with counts pseudobulked across spots within a spatial domain to identify differences in expression enriched in spatial domain L6a compared to spatial domain L6b using Student’s *t*-test statistics.

## Bibliography

1000 Genomes Project Consortium, Auton A, Brooks LD, Durbin RM, Garrison EP, Kang HM, et al. A global reference for human genetic variation. Nature. 2015 Oct 1;526(7571):68–74.

Allman JM, Hakeem A, Erwin JM, Nimchinsky E, Hof P. The anterior cingulate cortex. The evolution of an interface between emotion and cognition. Ann N Y Acad Sci. 2001 May;935:107–17.

Allman JM, Tetreault NA, Hakeem AY, Manaye KF, Semendeferi K, Erwin JM, et al. The von Economo neurons in the frontoinsular and anterior cingulate cortex. Ann N Y Acad Sci. 2011 Apr;1225:59–71.

Amezquita RA, Lun ATL, Becht E, Carey VJ, Carpp LN, Geistlinger L, et al. Orchestrating single-cell analysis with Bioconductor. Nat Methods. 2020 Feb;17(2):137–45.

Atta L, Clifton K, Anant M, Aihara G, Fan J. Gene count normalization in single-cell imaging-based spatially resolved transcriptomics. Genome Biol. 2024 Jun 12;25(1):153.

Bakken TE, Jorstad NL, Hu Q, Lake BB, Tian W, Kalmbach BE, et al. Comparative cellular analysis of motor cortex in human, marmoset and mouse. Nature. 2021 Oct 6;598(7879):111–9.

Batiuk MY, Tyler T, Dragicevic K, Mei S, Rydbirk R, Petukhov V, et al. Upper cortical layer-driven network impairment in schizophrenia. Sci Adv. 2022 Oct 14;8(41):eabn8367.

Becker LJ, Fillinger C, Waegaert R, Journée SH, Hener P, Ayazgok B, et al. The basolateral amygdala-anterior cingulate pathway contributes to depression-like behaviors and comorbidity with chronic pain behaviors in male mice. Nat Commun. 2023 Apr 17;14(1):2198.

Bhattacherjee A, Zhang C, Watson BR, Djekidel MN, Moffitt JR, Zhang Y. Spatial transcriptomics reveals the distinct organization of mouse prefrontal cortex and neuronal subtypes regulating chronic pain. Nat Neurosci. 2023 Nov;26(11):1880–93.

Blighe K, Rana S, Turkes E, Ostendorf B, Grioni A, Lewis M. EnhancedVolcano: Publication-ready volcano plots with enhanced colouring and labeling. R package; 2018.

Brüne M, Schöbel A, Karau R, Benali A, Faustmann PM, Juckel G, et al. Von Economo neuron density in the anterior cingulate cortex is reduced in early onset schizophrenia. Acta Neuropathol. 2010 Jun;119(6):771–8.

Brüne M, Schöbel A, Karau R, Faustmann PM, Dermietzel R, Juckel G, et al. Neuroanatomical correlates of suicide in psychosis: the possible role of von Economo neurons. PLoS ONE. 2011 Jun 22;6(6):e20936.

Bryden DW, Johnson EE, Tobia SC, Kashtelyan V, Roesch MR. Attention for learning signals in anterior cingulate cortex. J Neurosci. 2011 Dec 14;31(50):18266–74.

Bush G, Luu P, Posner MI. Cognitive and emotional influences in anterior cingulate cortex. Trends Cogn Sci (Regul Ed). 2000 Jun;4(6):215–22.

Bush G, Vogt BA, Holmes J, Dale AM, Greve D, Jenike MA, et al. Dorsal anterior cingulate cortex: a role in reward-based decision making. Proc Natl Acad Sci USA. 2002 Jan 8;99(1):523–8.

Carter CS, Braver TS, Barch DM, Botvinick MM, Noll D, Cohen JD. Anterior cingulate cortex, error detection, and the online monitoring of performance. Science. 1998 May 1;280(5364):747–9.

Chen Y, Chen L, Lun ATL, Baldoni PL, Smyth GK. edgeR v4: powerful differential analysis of sequencing data with expanded functionality and improved support for small counts and larger datasets. Nucleic Acids Res. 2025 Jan 11;53(2).

DeBruine ZJ, Melcher K, Triche TJ. Fast and robust non-negative matrix factorization for single-cell experiments. BioRxiv. 2021 Sep 1;

DeBruine Z. Rcpp Machine Learning: Fast robust NMF, divisive clustering, and more [Internet]. 2022a [cited 2025 Mar 20]. Available from: https://github.com/zdebruine/RcppML

DeBruine Z. Single-cell analysis with non-negative matrix factorization [Internet]. 2022b [cited 2025 Mar 20]. Available from: https://github.com/zdebruine/singlet

Devinsky O, Morrell MJ, Vogt BA. Contributions of anterior cingulate cortex to behaviour. Brain. 1995 Feb;118 ( Pt 1):279–306.

Dyjack N, Baker DN, Braverman V, Langmead B, Hicks SC. A scalable and unbiased discordance metric with H. Biostatistics. 2023 Dec 15;25(1):188–202.

Emani PS, Liu JJ, Clarke D, Jensen M, Warrell J, Gupta C, et al. Single-cell genomics and regulatory networks for 388 human brains. Science. 2024 May 24;384(6698):eadi5199.

Etkin A, Egner T, Kalisch R. Emotional processing in anterior cingulate and medial prefrontal cortex. Trends Cogn Sci (Regul Ed). 2011 Feb;15(2):85–93.

Germain P-L, Lun A, Garcia Meixide C, Macnair W, Robinson MD. Doublet identification in single-cell sequencing data using scDblFinder. F1000Res. 2021 Sep 28;10:979.

Griffiths JA, Richard AC, Bach K, Lun ATL, Marioni JC. Detection and removal of barcode swapping in single-cell RNA-seq data. Nat Commun. 2018 Jul 10;9(1):2667.

Guo B, Huuki-Myers LA, Grant-Peters M, Collado-Torres L, Hicks SC. escheR: unified multi-dimensional visualizations with Gestalt principles. Bioinformatics Advances. 2023 Dec 6;3(1):vbad179.

Gu Z, Eils R, Schlesner M. Complex heatmaps reveal patterns and correlations in multidimensional genomic data. Bioinformatics. 2016 Sep 15;32(18):2847–9.

Hansen DV, Hanson JE, Sheng M. Microglia in Alzheimer’s disease. J Cell Biol. 2018 Feb 5;217(2):459–72.

Hao Y, Hao S, Andersen-Nissen E, Mauck WM, Zheng S, Butler A, et al. Integrated analysis of multimodal single-cell data. Cell. 2021 Jun 24;184(13):3573–3587.

Hao Y, Stuart T, Kowalski MH, Choudhary S, Hoffman P, Hartman A, et al. Dictionary learning for integrative, multimodal and scalable single-cell analysis. Nat Biotechnol. 2024 Feb;42(2):293–304.

Hayden BY, Heilbronner SR, Pearson JM, Platt ML. Surprise signals in anterior cingulate cortex: neuronal encoding of unsigned reward prediction errors driving adjustment in behavior. J Neurosci. 2011 Mar 16;31(11):4178–87.

Heilbronner SR, Hayden BY. Dorsal Anterior Cingulate Cortex: A Bottom-Up View. Annu Rev Neurosci. 2016 Jul 8;39:149–70.

van Heukelum S, Mars RB, Guthrie M, Buitelaar JK, Beckmann CF, Tiesinga PHE, et al. Where is Cingulate Cortex? A Cross-Species View. Trends Neurosci. 2020 May;43(5):285–99.

Hodge RD, Miller JA, Novotny M, Kalmbach BE, Ting JT, Bakken TE, et al. Transcriptomic evidence that von Economo neurons are regionally specialized extratelencephalic-projecting excitatory neurons. Nat Commun. 2020 Mar 3;11(1):1172.

Hoffman GE, Roussos P. Dream: powerful differential expression analysis for repeated measures designs. Bioinformatics. 2021 Apr 19;37(2):192–201.

Hoffman GE, Schadt EE. variancePartition: interpreting drivers of variation in complex gene expression studies. BMC Bioinformatics. 2016 Nov 25;17(1):483.

Huuki-Myers LA, Spangler A, Eagles NJ, Montgomery KD, Kwon SH, Guo B, et al. A data-driven single-cell and spatial transcriptomic map of the human prefrontal cortex. Science. 2024 May 24;384(6698):eadh1938.

International HapMap 3 Consortium, Altshuler DM, Gibbs RA, Peltonen L, Altshuler DM, Gibbs RA, et al. Integrating common and rare genetic variation in diverse human populations. Nature. 2010 Sep 2;467(7311):52–8.

Jaffe AE, Tao R, Page SC, Maynard KR, Pattie EA, Nguyen CV, et al. Decoding Shared Versus Divergent Transcriptomic Signatures Across Cortico-Amygdala Circuitry in PTSD and Depressive Disorders. Am J Psychiatry. 2022 Sep;179(9):673–86.

Kim B, Kim D, Schulmann A, Patel Y, Caban-Rivera C, Kim P, et al. Cellular Diversity in Human Subgenual Anterior Cingulate and Dorsolateral Prefrontal Cortex by Single-Nucleus RNA-Sequencing. J Neurosci. 2023 May 10;43(19):3582–97.

Korsunsky I, Millard N, Fan J, Slowikowski K, Zhang F, Wei K, et al. Fast, sensitive and accurate integration of single-cell data with Harmony. Nat Methods. 2019 Dec;16(12):1289–96.

Law CW, Chen Y, Shi W, Smyth GK. voom: Precision weights unlock linear model analysis tools for RNA-seq read counts. Genome Biol. 2014 Feb 3;15(2):R29.

Lieberman MD, Eisenberger NI. The dorsal anterior cingulate cortex is selective for pain: Results from large-scale reverse inference. Proc Natl Acad Sci USA. 2015 Dec 8;112(49):15250–5.

Lipska BK, Deep-Soboslay A, Weickert CS, Hyde TM, Martin CE, Herman MM, et al. Critical factors in gene expression in postmortem human brain: Focus on studies in schizophrenia. Biol Psychiatry. 2006 Sep 15;60(6):650–8.

Liu W, Liao X, Luo Z, Yang Y, Lau MC, Jiao Y, et al. Probabilistic embedding, clustering, and alignment for integrating spatial transcriptomics data with PRECAST. Nat Commun. 2023 Jan 18;14(1):296.

Lui JH, Nguyen ND, Grutzner SM, Darmanis S, Peixoto D, Wagner MJ, et al. Differential encoding in prefrontal cortex projection neuron classes across cognitive tasks. Cell. 2021 Jan 21;184(2):489–506.e26.

Lun ATL, Riesenfeld S, Andrews T, Dao TP, Gomes T, participants in the 1st Human Cell Atlas Jamboree, et al. EmptyDrops: distinguishing cells from empty droplets in droplet-based single-cell RNA sequencing data. Genome Biol. 2019 Mar 22;20(1):63.

Lun A. Bioconductor - bluster [Internet]. Bioconductor - bluster. 2024 [cited 2025 Mar 30]. Available from: https://www.bioconductor.org/packages/release/bioc/html/bluster.html

Luscher B, Shen Q, Sahir N. The GABAergic deficit hypothesis of major depressive disorder. Mol Psychiatry. 2011 Apr;16(4):383–406.

Mai JK, Majtanik M, George Paxinos AO (BA MA PhD DSc) FASSA FAA. Atlas of the Human Brain. 4th ed. Academic Press; 2015. p. 456.

Maynard KR, Collado-Torres L, Weber LM, Uytingco C, Barry BK, Williams SR, et al. Transcriptome-scale spatial gene expression in the human dorsolateral prefrontal cortex. Nat Neurosci. 2021;24(3):425–36.

Maynard KR, Tippani M, Takahashi Y, Phan BN, Hyde TM, Jaffe AE, et al. dotdotdot: an automated approach to quantify multiplex single molecule fluorescent in situ hybridization (smFISH) images in complex tissues. Nucleic Acids Res. 2020 May 8;48(11):e66.

McCarthy DJ, Campbell KR, Lun ATL, Wills QF. Scater: pre-processing, quality control, normalization and visualization of single-cell RNA-seq data in R. Bioinformatics. 2017 Apr 15;33(8):1179–86.

Milad MR, Quirk GJ, Pitman RK, Orr SP, Fischl B, Rauch SL. A role for the human dorsal anterior cingulate cortex in fear expression. Biol Psychiatry. 2007 Nov 15;62(10):1191–4.

Mulvey B, Wang Y, Divecha HR, Bach SV, Montgomery KD, Cinquemani S, et al. Spatially-resolved molecular sex differences at single cell resolution in the adult human hypothalamus. BioRxiv. 2024 Dec 9;

Nimchinsky EA, Gilissen E, Allman JM, Perl DP, Erwin JM, Hof PR. A neuronal morphologic type unique to humans and great apes. Proc Natl Acad Sci USA. 1999 Apr 27;96(9):5268–73.

Pardo B, Spangler A, Weber LM, Page SC, Hicks SC, Jaffe AE, et al. spatialLIBD: an R/Bioconductor package to visualize spatially-resolved transcriptomics data. BMC Genomics. 2022 Jun 10;23(1):434.

Peng H, Xie P, Liu L, Kuang X, Wang Y, Qu L, et al. Morphological diversity of single neurons in molecularly defined cell types. Nature. 2021 Oct 6;598(7879):174–81.

Petrides M, Pandya DN. Comparative cytoarchitectonic analysis of the human and the macaque ventrolateral prefrontal cortex and corticocortical connection patterns in the monkey. Eur J Neurosci. 2002 Jul;16(2):291–310.

Rainville P, Duncan GH, Price DD, Carrier B, Bushnell MC. Pain affect encoded in human anterior cingulate but not somatosensory cortex. Science. 1997 Aug 15;277(5328):968–71.

Righelli D, Weber LM, Crowell HL, Pardo B, Collado-Torres L, Ghazanfar S, et al. SpatialExperiment: infrastructure for spatially-resolved transcriptomics data in R using Bioconductor. Bioinformatics. 2022 May 26;38(11):3128–31.

Robinson MD, McCarthy DJ, Smyth GK. edgeR: a Bioconductor package for differential expression analysis of digital gene expression data. Bioinformatics. 2010 Jan 1;26(1):139–40.

Rue-Albrecht K, Marini F, Soneson C, Lun ATL. iSEE: Interactive SummarizedExperiment Explorer. [version 1; peer review: 3 approved]. F1000Res. 2018 Jun 14;7:741.

Santos M, Uppal N, Butti C, Wicinski B, Schmeidler J, Giannakopoulos P, et al. Von Economo neurons in autism: a stereologic study of the frontoinsular cortex in children. Brain Res. 2011 Mar 22;1380:206–17.

Scrucca L, Fraley C, Murphy TB, Adrian E. R. Model-Based Clustering, Classification, and Density Estimation Using mclust in R. Boca Raton: Chapman and Hall/CRC; 2023.

Seidman LJ, Valera EM, Makris N, Monuteaux MC, Boriel DL, Kelkar K, et al. Dorsolateral prefrontal and anterior cingulate cortex volumetric abnormalities in adults with attention-deficit/hyperactivity disorder identified by magnetic resonance imaging. Biol Psychiatry. 2006 Nov 15;60(10):1071–80.

Seo H, Lee D. Temporal filtering of reward signals in the dorsal anterior cingulate cortex during a mixed-strategy game. J Neurosci. 2007 Aug 1;27(31):8366–77.

Shackman AJ, Salomons TV, Slagter HA, Fox AS, Winter JJ, Davidson RJ. The integration of negative affect, pain and cognitive control in the cingulate cortex. Nat Rev Neurosci. 2011 Mar;12(3):154–67.

Skene NG, Bryois J, Bakken TE, Breen G, Crowley JJ, Gaspar HA, et al. Genetic identification of brain cell types underlying schizophrenia. Nat Genet. 2018 Jun;50(6):825–33.

Spocter MA, Hopkins WD, Barks SK, Bianchi S, Hehmeyer AE, Anderson SM, et al. Neuropil distribution in the cerebral cortex differs between humans and chimpanzees. J Comp Neurol. 2012 Sep 1;520(13):2917–29.

Sriworarat C, Nguyen A, Eagles NJ, Collado-Torres L, Martinowich K, Maynard KR, et al. Performant web-based interactive visualization tool for spatially-resolved transcriptomics experiments. Biol Imaging. 2023 Jul 12;3:e15.

Stein-O’Brien GL, Clark BS, Sherman T, Zibetti C, Hu Q, Sealfon R, et al. Decomposing Cell Identity for Transfer Learning across Cellular Measurements, Platforms, Tissues, and Species. Cell Syst. 2019 May 22;8(5):395–411.e8.

Stevens FL, Hurley RA, Taber KH. Anterior cingulate cortex: unique role in cognition and emotion. J Neuropsychiatry Clin Neurosci. 2011;23(2):121–5.

Thompson JR, Nelson ED, Tippani M, Ramnauth AD, Divecha HR, Miller RA, et al. An integrated single-nucleus and spatial transcriptomics atlas reveals the molecular landscape of the human hippocampus. BioRxiv. 2024 Apr 28;

Tippani M, Divecha HR, Catallini JL, Kwon SH, Weber LM, Spangler A, et al. VistoSeg: Processing utilities for high-resolution images for spatially resolved transcriptomics data. Biol Imaging. 2023 Nov 13;3:e23.

Townes FW, Hicks SC, Aryee MJ, Irizarry RA. Feature selection and dimension reduction for single-cell RNA-Seq based on a multinomial model. Genome Biol. 2019 Dec 23;20(1):295.

Vogt BA, Nimchinsky EA, Vogt LJ, Hof PR. Human cingulate cortex: surface features, flat maps, and cytoarchitecture. J Comp Neurol. 1995 Aug 28;359(3):490–506.

Vogt BA, editor. Cingulate neurobiology and disease. Oxford University PressOxford; 2009.

Wamsley B, Bicks L, Cheng Y, Kawaguchi R, Quintero D, Margolis M, et al. Molecular cascades and cell type-specific signatures in ASD revealed by single-cell genomics. Science. 2024 May 24;384(6698):eadh2602.

Watson KK, Jones TK, Allman JM. Dendritic architecture of the von Economo neurons. Neuroscience. 2006 Sep 1;141(3):1107–12.

Weber LM, Saha A, Datta A, Hansen KD, Hicks SC. nnSVG for the scalable identification of spatially variable genes using nearest-neighbor Gaussian processes. Nat Commun. 2023 Jul 10;14(1):4059.

Weissman DH, Gopalakrishnan A, Hazlett CJ, Woldorff MG. Dorsal anterior cingulate cortex resolves conflict from distracting stimuli by boosting attention toward relevant events. Cereb Cortex. 2005 Feb;15(2):229–37.

Wickham H. ggplot2: Elegant Graphics for Data Analysis (Use R!). 2nd ed. Cham: Springer; 2016. p. 276.

Yang L, Yang Y, Yuan J, Sun Y, Dai J, Su B. Transcriptomic Landscape of von Economo Neurons in Human Anterior Cingulate Cortex Revealed by Microdissected-Cell RNA Sequencing. Cereb Cortex. 2019 Feb 1;29(2):838–51.

Zhou J, Zhang Z, Wu M, Liu H, Pang Y, Bartlett A, et al. Brain-wide correspondence of neuronal epigenomics and distant projections. Nature. 2023 Dec 13;624(7991):355–65.

Zhu Y, Sousa AMM, Gao T, Skarica M, Li M, Santpere G, et al. Spatiotemporal transcriptomic divergence across human and macaque brain development. Science. 2018 Dec 14;362(6420). LieberInstitute/spatialdACC: preprint-v1.0.0. Zenodo. 2025;

Rue-Albrecht, K., Marini, F., Soneson, C., & Lun, A. T. L. (2018). ISEE: Interactive SummarizedExperiment Explorer. F1000Research, 7, 741. doi:10.12688/f1000research.14966.1

Shah, K., Tippani, M., Miller, R., Divecha, H., Nick-Eagles, & Totty, M. (2025). LieberInstitute/spatialdACC: preprint-v1.0.0. doi:10.5281/ZENODO.15830481

